# Single-cell X-chromosome inactivation analysis links biased chimerism to differential gene expression and epigenetic erosion

**DOI:** 10.1101/2024.08.29.610317

**Authors:** Robert H. Henning, Thomas M. Rust, Kasper Dijksterhuis, Bart J.L. Eggen, Victor Guryev

## Abstract

Female cells randomly inactivate one X-chromosome, resulting in cellular mosaicism. Determining the bias in X-chromosomal inactivation (XCI) at single cell level may be relevant for understanding diseases prevalent in females. Here, we introduce a computational method that determines XCI profiles at single-cell level using solely sc/snRNA-Seq data. XCI analysis of skin cells from hybrid mice validates our approach and reveals biased inactivation of X-chromosomes among cell types. In human lung and brain cells, XCI status can be determined in 33.8% and 23.6% of cells. Among the patients, cells with opposite inactivation patterns differently express members of specific gene families and pathways. Alzheimer’s disease patients show reversal of XCI in cortical microglia and regional increase in biallelic expression denoting epigenetic erosion. We provide a robust utility to explore the degree and impact of XCI in single cell expression data.

---

In placental mammals, random X-chromosome inactivation (XCI) in females occurs early in development to balance X-chromosomal dosage between sexes^1^. Consequently, XCI results in cellular mosaicism with silenced maternal or paternal X-chromosomes. Up to 25% of X-linked genes escape XCI in humans – about 15% constitutively and 10% variably among individuals^2^. Due to its randomness or selective growth advantage, XCI may result in a skewed distribution of cells with inactivated paternal or maternal X-chromosome, showing large variability between individuals and tissues^3^. Skewed XCI may profoundly affect the penetrance of X-linked disease genes in females, but also impact other diseases as most X-chromosome genes serve general cellular functions^4^. While firmly established, the effects of X-chromosome dosage and XCI escape on disease severity have mostly been documented in case studies. Advancement of our general understanding of the relationship between XCI and phenotype severity requires larger sample sizes and - given its large variation - a detailed insight of XCI skewing across different cell types in individuals.

Next-generation sequencing technologies for transcriptomics are increasingly characterizing individual cells through single-cell RNA sequencing (scRNA-seq) rather than using conventional bulk RNA-seq. Beyond exploration of variation in gene expression between different cell types, we reasoned that it allows investigation of XCI, XCI escape and reactivation of inactivated X on gene expression in females, revealing their relevance for disease penetrance and progression. Here, we developed XCISE (XCI calling from Single cell Expression data), a computational pipeline that identifies cells with different active parental X-chromosomes using solely scRNA-seq data (Fig. 1a). XCISE workflow calls X-linked heterozygous single nucleotide variants (SNVs) and clusters cells with overlapping expressed alleles to deliver the cell status (X_1_ active, X_2_ active, both active, unknown/low coverage) and expression status of each SNV (monoallelic or biallelic expression). We validated our approach by exploring scRNA-seq data of mouse F1 hybrids (for which X-chromosomal haplotypes are known) and studied the impact of XCI on cell-type specific gene expression and disease phenotypes in females using scRNA-seq data from subjects with/without lung disease and dementia. The pipeline effectively detects XCI-informative SNVs and identifies cell pools, both in single-cell and single-nuclei RNA-seq of mouse F1 hybrids and patient cohorts. We found expression of specific genes to depend on XCI status. In addition, Alzheimer’s disease patients show opposite bias in XCI in cortical microglia and increased epigenetic erosion compared to matched controls. Assessment of XCI status provides valuable insights into cell type-specific variations in gene expression and gene regulation observed in sc/snRNA-seq data from females.

**Figure 1.**
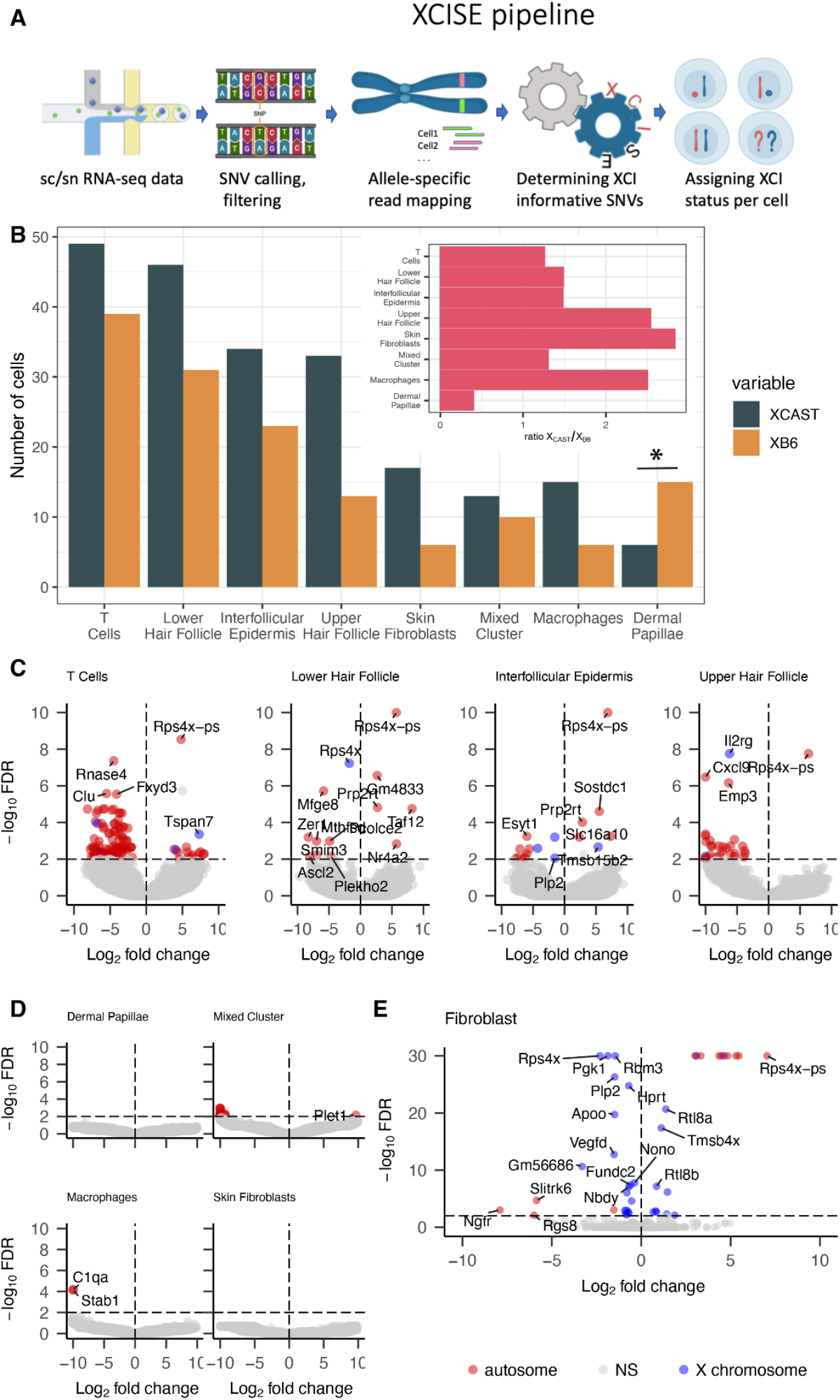
XCISE pipeline validated by analysis of single-cell RNA-seq from the F1 cross of CAST x B6 mice. **A**. Schematic representation of XCISE pipeline. **B**. Distribution of skin cells with active X_CAST_ and X_B6_ chromosomes in F1 of CASTxB6 mouse across the various cell types. Only cell types with total cell count > 10 are included. * denotes significance deviation from distribution in the rest of cell types (χ^2^-test) **C**,**D**. Volcano plots of expression differences in skin cells depending on XCI in the 8 most abundant cell types, with cells with active X_B6_ denoted up-regulated and those with active X_CAST_ downregulated. **E**. Volcano plot of expression differences in fibroblasts depending on XCI. DEGs at FDR < 0.01 are partially annotated and indicated by red dots for autosomal genes and blue dots for X chromosomal genes. FDR values for Low Hair Follicle and Interfollicular Epidermis (panel C) are clipped at 1×10^−10^ and for fibroblasts (panel E) at 1×10^−30^.

## Results

### Cell-specific X-chromosome inactivation can be deduced from scRNA-seq data

To validate the algorithm, we called X-chromosomal inactivation in females from a cross of two inbred mouse strains with known SNV alleles of parental strains (Fig. 1a). Hereto, we used the public single-cell RNA sequencing (scRNA-seq) dataset from skin of *Mus musculus castaneus* (CAST/EiJ) x *Mus musculus* C57BL/6J F1 hybrid (referred to as CAST x B6)^6^. The allele-specific expression patterns of 1555 informative SNVs largely correspond to activation of the X_CAST_ (1534 SNVs; Supplementary Table 1). The B6 alleles of the remaining 21 SNVs map to two X-chromosomal loci. Expectedly, the majority (n=19) maps to *Xist*, collectively representing all SNVs called for that gene, reflecting *Xist-*maintained silencing of the inactivated X-chromosome. The remaining 2 SNVs are located in *Pbdc1*, explained by haplotype-specific alternative splicing of this X chromosomal gene known to escape XCI in mice^7^ (Extended Data Fig. 1). Out of 384 cells, the majority have an active X_CAST_ (218 cells, 56.8%), while 148 cells (38.5%) have an active X_B6_ (Fig. 1b, Supplementary Table 1). Seventeen cells (4.4%) express both X-chromosomes, which is not due to presence of doublets (i.e. two cells in a well) as these cells show even significantly lower total reads than X_B6_ or X_CAST_ pools (Extended Data Fig. 2). The remaining cell has only a single read overlapping an XCI-informative SNV (Extended Data Fig. 2). Moreover, the estimated level of discordant alleles (proportion of reads overlapping XCI-informative SNVs with alleles not matching to the reconstructed XCI of that cell) is 1.3%, signifying that only a minimal proportion of haplotype-informative reads have alleles contradicting the invoked XCI. Next, we examined differences in XCI among cell types (Fig. 1b). In skin cells, there is a clear bias towards excess silencing of X_B6_ chromosomes (p=5×10^−5^; binomial test). Individual cell types do not show significant deviations from the global ratio, likely due to low number of cells. The notable exception is dermal papillae cells (DPCs), which have a significant excess of cells with active X_B6_ (p=0.004, χ^2^-test). A larger-scale study is needed to confirm this finding and explore whether the excess of active X_B6_ in DPCs relates to survival or proliferation differences possibly due to their developmental origin (mesenchymal rather than epithelial).

The XCISE algorithm was further validated using fibroblasts from an independent CAST x B6 F1 cross scRNA-seq dataset (E-MTAB-6385)^6^. Here, alleles at 1965 haplotype-informative SNVs result in 372 cells with either active X_CAST_ (n=273) or active X_B6_ (n=99). Only 2 fibroblasts showed expression from both X-chromosomes. Further, transcripts in cells with an active X_CAST_ have a B6 allele for 14 SNVs, which all (and only) map to the *Xist* gene (Supplementary Table 1).

The CAST x B6 F1 mouse hybrid has more informative SNVs (21.7×10^6^ per genome)^8,9^ than humans (∼3×10^6^ per genome)^10^. To simulate performance under lower abundant SNV conditions, we randomly removed 90% of SNVs and repeated the XCISE analysis. Results of all 5 repetitions of this analysis are highly similar to the full skin dataset with cell numbers for active X_CAST_ and X_B6_ ranging between 211-216 and 145-150, respectively. Similarly to the full dataset, only alleles of *Xists* and *Pbdc1* show reads with alleles expressed from the opposite genetic background, proving that XCISE retrieves the correct XCI pattern even in datasets with reduced heterozygosity levels.

Finally, as a negative control, we analyzed fibroblasts from a male F1 mouse^6^. Here, of the 678 X-linked SNVs, the algorithm identifies only 36 informative ones that could discriminate between cell pools. The majority of these SNVs (n=24) are in or close to (within 500 kb) the PAR (pseudoautosomal region), a shared homologous segment of X and Y chromosomes that recombines during meiosis, signifying that the algorithm discriminates between corresponding parts of X and Y chromosomes (Supplementary Table 1).

### Differential gene expression between mouse skin cells with active X_CAST_ and X_B6_

We then investigated the impact of XCI on differentially expressed genes (DEGs) between X_CAST_ and X_B6_ at FDR < 0.01 (Fig. 2a, Supplementary Table 2). There are 356 DEGs of which only 2.8% originate from the X-chromosome, although it harbors 4.3% of all protein coding genes (Gencode release M34^11^), indicating a strong trans effect of XCI on autosomal gene expression. Cell types differ substantially in their number of DEGs, even among those with similar and highest numbers of haplotyped cells (T cells, lower hair follicle and interfollicular epidermis) in which DEGs amount 130, 13 and 18, respectively. The two top DEGs, a gene-pseudogene pair of *Rps4x* and *Rps4x-ps* (Fig. 1c), is likely a false positive finding. *Rps4x-ps* is a chromosome 5 located polymorphic retrocopy present in B6 mice (but not in CAST)^12^, which shares a region identical to the CAST *Rps4x* source gene (around T-allele of rs579147927 SNV). It shows preferential mapping of *Rps4x* reads from the CAST haplotype in X_CAST_ cells, rendering gene and pseudogene seemingly differentially expressed in opposite directions. A remarkable finding is the highly skewed distribution of upregulated transcripts in T cells with active X_CAST_ *vs* active X_B6_ (102 *vs* 18 genes). The significance of this finding is unclear and subsequent gene ontology (GO) analysis of DEGs in T-cells, as in other cell types, does not show specific enrichment of functional terms. Mouse fibroblast has 42 DEGs (Fig. 1e) with a much higher fraction of X chromosomal genes compared to skin cells (30 genes = 71.4 %, although showing smaller fold changes). Interestingly, multiple genes show consistent differential expression across multiple cell types, of which the functional consequences are unclear.

**Figure 2.**
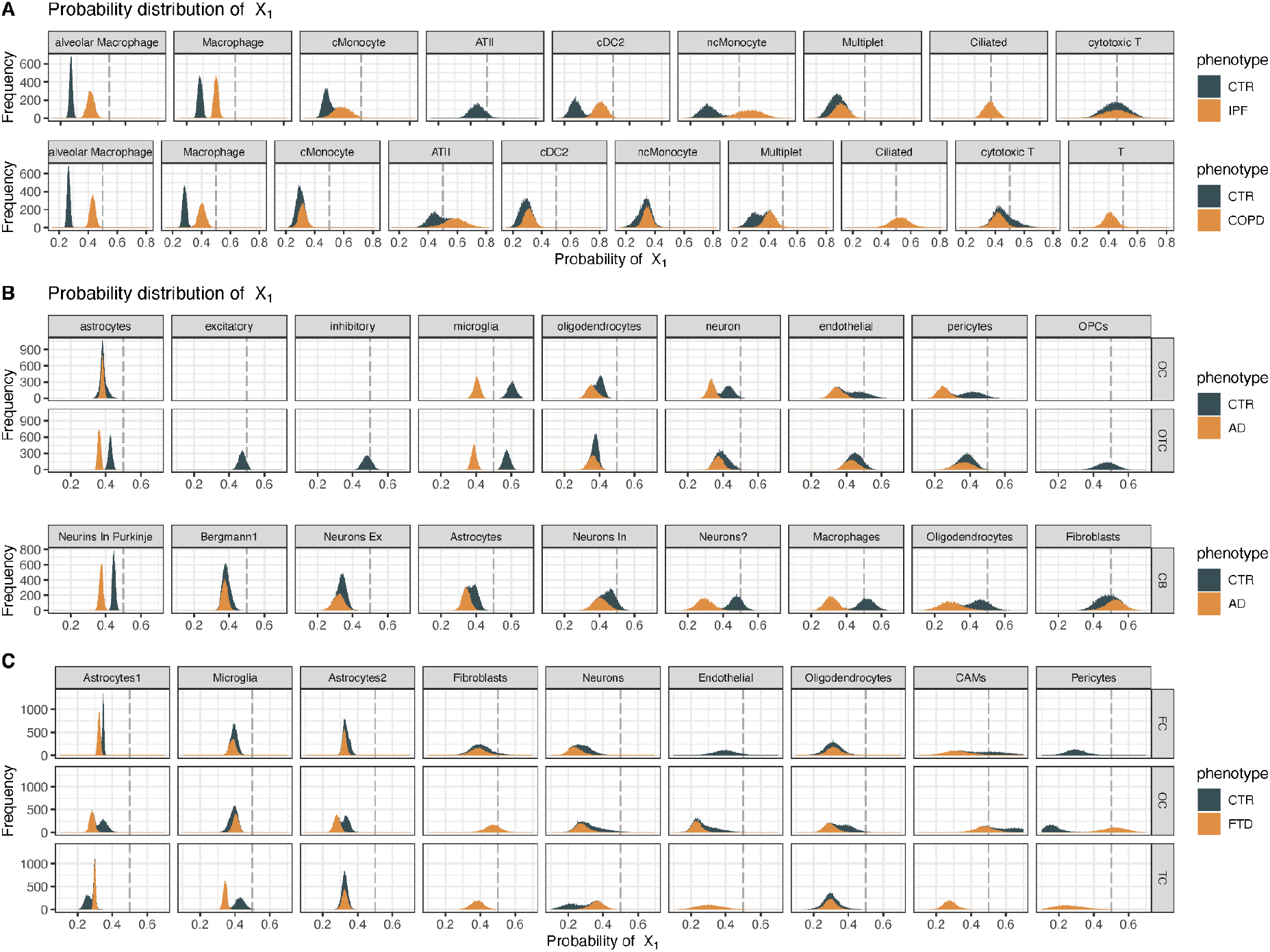
Posterior probability distribution of the fraction of X_1_ cells of the total of cells with known XCI status across cell types and brain regions and disease conditions. The cells with less frequently activated X-chromosome across all cells of each patient were designated to the X_1_ pool and the others to X_2_. For each cell type and condition, the number of cells of all subjects with 10 or more cells with known XCI pattern were included as prior data. Cell types are left to right arranged from most to least abundant. **A**. Analysis of a public scRNA-Seq dataset from controls (CTR) and pulmonary patients with interstitial pulmonary fibrosis (IPF) or chronic obstructive pulmonary disease (COPD). **B, C**. Analysis of a snRNA-Seq dataset from controls (CTR) and dementia patients with Alzheimer’s disease (AD) or fronto-temporal dementia (FTD) arranged by brain region sampled (OC = occipital cortex, OTC = occipito-temporal cortex, CB = cerebellum, FC = frontal cortex, TC = temporal cortex).

Fibroblasts show a high CAST/B6 ratio of active X in both mouse datasets (2.83 and 2.76), representing an obvious and significant (p=0.017; p=3·10^−20^, binomial test) bias towards an active X_CAST_ in 3/4 of the cells. While the general preference for X_CAST_ may result from higher expression of the B6 *Xist* allele, observed in CAST x B6 mice before^13^, the cell type differences (e.g. in DPCs) warrant further research into the impact of XCI on cell proliferation among cell types.

Collectively, these analyses demonstrate that scRNA-seq data analysis using XCISE defines XCI-informative SNVs and identifies cell-specific expression from the active X, of *Xist* from the inactivated X as well as allelic disbalances due to haplotype-specific alternative splicing events, enabling the exploration of XCI dependent gene expression in single cell RNA-Seq datasets.

### X-chromosome inactivation analysis in human single-cell RNA-seq

To demonstrate suitability of the XCISE algorithm to interrogate routinely collected human scRNA-seq data, we analyzed female controls and lung patients using a public dataset from 9 controls, 8 COPD and 5 IPF patients (Supplementary Tables 1,3)^14^. We designated cells with the most frequently inactivated X-chromosome in each subject as X_1_ and others as X_2_. Of the 59,213 cells sequenced, 20,032 (33.8 %) showed reliable XCI status, representing either X_1_ or X_2_ cell pools (98.2% of cells) or both X-chromosomes (1.8% of cells). Under the constraint that at least 3 subjects have 100 or more cells from a cell type sequenced, the silenced X is successfully determined in 48.5%, 54.5% and 37.2% of the cells in controls, COPD and IPF patients, respectively. In addition, under the same constraint, XCI status varies largely per cell type per patient ranging from 14.5% to 89.9% (see Extended Data Fig. 3).

We then asked whether X-chromosome inactivation differs related to disease or cell type. When XCI status is determined for 3 or more subjects in 10 or more cells per cell type, 40.7 ± 5.3 % of cells belong to the X_1_ pool, ranging from 31% to 52% across cell types. Visualizing differential X-chromosome inactivation for the most abundant cell types revealed marked differences between cell types and conditions (Fig. 2a). IPF patients lack sufficient alveolar type II pneumocytes. Controls lack ciliated cells when compared to IPF and COPD, and lack T cells compared to COPD. Notably, controls show an extreme X_1_ bias in macrophages and alveolar macrophages, which is less extreme in both IPF and COPD patients (Fig. 2a).

### X-chromosome inactivation analysis in human single-nuclei RNA-seq

Additionally, we analyzed single-nucleus RNA-seq of enriched cell populations in different brain regions in two cohorts of dementia patients and non-demented matched control subjects (Supplementary Tables 1,4). Patients were diagnosed with Alzheimer’s disease (AD cohort)^15^ and Frontotemporal dementia (FTD cohort)^16^. Similarly, cells with the “least active” X-chromosome across all cells of each subject are designated X_1_. Among 491,824 nuclei analyzed, 116,020 (23.6%) were assigned to either the X_1_ or X_2_ pool and 10,836 to X_BOTH_ (2.2%). XCI rate varies widely by cell type and brain region (XCI range: 1.9 - 77.5%; Extended Data Fig. 4), even with adequate numbers of cells sequenced (cells per patient per region: median = 8,386, range = 1,299-15,108). Expectedly, smaller cells (fewer UMIs, e.g. microglia) show lower XCI rate.

Next we examined differences in XCI across conditions, brain regions and cell types, provided XCI status is determined in 3 or more subjects and in 10 or more cells. Bayesian posterior distribution shows that the X_1_ pool comprises 37.7 ± 9.6 % of cells, ranging between 21.7 % and 46.2 % across patients and cell types (Fig. 2b,c). XCI in microglia of cortical regions in the AD cohort is intriguing (Fig. 2b), as controls show an opposite X-inactivation pattern in OC and OTC compared to other cell types (i.e. control subject’s microglia have >50% active X_1_), which is reversed in microglia of AD patients. In the temporal cortex (TC) of the FTD cohort, XCI in microglia is similar to that observed for AD patients.

To test the algorithm’s rigidity in snRNA-seq, we included a male control from the FTD cohort. Of the 315 SNVs mapped on the X-chromosome, 149 were XCI informative, mainly located in the PAR1 region (n=143). The low number of SNVs in male and their overall mapping to PAR1 signify that the XCISE algorithm, also in human snRNA-Seq, readily discriminates between female and male expression data.

### Differential gene expression between cells with different active X in human datasets

We next determined DEGs between X_1_ and X_2_ cells per cell type. In lung samples, the most prominent DEGs are present both in COPD and IPF patients (Fig. 3a, Supplementary Table 5). *RPL10P9*, a chromosome 5 encoded pseudogene, is XCI-dependently expressed in diverse cell types of mainly COPD patients. Among the most frequent DEGs are also *SCGB1A1, SCGB3A1* and *SCGB3A2*, encoded on chromosomes 11 and 5 and representing three Club cell markers and members of the secretoglobin family - ill-characterized cytokine-like proteins putatively implicated in inflammatory responses and tissue repair^17^. Because differential expression between X_1_ and X_2_ cells is biased towards COPD and IPF patients, this signifies that XCI may modulate lung disease via secretoglobins. In addition, the CC chemokine ligands (CCL) family, the surfactant protein genes (SFTP) and the metallothionein family (MTI) each have 3 members showing increased expression in X_2_ both in controls and patients.

**Figure 3.**
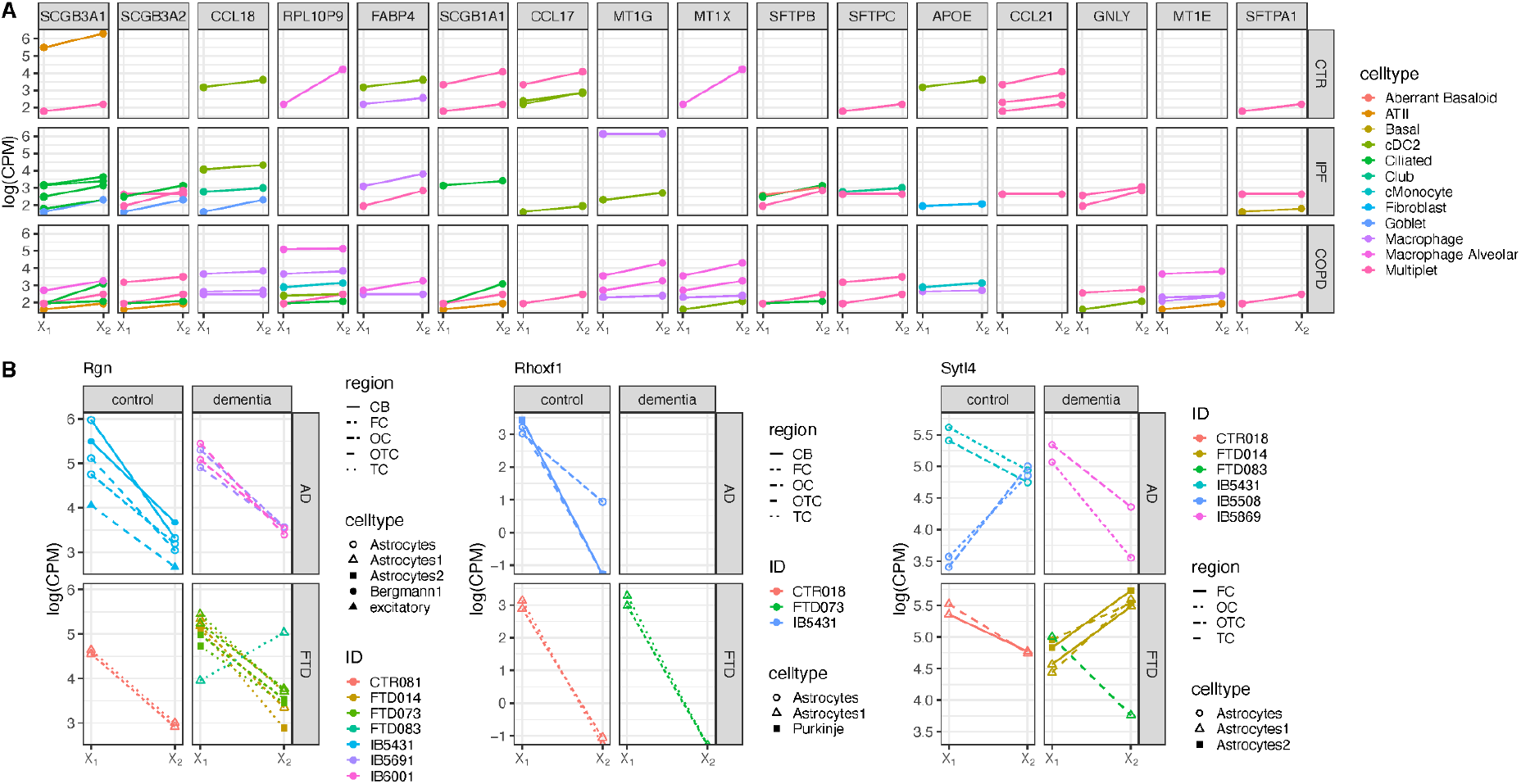
Differential gene expression between X_1_ and X_2_ cells. The cells with an active X-chromosome less frequently activated across all cells of each patient were designated as X_1_ and the others as X_2_. **A**. Analysis of a public scRNA-Seq dataset from controls (CTR) and pulmonary patients with interstitial pulmonary fibrosis (IPF) or chronic obstructive pulmonary disease (COPD). Panels are ordered from left to right by the total number of cells with differential expression. **B**. Analysis of a snRNA-Seq dataset from controls (CTR) and dementia patients with Alzheimer’s disease (AD) or fronto-temporal dementia (FTD) arranged by brain region sampled (OC = occipital cortex, OTC = occipito-temporal cortex, CB = cerebellum, FC = frontal cortex, TC = temporal cortex). Data are included if XCI calling was successful in >= 10 cells per cell type and FDR <0.01.

A similar analysis was performed in dementia cohorts (Fig. 3b, Supplementary Table 6). Only 3 DEGs are regulated both in controls and dementia patients. Notably, *Regucalcin (Rgn)* gene expression is XCI-dependent in multiple subjects, cell types and brain areas. *Rgn* is a calcium binding protein implicated in various processes including cell proliferation^18^ and shows an almost unanimous upregulation in X_1_ cells. *Rgn’s* negative effect on cell proliferation^19^ suggests that higher allelic expression of this X-linked proliferation suppressor might cause lower numbers of X_1_ cells. A similar pattern is observed for the *Rhoxf1*, albeit in fewer patients. Gain-of-function experiments for *Rhoxf1* revealed its role in reduction of proliferation in favor of differentiation^20^, a scenario similar to *Rgn*. The last gene with XCI-specific expression difference, *Sytl4* is a calcium-independent effector of Rab27 involved in membrane trafficking of secretory vesicles^21^. However both X_1_ and X_2_ cells show excess expression, signifying that *Sytl4* represents a common factor discordant between X_1_ and X_2_ cells.

Collectively, XCI-dependent differential expression of genes in human sc/snRNA-Seq datasets reflect functional inequivalence between cells with a different active homolog of X which, in turn, may contribute to disease features.

### X-chromosome related epigenetic erosion in dementia

Determining XCI by investigating SNVs also offers insight into disease effects on the consistency of X-inactivation. For instance, an increase in non-informative SNVs is indicative of epigenetic erosion at the chromosomal level. Likewise, an increased number of cells that expresses alleles from both X-chromosomes or an overall increase of proportion of discordant alleles, would denote erosion at the cellular level and stochastic level, respectively. To investigate the signatures of such epigenetic erosions, we plotted these variables for the various patient datasets (Extended Data Fig. 5). Whereas COPD, IPF and FTD patients do not differ from their controls, AD patients show a considerable increase in the proportion of SNVs (and hence genes) with biallelic expression across all cell types (Fig. 4a, p < 0.005), which was significant for OC brain regions (Fig. 4b, p < 0.01).

**Fig. 4.**
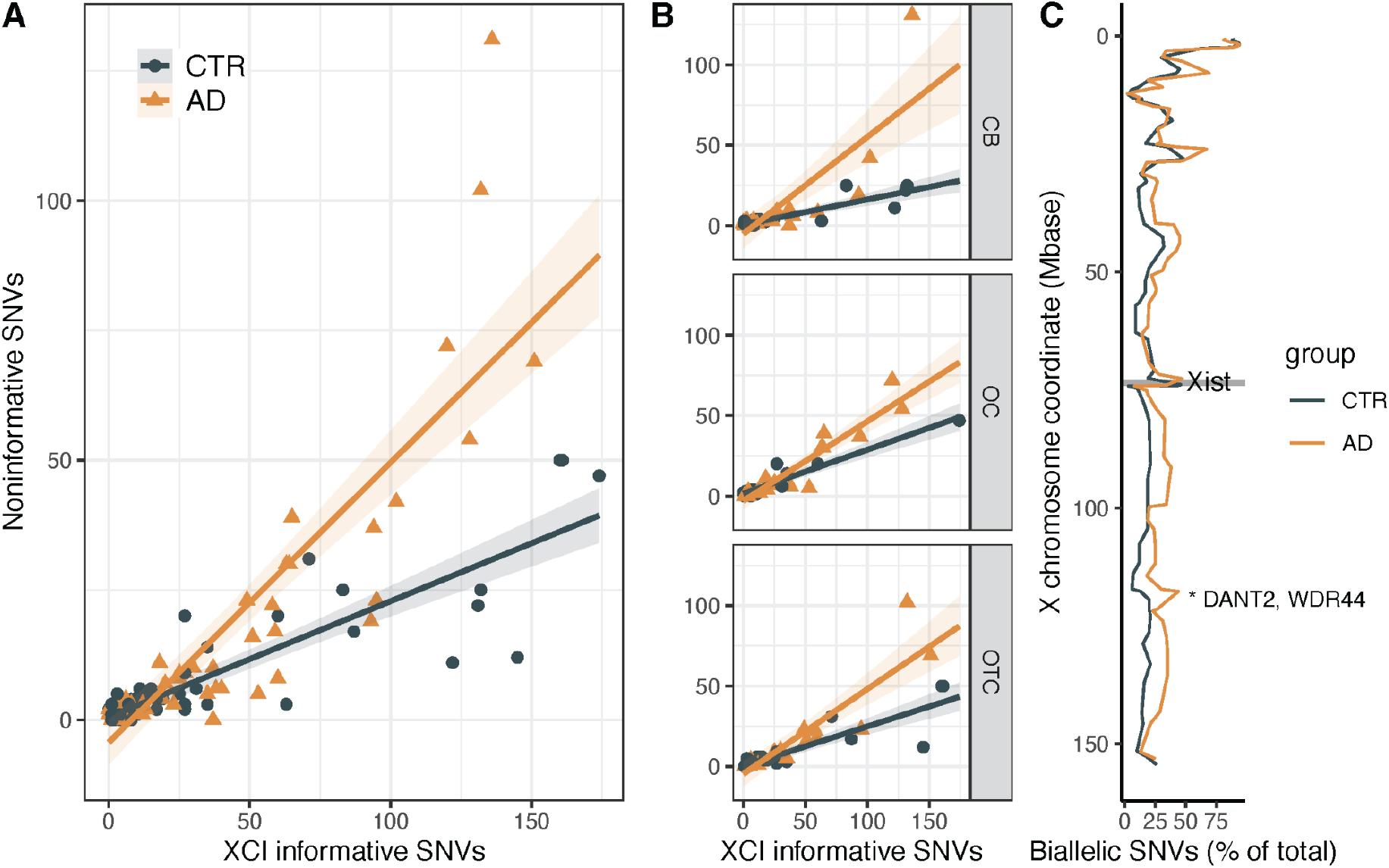
Epigenetic erosion analysis of XCI-informative versus non-informative SNVs in Alzheimer’s disease patients (AD) and controls (CTR). **A**. Numbers across all brain cell types and regions (p<0.005). **B**. Numbers per brain region (p <0.01 for OTC region, p = ns for CB and OC). The significant increase in noninformative SNVs in AD compared to CTR implies excess genetic erosion at the chromosomal level. (**C**) Percentage of biallelic cells across the X chromosome across a sliding window of 100 bp with gray rectangle indicating location of *Xist* gene. Panels A and B: p < 0.0001. Panel C: * = p<0.00001. CB = cerebellum, OC = occipital cortex, OTC = occipito-temporal cortex.

Next, we examined whether AD-specific biallelic expression may be explained by the existence of SNV clusters on the X-chromosome that escape XCI (Fig 4c, Extended data Fig. 6). Hereto, we sort SNVs by chromosome X coordinate and compare the distribution of XCI-informative versus non-informative SNVs between AD and CTR in a sliding window of 200 SNVs (2×2 independent χ2-test; 100 SNVs overlap between windows). Two adjacent regions with a higher proportion of biallelic cells in AD patients consistently show up (p-value < 10^−5^) and contain the genes *DANT2* (encoding a putative lncRNA) and *WDR44*, encoding a protein inhibiting ciliogenesis by interacting with Rab11^22^. Of note, impairment of cilia signaling and generation has been prominently linked to neurodegenerative diseases, including AD^22,23^.

## Discussion

Here we introduce a computational pipeline designed to determine X-chromosome inactivation status at single-cell level using solely sc/snRNA-seq datasets. The robustness of the algorithm is demonstrated with CAST x B6 mouse F1 hybrids, which have known genetic backgrounds. The method correctly separates alleles belonging to each genetic background and identifies the expected allelic reversal at *Xist* and an unexpected one at the *Pdbc1* locus due to haplotype-specific alternative splicing.

The XCISE algorithm reveals significant variation in X-chromosome inactivation among different cell types in all datasets. Noteworthy are the findings showing excess prevalence of active X_CAST_ in mouse skin cells, that is highest in fibroblasts, and reversed in dermal papillae cells. Differences also exist in the most active X-chromosome between (alveolar) macrophages in control subjects and lung patients, both COPD and IPF, and between microglia and other brain cell types in control subjects and AD patients. This disparity in XCI patterns in dermal papillae cells, macrophages and microglia may possibly be attributed to their distinct germ layer origin (mesoderm) compared to other skin and lung cell types (ectoderm and endoderm)^24,25^. A further observation concerns the differential gene expression between cells with opposite XCI. The number of DEGs depending on XCI and their presence in different individuals and cell types indicates that XCI may be responsible for functional variability within cell types. Multiple indicators suggest this phenomenon is biological rather than a technical or computational artifact. Notably, 12 of the 16 DEGs in lung patients originate from only four protein families. Moreover, in the brain, differential gene expression between X_1_ and X_2_ cells occurs across multiple brain regions in a similar direction in the vast majority of cases.

Finally, we noted a substantial increase in cells expressing alleles from both X homologs across different brain regions and cell types of AD patients compared to controls. While this observation strongly suggests a potential loss of epigenetic marks that effectuate silencing of the second X-chromosome^26^, further studies need to investigate causality and the sequence of events during disease development.

## Methods

### Data sources

#### Mouse skin dataset

The data are available through EBI Bioostudies portal under accession number E-MTAB-12181, https://www.ebi.ac.uk/biostudies/arrayexpress/studies/E-MTAB-12181

#### Mouse fibroblast dataset

The data are available through EBI Bioostudies portal under accession number E-MTAB-6385, https://www.ebi.ac.uk/biostudies/arrayexpress/studies/E-MTAB-6385

#### Lung disease (IPF, COPD) patients and controls

The data are available through the Gene Expression Omnibus at https://www-ncbi-nlm-nih-gov/geo with accession number GSE136831

#### Frontotemporal dementia (FTD) patients and controls

The data are available through the Gene Expression Omnibus at https://www-ncbi-nlm-nih-gov/geo with accession number GSE163122

#### Alzheimer’s Disease (AD) patients and controls

The data reported are available through Gene Expression Omnibus at https://www.ncbi.nlm.nih.gov/geo with accession number GSE148822

### X-chromosome inactivation analysis

Raw data have been downloaded from corresponding repositories and included FASTQ files and tables linking cell-specific barcodes, cell type and when applicable, brain regions. Mouse and human sc/snRNAseq data were aligned using STAR mapper (v.2.7.10b) against corresponding genome references and gene builds (GRCm39, Ensembl release 110 and GRCh38, Ensembl release 109 respectively). SNVs were called using bcftools (v.1.11). Uncommon polymorphisms (that have allele frequency below 1% in GNOMAD database and might represent RNA editing events, sequencing errors and misalignments) were removed. Heterozygous polymorphisms were used for second-round alignment using STARsolo (for 10x chromium) or STAR alignment (Smart-seq) with enabled functionality for unbiased calling of alleles at heterozygous positions, known as WASP functionality^27^ and implemented in STAR. The XCISE algorithm (deposited at https://github.com/Vityay/XCISE) uses an iterative algorithm to find best solution aiming for the highest proportion between total count of UMIs that are concordant *vs* those that are discordant with the cellular pool to which this cell was assigned. The solution is found by starting from random assignment of SNV alleles to haplotypes (or marking an SNV as biallelicly expressed, which is XCI-noninformative) and iterating over all SNV position and haplotype options retaining SNV reassignments if they increase score function. The procedure continues until no further improvements can be made. Within a solution, each cell is assigned to the X_1_, X_2_ or X_BOTH_ pool based on the dominant haplotype, supported by 90% or more UMIs. By default, the algorithm iterates 100 times, starting from random assignment of SNV alleles to haplotypes. The best solution is reported as the final result. Running the XCISE pipeline on BAM files yields files containing the general statistics of the run (e.g. allele discordance rate, number of cells per cell pool, number of total and XCI-informative SNVs, text file), XCI status per cell (tab-separated file), position, reads support and XCI-informativeness status for each SNVs (VCF file). The manual and detailed description of pipeline steps and commands used for running the pipeline for mouse and human datasets are added to the repository. Postprocessing and plotting of XCISE results were done with custom R scripts shared through the same github repository.

### Bayesian inference of X_1_ distribution

Using the number of observed occurrences of X_1_ and total number of phased cells per in each cell type (and brain region in case of dementia patient data), we modeled the number of observed occurrences in each individual of X_1,i_ as a binomial random variable and calculated its probability using RStan (version 2.32.5) under R (version 4.3.1). The probabilities per individual (θ_i_) were modeled as beta-distributed random variables, with each having a non-informative prior specified as a Beta(1, 1) distribution. We sampled from the posterior distribution of the model parameters using a total of 4000 iterations, discarding the first 2000 iterations as warm-up to ensure convergence and computed summary statistics including posterior means and standard deviation, and the probabilities θ_i_ representing the likelihood of observing X_1_ for each individual and cell type (and brain region).

## Acknowledgements

We thank Prof. M.C. Nawijn and M. Banchero for sharing scRNA-seq data used in pipeline development and discussions of the initial results.

