## Supplemental Table 1 for "Single-cell X-chromosome inactivation analysis links biased chimerism to differential gene expression and epigenetic erosion"

**MOUSE**

| ID | Type | XCI-inform SNPs | Non-inf SNVs | Cells X1 | Cells X2 | Cells, XCI-ed | Cells, X-Both | Biallelic_rate | Discordance rate per UMI, % | X1,% | |
|---|---|---|---|---|---|---|---|---|---|---|---|
| E-MTAB-12181 | scRNA-seq skin F1 Mou | 1555 | 67 | 148 | 218 | 366 | 17 | 4.44 | 1.31 | 40.44 | |
| E-MTAB-6385 | scRNA-seq fibroblasts F | 1965 | 36 | 99 | 273 | 372 | 2 | 0.53 | 0.97 | 26.61 | |
| E-MTAB-10148 | scRNA-seq male F1 Mo | 119 | 649 | | | | | | | | |
| SNV pruning (removal of 90% of SNVs), random replicates | | | | | | | | | | | |
| E-MTAB-12181, rep 1 | scRNA-seq F1 Mouse sk | 158 | 8 | 214 | 145 | 359 | 15 | 4.01 | 1.31 | 40.39 | 3 SNVs with diff phase - 2 in Xist, 1 in Pbdc1 |
| E-MTAB-12181, rep 2 | scRNA-seq F1 Mouse sk | 167 | 5 | 214 | 149 | 363 | 9 | 2.42 | 1.31 | 41.05 | 3 SNV with diff phase in Xist |
| E-MTAB-12181, rep 3 | scRNA-seq F1 Mouse sk | 169 | 9 | 216 | 150 | 366 | 9 | 2.40 | 1.31 | 40.98 | 2 SNV with diff phase in Xist |
| E-MTAB-12181, rep 4 | scRNA-seq F1 Mouse sk | 152 | 5 | 215 | 148 | 363 | 10 | 2.68 | 1.31 | 40.77 | 3 SNV with diff phase in Xist |
| E-MTAB-12181, rep 5 | scRNA-seq F1 Mouse sk | 180 | 7 | 211 | 148 | 359 | 13 | 3.49 | 1.31 | 41.23 | 4 SNVs with diff phase - 3 in Xist, 1 in Pbdc1 |

**LUNG COHORT**

| Patient ID | Type | XCI-inform SNPs | Non-inf SNVs | Cells X1 | Cells X2 | Cells, XCI-ed | Cells, X-Both | Biallelic_rate | Discordance rate per UMI, % | X1,% | Sequencing saturation | Het SNVs on X | Estmated cells | Reads Mapped to Gene |
|---|---|---|---|---|---|---|---|---|---|---|---|---|---|---|
| 157I | IPF, 66 years | 45 | 38 | 779 | 788 | 1567 | 93 | 5.60 | 3.78 | 49.71 | 0.873398 | 263849575 | 6044 | 7562 |
| 098C | Control, 41 years | 45 | 36 | 1330 | 2177 | 3507 | 256 | 6.80 | 2.95 | 37.92 | 0.800444 | 200132305 | 6631 | 7965 |
| 225I | IPF, 70 years | 25 | 22 | 345 | 522 | 867 | 58 | 6.27 | 5.00 | 39.79 | 0.684226 | 197205786 | 6103 | 7453 |
| 1372C | Control, 21 years | 26 | 23 | 484 | 486 | 970 | 74 | 7.09 | 4.20 | 49.90 | 0.877063 | 151588278 | 4629 | 5763 |
| 133C | Control, 32 years | 23 | 17 | 1233 | 1996 | 3229 | 197 | 5.75 | 3.34 | 38.19 | 0.656405 | 126979520 | 5988 | 7086 |
| 184CO | COPD, 55 years | 37 | 16 | 1019 | 1875 | 2894 | 183 | 5.95 | 3.15 | 35.21 | 0.888885 | 122401521 | 2609 | 3300 |
| 174I | IPF, 67 years | 21 | 15 | 247 | 528 | 775 | 48 | 5.83 | 4.88 | 31.87 | 0.721899 | 106416381 | 5694 | 6906 |
| 052CO | COPD, 62 years | 19 | 9 | 247 | 270 | 517 | 22 | 4.08 | 2.42 | 47.78 | 0.883948 | 87814002 | 2493 | 3368 |
| 003C | Control, 67 years | 1 | 1 | 1 | 2 | 3 | 0 | - | - | 33.33 | 0.786687 | 87650592 | 5998 | 7549 |
| 439C | Control, 66 years | 30 | 2 | 82 | 150 | 232 | 11 | 4.53 | 4.36 | 35.34 | 0.787262 | 81127308 | 4619 | 6083 |
| 209I | IPF, 65 years | 14 | 9 | 456 | 724 | 1180 | 79 | 6.27 | 3.69 | 38.64 | 0.702196 | 79101139 | 6576 | 8633 |
| 221I | IPF, 67 years | 16 | 2 | 26 | 60 | 86 | 3 | 3.37 | 2.78 | 30.23 | 0.679307 | 77202425 | 5181 | 6458 |
| 253C | Control, 66 years | 6 | 2 | 14 | 34 | 48 | 0 | - | - | 29.17 | 0.909069 | 72891867 | 4664 | 6646 |
| 217CO | COPD, 70 years | 19 | 8 | 32 | 45 | 77 | 0 | - | - | 41.56 | 0.892598 | 68723739 | 2280 | 3135 |
| 237CO | COPD, 57 years | 30 | 15 | 47 | 89 | 136 | 9 | 6.21 | 4.55 | 34.56 | 0.897077 | 60856532 | 1702 | 2307 |
| 056CO | COPD, 57 years | 24 | 12 | 70 | 131 | 201 | 11 | 5.19 | 4.01 | 34.83 | 0.851424 | 58663626 | 1802 | 2434 |
| 207CO | COPD, 60 years | 18 | 7 | 44 | 56 | 100 | 3 | 2.91 | 2.33 | 44.00 | 0.875962 | 55034996 | 2099 | 2966 |
| 178CO | COPD, 58 years | 37 | 18 | 106 | 118 | 224 | 8 | 3.45 | 2.88 | 47.32 | 0.752808 | 52231736 | 2543 | 3694 |
| 296C | Control, 80 years | 16 | 5 | 12 | 15 | 27 | 0 | - | - | 44.44 | 0.822947 | 48360070 | 2834 | 3738 |
| 235CO | COPD, 61 years | 16 | 4 | 412 | 445 | 857 | 64 | 6.95 | 4.31 | 48.07 | 0.730884 | 37387358 | 2818 | 3701 |
| 454C | Control, 48 years | 4 | 1 | 3 | 11 | 14 | 2 | 12.50 | 9.52 | 21.43 | 0.84747 | 30645128 | 2080 | 3168 |
| 396C | Control, 37 years | 10 | 9 | 121 | 252 | 373 | 45 | 10.77 | 6.06 | 32.44 | 0.839774 | 15539199 | 931 | 1354 |
| Excluded patients (after sex chromosomal read check)* | | | | | | | | | | | | | | |
| 137CO | COPD, 73 years | 0 | | 0 | 0 | 0 | 0 | #N/A | - | #N/A | 0.977169 | 59128555 | 434 | 705 |
| 59I | IPF, 66 years | 0 | | 0 | 0 | 0 | 0 | #N/A | - | #N/A | 0.832948 | 43539300 | 5043 | 7610 |
| 192C | Control, 62 years | 19 | | 553 | 838 | 1391 | 123 | 8.12 | 3.79 | 39.76 | 0.873562 | 306336023 | 6600 | 8272 |
| 002C | Control, 25 years | 2 | | 6 | 9 | 15 | 0 | - | - | 40.00 | 0.914957 | 47782055 | 3105 | 4282 |

| | | | | | | | | | | | | | |
|---|---|---|---|---|---|---|---|---|---|---|---|---|---|
| 065C | Control, 66 years | 7 | 6 | 11 | 17 | 0 | - | - | 35.29 | 0.888823 | 35629022 | 1695 | 2248 |

**DEMENTIA COHORTS**

| Patient ID | Type | XCI-inform SNPs | Non-inf SNVs | Cells X1 | Cells X2 | Cells, X-Both | Biallelic_rate, % | Concordant | Discordant | X1,% | |
|---|---|---|---|---|---|---|---|---|---|---|---|
| IB5431 | snRNA-seq Control | 1226 | 380 | 6038 | 7703 | 1367 | 9.05 | 32850 | 1778 | 43.94 | |
| IB5508 | snRNA-seq Control | 941 | 378 | 5004 | 5024 | 1084 | 9.76 | 21607 | 1359 | 49.90 | |
| IB5691 | snRNA-seq AD | 823 | 544 | 2170 | 3787 | 586 | 8.96 | 10087 | 663 | 36.43 | |
| IB5869 | snRNA-seq AD | 986 | 564 | 2929 | 6194 | 989 | 9.78 | 15542 | 1111 | 32.11 | |
| IB5888 | snRNA-seq AD | 521 | 280 | 1204 | 1430 | 156 | 5.59 | 3542 | 169 | 45.71 | |
| IB6001 | snRNA-seq AD | 990 | 633 | 3617 | 4682 | 877 | 9.56 | 13727 | 1025 | 43.58 | |
| IB6125 | snRNA-seq Control | 833 | 664 | 2425 | 2793 | 476 | 8.36 | 8297 | 536 | 46.47 | |
| CTR018 | snRNA-seq Control | 1324 | 727 | 4187 | 7147 | 1231 | 9.80 | 29066 | 1602 | 36.94 | |
| CTR081 | snRNA-seq Control | 638 | 212 | 2676 | 5219 | 573 | 6.77 | 14135 | 683 | 33.89 | |
| CTR148 | snRNA-seq Control | 649 | 273 | 2130 | 2861 | 427 | 7.88 | 10526 | 549 | 42.68 | |
| FTD014 | snRNA-seq FTD | 743 | 324 | 2285 | 3532 | 453 | 7.22 | 12188 | 583 | 39.28 | |
| FTD024 | snRNA-seq FTD | 886 | 268 | 3863 | 4224 | 688 | 7.84 | 14719 | 860 | 47.77 | |
| FTD038 | snRNA-seq FTD | 733 | 421 | 2213 | 5332 | 759 | 9.14 | 13777 | 963 | 29.33 | |
| FTD073 | snRNA-seq FTD | 1032 | 314 | 1598 | 9013 | 764 | 6.72 | 19867 | 907 | 15.06 | |
| FTD083 | snRNA-seq FTD | 573 | 247 | 1325 | 2227 | 295 | 7.67 | 7128 | 358 | 37.30 | |
| FTD6243 | snRNA-seq FTD | 202 | 87 | 560 | 628 | 111 | 8.55 | 2576 | 155 | 47.14 | |
| CTR097 | snRNA-seq Control mal | 143 | | 230 | 255 | 485 | 17 | 3.39 | 2.95 | 47.42 | PAR region |

* We found 5 subjects with very low numbers of cells in which XCI status is detected.
- XCI-informative SNVs are not called from one COPD and one IPF patient related to low number of cells (137CO and 59I)
- The three remaining subjects show substantial expression of transcripts from the male-specific region of the Y chromosome (MSY), suggesting contamination or misrecorded sex (Controls 002C, O65C and 192C)
