## Supplemental Figures for "Single-cell X-chromosome inactivation analysis links biased chimerism to differential gene expression and epigenetic erosion"

### Supplementary Figures

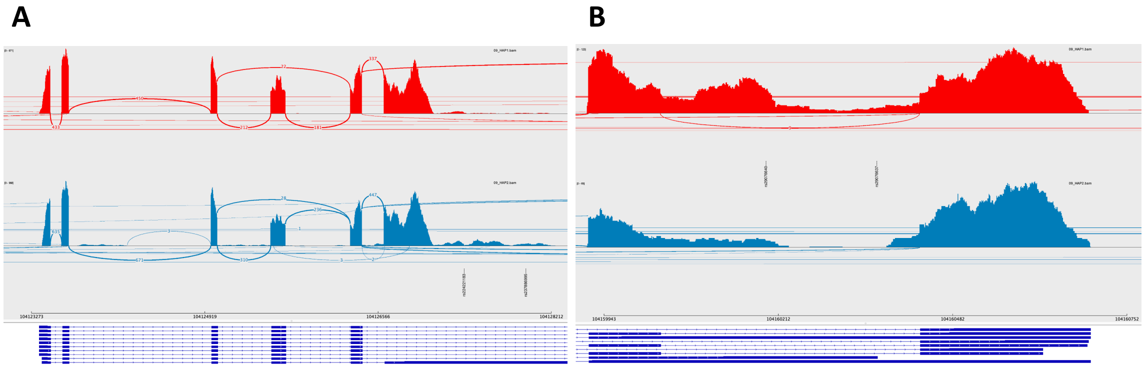


**Supplementary Fig. 1. Haplotype-specific alternative splicing of *Pbdc1* explaining its biased allelic expression in mouse skin cells.** *Pbdc1* is an X chromosomal gene known to escape XCI in mice, which consequently leads to expression of both X alleles ('biallelic expression'). The *Pbdc1* harbors 11 SNVs, seven of which are predicted as expressed by both X chromosomes. In addition, 2 SNVs (rs224221183, rs237886995) are biased towards expression of the CAST allele and 2 SNVs (rs29076640, rs29076637) towards the B6 allele. **A,B.** Sashimi plots showing location of two pairs of XCI-informative SNVs in the biallelicly expressed *Pbdc1* gene. The three tracks represent from top to bottom: exon coverage (filled areas) and reads crossing splice junctions (arcs) in cells with $XCI_{B6}$ (red color; active $X_{CAST}$ and $Xist_{B6}$ expression); exon coverage and splice junction usage in cells with $XCI_{CAST}$ (blue color; active $X_{B6}$ and $Xist_{CAST}$ expression) with position of the biased SNVs indicated (vertical text displaying reference IDs of SNVs); exon-intron structure of known *Pbdc1* transcripts. Haplotype-biased alternative splicing results in predominant coverage of these SNVs from either the cells with active $X_{CAST}$ (panel A) or active $X_{B6}$ (panel B), identifying them as XCI-informative SNVs that are located inside a gene that escapes X inactivation.

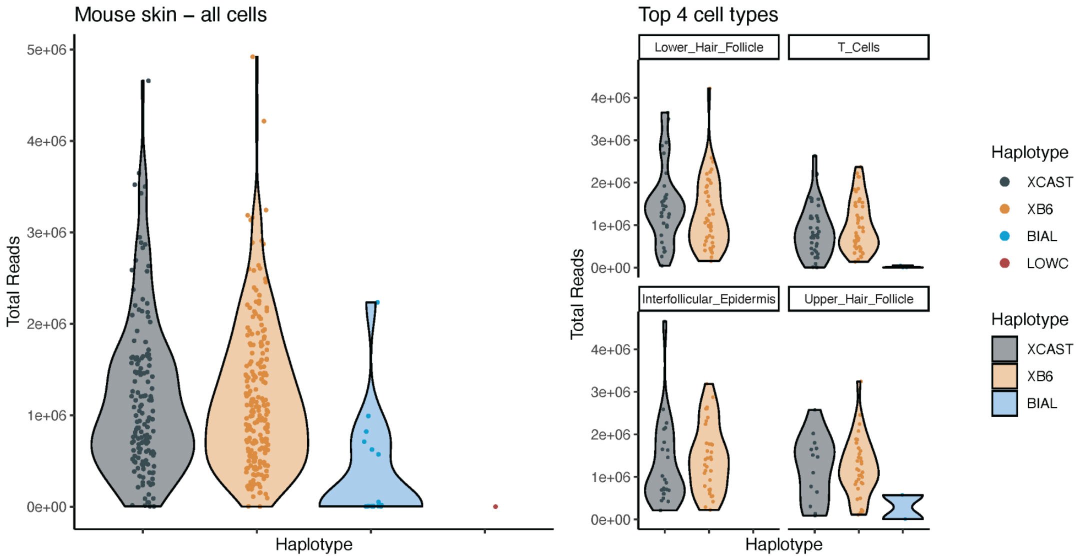


**Supplementary Fig. 2.** Total reads mapped to genes per cell in F1 mouse skin scRNA-seq data. Colors designate classification of cells by the XCISE algorithm: dark gray - active $X_{CAST}$, orange - active $X_{B6}$, blue - biallelic expression, red - unknown, representing a small number of reads overlapping haplotype-informative SNVs. Cells with biallelic expression at X chromosome do not seem to be due to doublets (two cells in a well), as they deliver significantly less reads compared to other cells.

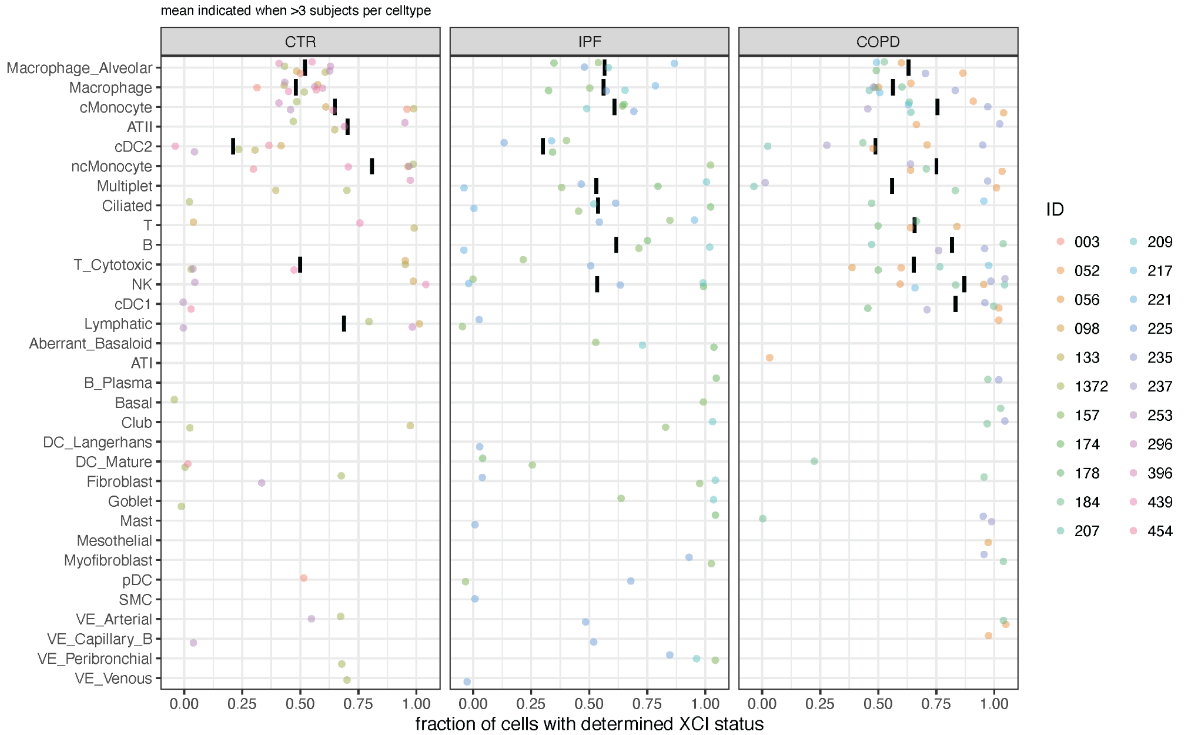


**Supplementary Fig. 3. Fraction of cells with determined XCI status per cell type in a public dataset of control and lung patients.** Data are included when in a subject for the cell type indicated >= 100 cells are sequenced and >= 10 cells have their XCI status determined. Cell types are sorted top to bottom according to the total number of cells phased. Mean value is indicated by black bars when XCI status was called for 4 or more subjects.

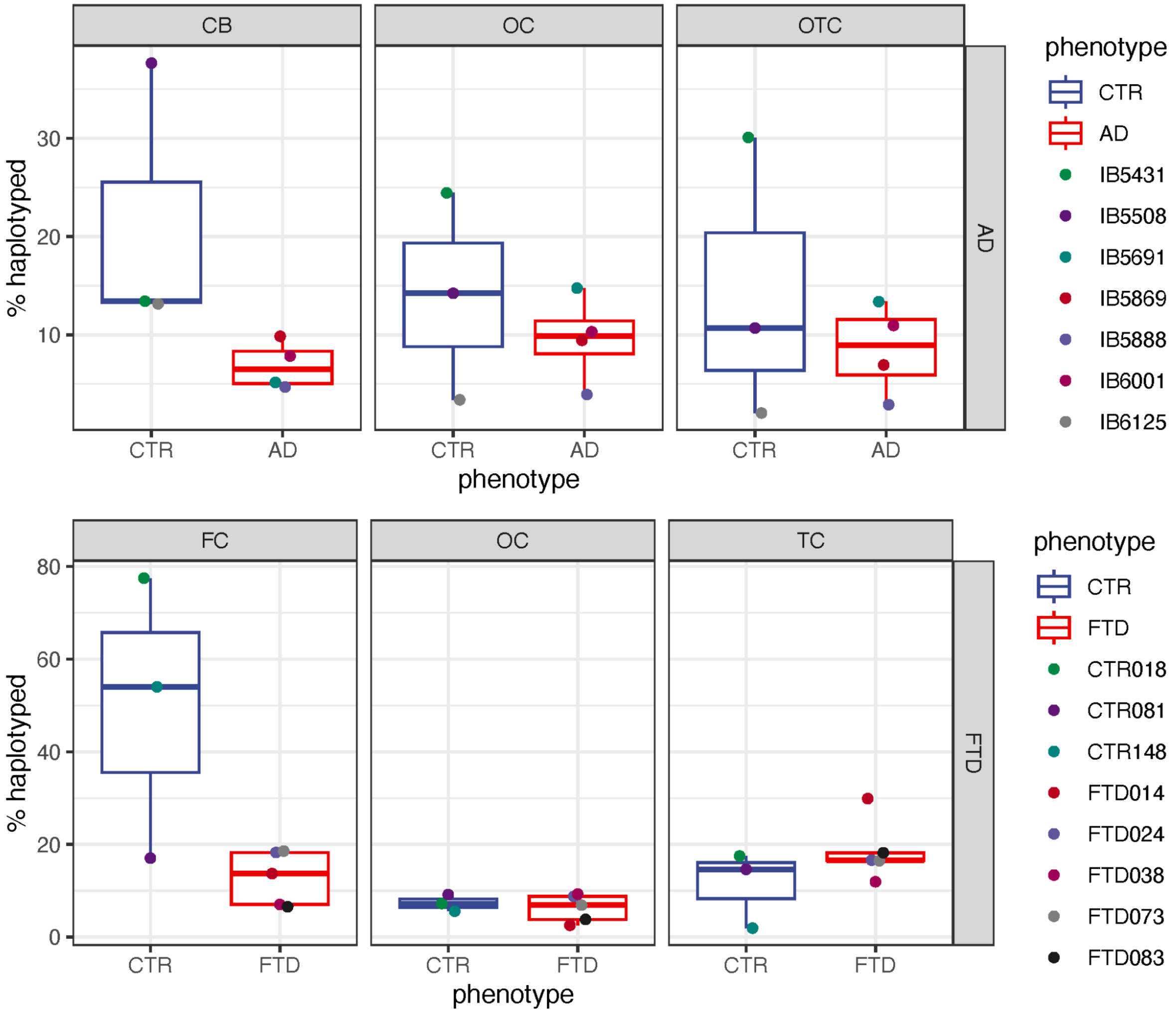


**Supplementary Fig. 4. Haplotyping in dementia cohorts.** Percentage of total cells successfully phased for X chromosomes per cohort and brain region. AD and FTD indicate the cohort. OC = occipital cortex, OTC = occipito-temporal cortex, CB = cerebellum, FC = frontal cortex, TC = temporal cortex.

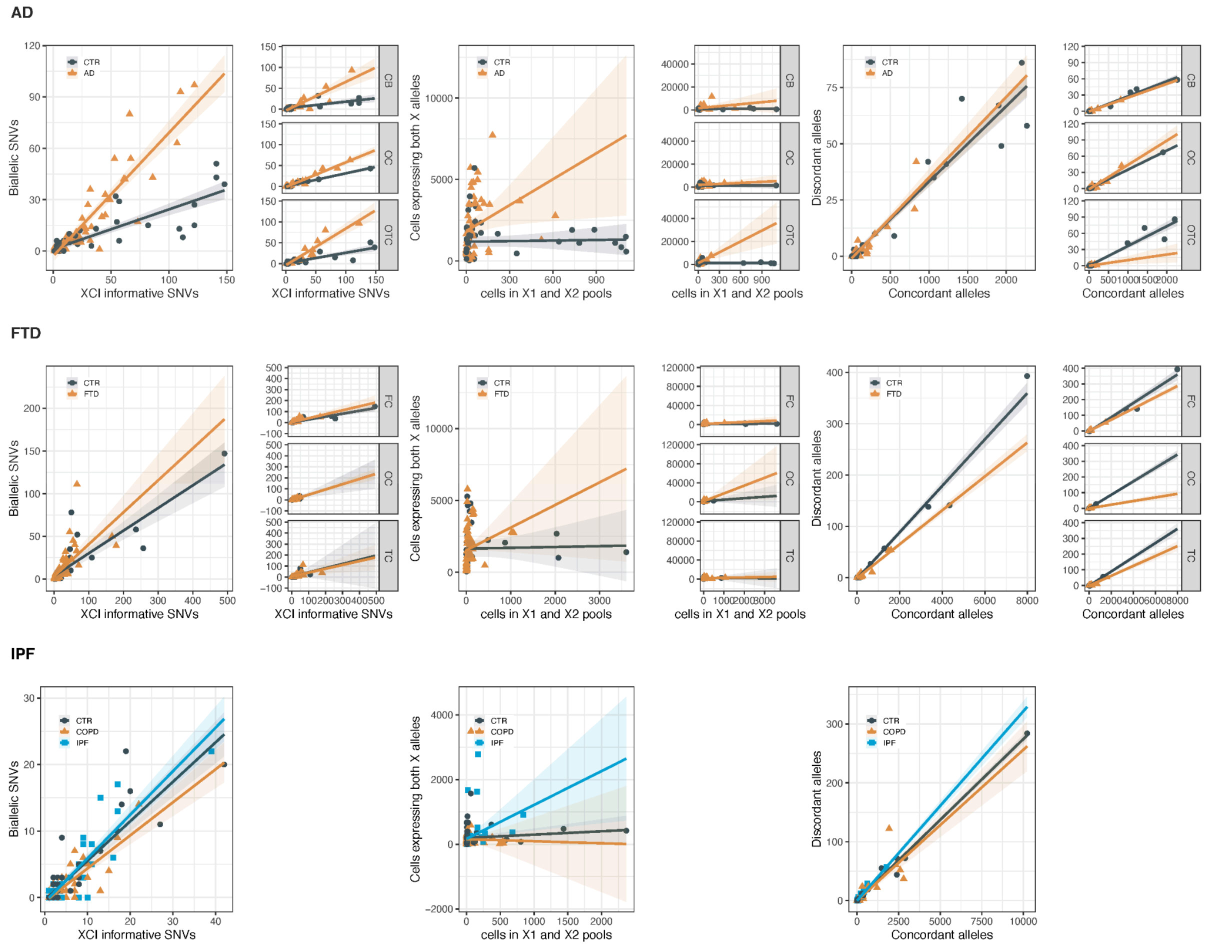


**Supplementary Fig. 5.** Analysis of epigenetic erosion at the chromosomal level (XCI informative *versus* biallelic SNVs; left panels), cell level (cells in X1 and X2 pools *versus* cells expressing both; middle panels) and stochastic level (Concordant *versus* Discordant alleles; right panels) in all patient cohorts. CTR = controls within the cohort, AD=Alzheimer's disease patients and non-dementia controls, FTD = fronto-temporal dementia and non-dementia controls, IPF = Interstitial pulmonary fibrosis, COPD = chronic obstructive lung disease and their controls. CB = cerebellum, OC = occipital cortex, OTC = occipito-temporal cortex, FC = frontal cortex, TC = temporal cortex.

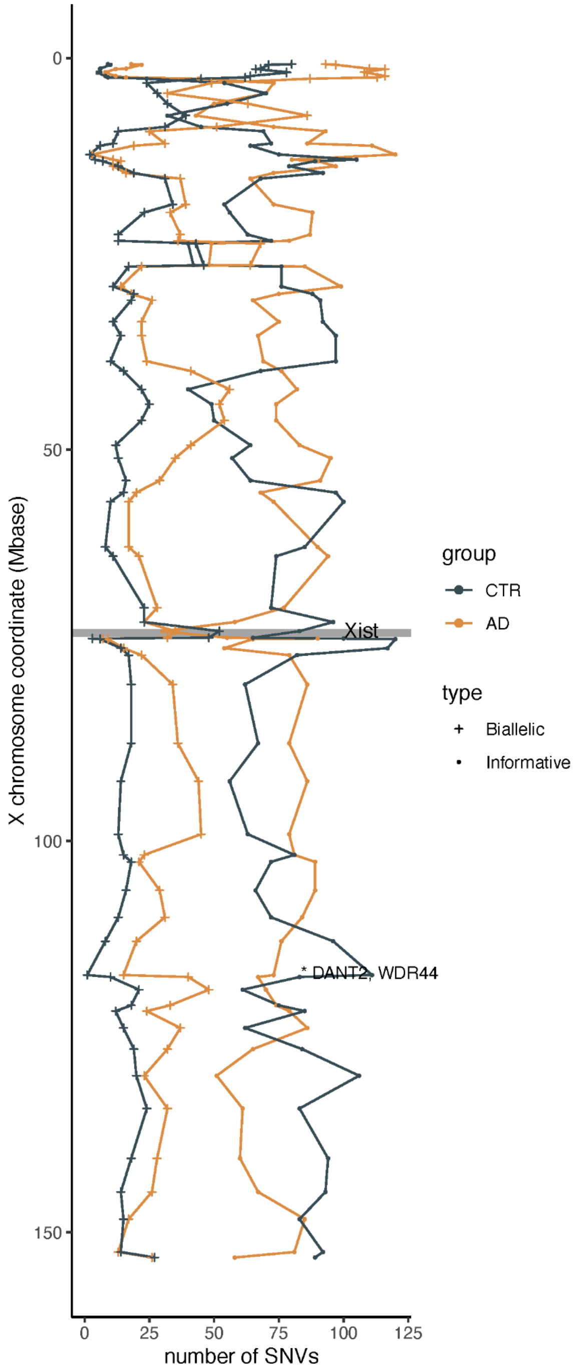


**Supplementary Fig. 6.** Number of XCI-informative and biallelic SNVs in Alzheimer's disease patients (AD) and controls (CTR) across a sliding window of 200 bp. Gray rectangle indicates the location of *Xist* gene. * = genes with p<0.00001 for AD versus CTR.
