## Supplemental Table 2 for "Single-cell X-chromosome inactivation analysis links biased chimerism to differential gene expression and epigenetic erosion"

--- gene_id: ENSMUSG00000104699 gene_name: Rps4x-ps gene_location: 5:3925362-3926154 ---

| Set | LogFC | LogCPM | P-val |
|---|---|---|---|
| Interfollicular_Epidermis | 6.87127479580715 | 5.72571100563765 | 4.24228433825961e-30 |
| Upper_Hair_Follicle | 6.35939557471073 | 5.79620645306141 | 1.75939701258799e-08 |
| T_Cells | 4.81741255186015 | 3.63452548635707 | 2.93937559510983e-09 |
| Lower_Hair_Follicle | 5.69825311039212 | 5.81842370439093 | 6.12081085733889e-31 |

--- gene_id: ENSMUSG00000057762 gene_name: Gm6169 gene_location: 13:97234716-97235745 ---

| Set | LogFC | LogCPM | P-val |
|---|---|---|---|
| Lower_Hair_Follicle | 2.69569544040154 | 2.73513172180023 | 1.52729784268442e-05 |
| Interfollicular_Epidermis | 2.83365077203222 | 2.93226861288009 | 9.81555933888701e-05 |

--- gene_id: ENSMUSG00000031662 gene_name: Snx20 gene_location: 8:89353191-89362756 ---

| Set | LogFC | LogCPM | P-val |
|---|---|---|---|
| Upper_Hair_Follicle | -10.740430499353 | 5.73999203071908 | 0.000425902657453564 |
| T_Cells | -6.57363400119543 | 3.17091296096738 | 0.00670740752629547 |

--- gene_id: ENSMUSG00000031320 gene_name: Rps4x gene_location: X:101228547-101233000 ---

| Set | LogFC | LogCPM | P-val |
|---|---|---|---|
| Lower_Hair_Follicle | -1.78243042309897 | 6.49742589211866 | 5.89978210945234e-08 |
| Interfollicular_Epidermis | -1.54648990692947 | 6.29178948500696 | 0.000629842432810218 |

--- gene_id: ENSMUSG00000117428 gene_name: Gm4833 gene_location: 18:8051203-8051992 ---

| Set | LogFC | LogCPM | P-val |
|---|---|---|---|
| Lower_Hair_Follicle | 2.64594791237354 | 1.56123442197253 | 2.71693896355171e-07 |
| Interfollicular_Epidermis | 2.34273333398438 | 1.49064623253777 | 0.000629842432810218 |

--- gene_id: ENSMUSG00000053536 gene_name: Cstf2t gene_location: 19:31060237-31064469 ---

| Set | LogFC | LogCPM | P-val |
|---|---|---|---|
| Upper_Hair_Follicle | -9.92900866053624 | 4.47768823783989 | 0.00111013686196283 |

--- gene_id: ENSMUSG00000026389 gene_name: Steap3 gene_location: 1:120118487-120200435 ---

| Set | LogFC | LogCPM | P-val |
|---|---|---|---|
| Interfollicular_Epidermis | -6.33894589728149 | 3.12710323932477 | 0.00877057321313802 |

--- gene_id: ENSMUSG00000029096 gene_name: Htra3 gene_location: 5:35809367-35837126 ---

| Set | LogFC | LogCPM | P-val |
|---|---|---|---|
| T_Cells | -3.36649025915093 | 3.0915174230506 | 0.00193406115942337 |

--- gene_id: ENSMUSG00000019849 gene_name: Prep gene_location: 10:44943299-45043294 ---

| Set | LogFC | LogCPM | P-val |
|---|---|---|---|
| T_Cells | 6.24463919182603 | 4.90990833546587 | 0.00706759509228491 |

--- gene_id: ENSMUSG00000001665 gene_name: Gstt3 gene_location: 10:75609949-75617248 ---

| Set | LogFC | LogCPM | P-val |
|---|---|---|---|
| T_Cells | -3.97684333576636 | 2.57372150046759 | 0.00223537304195952 |

--- gene_id: ENSMUSG00000037031 gene_name: Tspan15 gene_location: 10:62021175-62067030 ---

| Set | LogFC | LogCPM | P-val |
|---|---|---|---|
| Mixed_Cluster | -10.7629783011089 | 6.83565011716282 | 0.00485094649090404 |

--- gene_id: ENSMUSG00000035493 gene_name: Tgfbi gene_location: 13:56757336-56787375 ---

| Set | LogFC | LogCPM | P-val |
|---|---|---|---|
| T_Cells | -2.70490979636006 | 3.26584627974513 | 0.000984844988950389 |

--- gene_id: ENSMUSG00000023224 gene_name: Serping1 gene_location: 2:84595731-84605788 ---

| Set | LogFC | LogCPM | P-val |
|---|---|---|---|
| Upper_Hair_Follicle | -3.69715389883555 | 4.11472862334262 | 0.00494583367397433 |

```
--- gene_id: ENSMUSG00000023156 gene_name: Rpp14 gene_location: 14:14377939-14389406 ---
Set                          LogFC                 LogCPM              P-val
T_Cells                      -6.17354045237288     2.97850047244772    0.00396444388666427
--- gene_id: ENSMUSG00000053730 gene_name: Tmem39b gene_location: 4:129570148-129590631 ---
Set                          LogFC                 LogCPM              P-val
T_Cells                      -5.69547688566817     2.13478006725325    0.00667305182491627
--- gene_id: ENSMUSG00000039745 gene_name: Htatip2 gene_location: 7:49408863-49423723 ---
Set                          LogFC                 LogCPM              P-val
T_Cells                      -6.2001630674352      4.42730345586871    0.00491662815936797
--- gene_id: ENSMUSG00000028480 gene_name: Glipr2 gene_location: 4:43957401-43979118 ---
Set                          LogFC                 LogCPM              P-val
T_Cells                      -7.54993989858607     4.65000227307137    0.00223537304195952
--- gene_id: ENSMUSG00000038463 gene_name: Olfml2b gene_location: 1:170472101-170510358 ---
Set                          LogFC                 LogCPM              P-val
Mixed_Cluster                -9.5849974870549      7.29475250415079    0.00885739029605146
--- gene_id: ENSMUSG00000028378 gene_name: Ptgr1 gene_location: 4:58965439-58987119 ---
Set                          LogFC                 LogCPM              P-val
T_Cells                      -3.39275593804221     2.25159273312987    0.00434241283203604
--- gene_id: ENSMUSG00000073888 gene_name: Ccl27a gene_location: 4:41769467-41774247 ---
Set                          LogFC                 LogCPM              P-val
T_Cells                      -3.3514214918508      4.42797248073957    3.08894716714478e-05
--- gene_id: ENSMUSG00000027088 gene_name: Phospho2 gene_location: 2:69619967-69630349 ---
Set                          LogFC                 LogCPM              P-val
T_Cells                      -6.34631293633654     2.9240624593339     0.00670740752629547
--- gene_id: ENSMUSG00000025492 gene_name: Ifitm3 gene_location: 7:140589499-140590683 ---
Set                          LogFC                 LogCPM              P-val
T_Cells                      -3.29737932517524     6.06690687709404    0.000172601698813066
--- gene_id: ENSMUSG00000024247 gene_name: Pkdcc gene_location: 17:83522721-83532499 ---
Set                          LogFC                 LogCPM              P-val
Upper_Hair_Follicle          -9.55926184797492     4.35419762445909    0.00831319482634756
--- gene_id: ENSMUSG00000043681 gene_name: Fam25c gene_location: 14:34073838-34077390 ---
Set                          LogFC                 LogCPM              P-val
T_Cells                      -4.70880296052943     3.73367604082361    0.000792829749819312
--- gene_id: ENSMUSG00000029781 gene_name: Fkbp9 gene_location: 6:56809044-56856343 ---
Set                          LogFC                 LogCPM              P-val
T_Cells                      -4.36504743280962     2.83668626560113    0.00434241283203604
--- gene_id: ENSMUSG00000026826 gene_name: Nr4a2 gene_location: 2:56996842-57014015 ---
Set                          LogFC                 LogCPM              P-val
Lower_Hair_Follicle          5.7316463148292       4.59943491802902    0.00148042633873175
--- gene_id: ENSMUSG00000028393 gene_name: Alad gene_location: 4:62427406-62438155 ---
Set                          LogFC                 LogCPM              P-val
T_Cells                      -3.99518023684011     2.84868076011822    0.00498854951602768
--- gene_id: ENSMUSG00000024965 gene_name: Fermt3 gene_location: 19:6976326-6996837 ---
Set                          LogFC                 LogCPM              P-val
Upper_Hair_Follicle          -5.33969399069764     2.84727214101629    0.00494583367397433
--- gene_id: ENSMUSG00000027709 gene_name: Mccc1 gene_location: 3:36013461-36054827 ---
```

```
Set                            LogFC                 LogCPM               P-val
T_Cells                        -7.30117004903378     4.05862653384223     0.00670740752629547
--- gene_id: ENSMUSG00000031503 gene_name: Col4a2 gene_location: 8:11362805-11499287 ---
Set                            LogFC                 LogCPM               P-val
T_Cells                        -5.39135181800367     3.14210159826472     0.00991592834485912
--- gene_id: ENSMUSG00000075602 gene_name: Ly6a gene_location: 15:74866726-74869880 ---
Set                            LogFC                 LogCPM               P-val
T_Cells                        -4.45074316425878     6.3100317909252      0.000118983303250242
--- gene_id: ENSMUSG00000055632 gene_name: Hmcn2 gene_location: 2:31204427-31350750 ---
Set                            LogFC                 LogCPM               P-val
Mixed_Cluster                  -10.5494453371743     7.53379961343298     0.0046277826535293
--- gene_id: ENSMUSG00000028716 gene_name: Pdzk1ip1 gene_location: 4:114945905-114951096 ---
Set                            LogFC                 LogCPM               P-val
T_Cells                        -3.8137347075548      2.384286957065       0.00233898780425338
--- gene_id: ENSMUSG00000004791 gene_name: Pgf gene_location: 12:85213409-85224564 ---
Set                            LogFC                 LogCPM               P-val
Mixed_Cluster                  -10.0735209293783     7.7992902432437      0.00636243947550316
--- gene_id: ENSMUSG00000028899 gene_name: Taf12 gene_location: 4:132001686-132023077 ---
Set                            LogFC                 LogCPM               P-val
Lower_Hair_Follicle            8.1877653999303       4.56878385591333     1.76406463776194e-05
--- gene_id: ENSMUSG00000031146 gene_name: Plp2 gene_location: X:7534180-7537629 ---
Set                            LogFC                 LogCPM               P-val
Interfollicular_Epidermis      -1.53172484894829     6.13830749842025     0.00877057321313802
--- gene_id: ENSMUSG00000057092 gene_name: Fxyd3 gene_location: 7:30767597-30776129 ---
Set                            LogFC                 LogCPM               P-val
T_Cells                        -4.11923530103337     4.5786318685669      2.86467444773236e-06
--- gene_id: ENSMUSG00000002944 gene_name: Cd36 gene_location: 5:17986688-18093799 ---
Set                            LogFC                 LogCPM               P-val
T_Cells                        -6.93953962513339     4.58992017895346     0.00287725441857
--- gene_id: ENSMUSG00000009248 gene_name: Ascl2 gene_location: 7:142520566-142523001 ---
Set                            LogFC                 LogCPM               P-val
Lower_Hair_Follicle            -8.0735616731607      3.67305621711172     0.00707562212721825
--- gene_id: ENSMUSG00000026073 gene_name: Il1r2 gene_location: 1:40113239-40164391 ---
Set                            LogFC                 LogCPM               P-val
Mixed_Cluster                  -9.24523988778384     7.92082198129981     0.0065811096573842
--- gene_id: ENSMUSG00000025812 gene_name: Pard3 gene_location: 8:127790643-128338767 ---
Set                            LogFC                 LogCPM               P-val
T_Cells                        -5.34293606545251     2.35991899640858     0.00320147436747132
--- gene_id: ENSMUSG00000062515 gene_name: Fabp4 gene_location: 3:10269148-10273636 ---
Set                            LogFC                 LogCPM               P-val
T_Cells                        -7.73178477926329     5.78617666024203     0.00305158117397006
--- gene_id: ENSMUSG00000042250 gene_name: Pglyrp4 gene_location: 3:90634213-90648824 ---
Set                            LogFC                 LogCPM               P-val
Upper_Hair_Follicle            -7.28522157400808     4.53685817318403     0.00207819974486132
--- gene_id: ENSMUSG00000037868 gene_name: Egr2 gene_location: 10:67371305-67378018 ---
Set                            LogFC                 LogCPM               P-val
```

```
T_Cells                       -3.7553139166333     2.99104027113902    0.000892196031139953
--- gene_id: ENSMUSG00000036594 gene_name: H2-Aa gene_location: 17:34501718-34506797 ---
Set                           LogFC                LogCPM              P-val
Upper_Hair_Follicle           -3.94138307919221    5.19613482835959    0.00760042535172256
--- gene_id: ENSMUSG00000050953 gene_name: Gja1 gene_location: 10:56253426-56278609 ---
Set                           LogFC                LogCPM              P-val
T_Cells                       -5.17674110600648    3.47868981613699    0.00667305182491627
--- gene_id: ENSMUSG00000034201 gene_name: Gas2l1 gene_location: 11:5004132-5015327 ---
Set                           LogFC                LogCPM              P-val
T_Cells                       -5.76329535019131    3.08237262542188    0.00470101880546735
--- gene_id: ENSMUSG00000034634 gene_name: Ly6d gene_location: 15:74633905-74635469 ---
Set                           LogFC                LogCPM              P-val
T_Cells                       -2.59089980251959    5.3867504845131     0.00320147436747132
--- gene_id: ENSMUSG00000053137 gene_name: Mapk11 gene_location: 15:89026689-89033831 ---
Set                           LogFC                LogCPM              P-val
Upper_Hair_Follicle           -9.14080957566342    4.89148993721974    0.00190609679133284
--- gene_id: ENSMUSG00000020717 gene_name: Pecam1 gene_location: 11:106545043-106641454 ---
Set                           LogFC                LogCPM              P-val
Upper_Hair_Follicle           -9.26366255875141    5.04082177800958    0.00580907468179895
--- gene_id: ENSMUSG00000056708 gene_name: Ier5 gene_location: 1:154972107-154975382 ---
Set                           LogFC                LogCPM              P-val
Upper_Hair_Follicle           -6.0466211001329     2.75549478880975    0.00190609679133284
--- gene_id: ENSMUSG00000048756 gene_name: Foxo3 gene_location: 10:42057837-42152751 ---
Set                           LogFC                LogCPM              P-val
T_Cells                       5.75715614359508     4.26237003624113    0.00769973449418461
--- gene_id: ENSMUSG00000034845 gene_name: Plvap gene_location: 8:71950409-71964396 ---
Set                           LogFC                LogCPM              P-val
T_Cells                       -6.69417138547443    4.63949048263344    2.14996848196659e-05
--- gene_id: ENSMUSG00000029177 gene_name: Cenpa gene_location: 5:30824121-30832174 ---
Set                           LogFC                LogCPM              P-val
Upper_Hair_Follicle           -7.29220845547293    3.94203124378919    0.00375034200000489
--- gene_id: ENSMUSG00000028780 gene_name: Sema3c gene_location: 5:17779279-17935266 ---
Set                           LogFC                LogCPM              P-val
T_Cells                       -3.79193884732003    2.81948286629385    0.00434241283203604
--- gene_id: ENSMUSG00000044337 gene_name: Ackr3 gene_location: 1:90131702-90144473 ---
Set                           LogFC                LogCPM              P-val
T_Cells                       -3.17574872871855    1.9110432628978     0.00284526573091308
--- gene_id: ENSMUSG00000038059 gene_name: Smim3 gene_location: 18:60607263-60635059 ---
Set                           LogFC                LogCPM              P-val
Lower_Hair_Follicle           -6.96151713331908    3.91283539623861    0.00580798311751291
--- gene_id: ENSMUSG00000019838 gene_name: Slc16a10 gene_location: 10:39909528-40018254 ---
Set                           LogFC                LogCPM              P-val
Interfollicular_Epidermis     7.28798438026964     4.63439151949643    0.000574687955473636
--- gene_id: ENSMUSG00000022270 gene_name: Retreg1 gene_location: 15:25843266-25973773 ---
Set                           LogFC                LogCPM              P-val
T_Cells                       -3.87397715642882    2.1227438299805     0.00321714219669433
```

```
--- gene_id: ENSMUSG00000031078 gene_name: Cttn gene_location: 7:143989470-144024746 ---
Set                          LogFC                 LogCPM              P-val
T_Cells                      -5.51896437695824     2.9989238693753     0.00689813563966043
--- gene_id: ENSMUSG00000036887 gene_name: C1qa gene_location: 4:136623228-136626114 ---
Set                          LogFC                 LogCPM              P-val
Macrophages                  -12.5763298911542     6.97744795631801    6.78307685804257e-05
--- gene_id: ENSMUSG00000030727 gene_name: Rabep2 gene_location: 7:126027931-126048417 ---
Set                          LogFC                 LogCPM              P-val
T_Cells                      7.50540995931222      5.09032960322609    0.00396444388666427
--- gene_id: ENSMUSG00000032193 gene_name: Ldlr gene_location: 9:21634779-21661215 ---
Set                          LogFC                 LogCPM              P-val
T_Cells                      -5.64469762367921     3.18385482251164    0.00984198033294362
--- gene_id: ENSMUSG00000024436 gene_name: Mrps18b gene_location: 17:36221271-36227281 ---
Set                          LogFC                 LogCPM              P-val
T_Cells                      -5.66448074691109     2.82790459484294    0.00506351524109112
--- gene_id: ENSMUSG00000021950 gene_name: Anxa8 gene_location: 14:33807938-33822528 ---
Set                          LogFC                 LogCPM              P-val
T_Cells                      -2.55516787076009     5.07420954167392    0.000202802264042289
--- gene_id: ENSMUSG00000035372 gene_name: 1810055G02Rik gene_location: 19:3758293-3767881 ---
Set                          LogFC                 LogCPM              P-val
Upper_Hair_Follicle          -5.41837382574484     1.72084952847996    0.00718971603445386
--- gene_id: ENSMUSG00000002980 gene_name: Bcam gene_location: 7:19490056-19504941 ---
Set                          LogFC                 LogCPM              P-val
T_Cells                      -3.08843029489382     3.12198287603867    0.00220643616803176
--- gene_id: ENSMUSG00000050721 gene_name: Plekho2 gene_location: 9:65459980-65487322 ---
Set                          LogFC                 LogCPM              P-val
Lower_Hair_Follicle          -4.66241830353755     2.38358818410781    0.00707562212721825
--- gene_id: ENSMUSG00000025366 gene_name: Esyt1 gene_location: 10:128345834-128361740 ---
Set                          LogFC                 LogCPM              P-val
Interfollicular_Epidermis    -5.97906647725947     3.74576716114019    0.000570520249335575
--- gene_id: ENSMUSG00000053279 gene_name: Aldh1a1 gene_location: 19:20470079-20620829 ---
Set                          LogFC                 LogCPM              P-val
Mixed_Cluster                -10.5294597856312     6.60727250053895    0.00733415238455528
--- gene_id: ENSMUSG00000036381 gene_name: P2ry14 gene_location: 3:59021276-59061039 ---
Set                          LogFC                 LogCPM              P-val
T_Cells                      -6.96577794629049     3.40692858700471    0.00580754947327227
--- gene_id: ENSMUSG00000021186 gene_name: Fbln5 gene_location: 12:101712824-101785314 ---
Set                          LogFC                 LogCPM              P-val
T_Cells                      -4.95465929840301     2.9543329492431     0.00434241283203604
--- gene_id: ENSMUSG00000067006 gene_name: Serpinb5 gene_location: 1:106788903-106811078 ---
Set                          LogFC                 LogCPM              P-val
T_Cells                      -3.88555281224391     3.27233494952551    7.3693504909325e-05
--- gene_id: ENSMUSG00000046562 gene_name: Unc119b gene_location: 5:115260609-115273034 ---
Set                          LogFC                 LogCPM              P-val
T_Cells                      -6.52402562171845     3.96070323723816    0.00105477575627641
--- gene_id: ENSMUSG00000039686 gene_name: Zer1 gene_location: 2:29987295-30014597 ---
```

Set LogFC LogCPM P-val
Lower_Hair_Follicle -8.30023996769578 3.4054162619801 0.000637551509486392
--- gene_id: ENSMUSG00000030605 gene_name: Mfge8 gene_location: 7:78783516-78798808 ---
Set LogFC LogCPM P-val
Lower_Hair_Follicle -5.86189182100693 4.01345789547746 1.90266233650814e-06
--- gene_id: ENSMUSG00000024792 gene_name: Zfpl1 gene_location: 19:6130792-6134986 ---
Set LogFC LogCPM P-val
T_Cells 6.3784111271812 4.35119824084731 0.00580754947327227
--- gene_id: ENSMUSG00000031834 gene_name: Pik3r2 gene_location: 8:71220820-71229357 ---
Set LogFC LogCPM P-val
T_Cells 4.39227832766106 4.08661482425869 0.00431800642160075
--- gene_id: ENSMUSG00000042745 gene_name: Id1 gene_location: 2:152578171-152579330 ---
Set LogFC LogCPM P-val
T_Cells -4.76010211049417 3.15209590602989 0.00223537304195952
--- gene_id: ENSMUSG00000024907 gene_name: Gal gene_location: 19:3459915-3464544 ---
Set LogFC LogCPM P-val
Mixed_Cluster -11.6572819538951 7.71559091544276 0.00113689005914623
--- gene_id: ENSMUSG00000029417 gene_name: Cxcl9 gene_location: 5:92469206-92475938 ---
Set LogFC LogCPM P-val
Upper_Hair_Follicle -11.4909380649712 6.85813632784398 3.3520046329851e-07
--- gene_id: ENSMUSG00000024806 gene_name: Mlana gene_location: 19:29675224-29686034 ---
Set LogFC LogCPM P-val
Mixed_Cluster 11.107884855041 7.54543707892924 0.00733415238455528
--- gene_id: ENSMUSG00000027999 gene_name: Pla2g12a gene_location: 3:129672255-129689474 ---
Set LogFC LogCPM P-val
T_Cells 6.73984728924272 3.92329391488785 0.00886533612706817
--- gene_id: ENSMUSG00000036880 gene_name: Acaa2 gene_location: 18:74912268-74939279 ---
Set LogFC LogCPM P-val
T_Cells -3.2865744644886 2.93659796385074 0.00223537304195952
--- gene_id: ENSMUSG00000026342 gene_name: Slc35f5 gene_location: 1:125488332-125523557 ---
Set LogFC LogCPM P-val
T_Cells -7.05110169186788 3.38249570997854 8.82803175866674e-05
--- gene_id: ENSMUSG00000021710 gene_name: Nln gene_location: 13:104159565-104246122 ---
Set LogFC LogCPM P-val
T_Cells -7.31409085336741 3.73014021737536 0.00223537304195952
--- gene_id: ENSMUSG00000023043 gene_name: Krt18 gene_location: 15:101936615-101940462 ---
Set LogFC LogCPM P-val
Mixed_Cluster 12.2257016601014 8.6536728821683 0.0020312286061766
--- gene_id: ENSMUSG00000039521 gene_name: Foxp3 gene_location: X:7439883-7461484 ---
Set LogFC LogCPM P-val
Upper_Hair_Follicle -10.4910878642095 5.01268115878479 0.00718971603445386
--- gene_id: ENSMUSG00000058254 gene_name: Tspan7 gene_location: X:10351397-10462844 ---
Set LogFC LogCPM P-val
T_Cells 7.32238661296555 5.18484381652838 0.000439783859834232
--- gene_id: ENSMUSG00000040212 gene_name: Emp3 gene_location: 7:45567447-45570828 ---
Set LogFC LogCPM P-val

```
Upper_Hair_Follicle          -6.33423166571921    4.43201119129873    6.91490166228376e-07
--- gene_id: ENSMUSG00000066440 gene_name: Zfyve26 gene_location: 12:79279120-79343078 ---
Set                          LogFC                LogCPM              P-val
T_Cells                      7.89889274338848     4.08512834302841    0.00535554501396014
--- gene_id: ENSMUSG00000026213 gene_name: Stk11ip gene_location: 1:75498173-75513979 ---
Set                          LogFC                LogCPM              P-val
Mixed_Cluster                11.6656733871289     8.09747411384016    0.00385643389064778
--- gene_id: ENSMUSG00000033508 gene_name: Asprv1 gene_location: 6:86605146-86606692 ---
Set                          LogFC                LogCPM              P-val
Upper_Hair_Follicle          -9.27494698858443    4.69954897185987    0.000960118796980849
--- gene_id: ENSMUSG00000045545 gene_name: Krt14 gene_location: 11:100093988-100098374 ---
Set                          LogFC                LogCPM              P-val
T_Cells                      -2.16716295600045    6.85594679195778    0.0022488407656051
--- gene_id: ENSMUSG00000060591 gene_name: Ifitm2 gene_location: 7:140534750-140535900 ---
Set                          LogFC                LogCPM              P-val
T_Cells                      -4.52881731593692    3.84810824315408    1.84661201406147e-05
--- gene_id: ENSMUSG00000029504 gene_name: Ddx51 gene_location: 5:110801317-110808362 ---
Set                          LogFC                LogCPM              P-val
T_Cells                      -7.14530472639656    3.28161855293575    0.00273663070274176
--- gene_id: ENSMUSG00000015354 gene_name: Pcolce2 gene_location: 9:95519654-95580149 ---
Set                          LogFC                LogCPM              P-val
Lower_Hair_Follicle          -4.8898941540738     3.1851543276874     0.00103962849089551
--- gene_id: ENSMUSG00000039145 gene_name: Camk1d gene_location: 2:5298268-5719326 ---
Set                          LogFC                LogCPM              P-val
T_Cells                      -5.33521567206538    3.02925485142243    0.00553914193853242
--- gene_id: ENSMUSG00000004929 gene_name: Thop1 gene_location: 10:80905869-80918393 ---
Set                          LogFC                LogCPM              P-val
T_Cells                      -7.0364341051517     4.1200690786499     0.00670740752629547
--- gene_id: ENSMUSG00000008601 gene_name: Rab25 gene_location: 3:88449336-88455607 ---
Set                          LogFC                LogCPM              P-val
T_Cells                      -5.19897783996499    3.80487431568473    0.000403511848310349
--- gene_id: ENSMUSG00000008575 gene_name: Nfib gene_location: 4:82208410-82623987 ---
Set                          LogFC                LogCPM              P-val
T_Cells                      -5.98280554950334    4.39927937588305    1.84661201406147e-05
--- gene_id: ENSMUSG00000025150 gene_name: Cbr2 gene_location: 11:120620315-120622940 ---
Set                          LogFC                LogCPM              P-val
T_Cells                      -4.30483357096635    3.54451668676417    0.000300954737527096
--- gene_id: ENSMUSG00000019851 gene_name: Perp gene_location: 10:18720768-18732821 ---
Set                          LogFC                LogCPM              P-val
T_Cells                      -1.91840726168805    4.6117094797691     0.00805761779278866
--- gene_id: ENSMUSG00000016283 gene_name: H2-M2 gene_location: 17:37791742-37794443 ---
Set                          LogFC                LogCPM              P-val
Upper_Hair_Follicle          -7.73713531303024    3.86992589610659    0.00580907468179895
--- gene_id: ENSMUSG00000029309 gene_name: Sparcl1 gene_location: 5:104226977-104261599 ---
Set                          LogFC                LogCPM              P-val
T_Cells                      -5.8538271047103     4.20767040850698    0.000171206146252493
```

```
--- gene_id: ENSMUSG00000042271 gene_name: Nxt2 gene_location: X:141009766-141022688 ---
Set                          LogFC                LogCPM              P-val
T_Cells                      -6.27063066133252    2.50995667261945    0.00553914193853242
--- gene_id: ENSMUSG00000051329 gene_name: Nup160 gene_location: 2:90507559-90566672 ---
Set                          LogFC                LogCPM              P-val
T_Cells                      -7.0046828352521     3.16187540596527    0.00535991068124455
--- gene_id: ENSMUSG00000020083 gene_name: Fam241b gene_location: 10:61943434-61979699 ---
Set                          LogFC                LogCPM              P-val
T_Cells                      -5.94788149921792    2.9639791739261     0.00470101880546735
--- gene_id: ENSMUSG00000015843 gene_name: Rxrg gene_location: 1:167425953-167467192 ---
Set                          LogFC                LogCPM              P-val
Mixed_Cluster                -10.7820392969617    6.85413514750939    0.00636243947550316
--- gene_id: ENSMUSG00000044199 gene_name: S1pr4 gene_location: 10:81333581-81335966 ---
Set                          LogFC                LogCPM              P-val
Upper_Hair_Follicle          -10.428470439233     4.9519263082933     0.00830727059291126
--- gene_id: ENSMUSG00000054404 gene_name: Slfn5 gene_location: 11:82842175-82855666 ---
Set                          LogFC                LogCPM              P-val
T_Cells                      -4.27767078913618    2.39133591453633    0.00415616482663946
--- gene_id: ENSMUSG00000061740 gene_name: Cyp2d22 gene_location: 15:82254728-82264461 ---
Set                          LogFC                LogCPM              P-val
T_Cells                      -6.50783942792866    3.29722117495423    0.00396444388666427
--- gene_id: ENSMUSG00000000739 gene_name: Sult5a1 gene_location: 8:123866931-123885054 ---
Set                          LogFC                LogCPM              P-val
T_Cells                      -2.70425769529185    3.36086426681984    0.00094972774115982
--- gene_id: ENSMUSG00000063450 gene_name: Syne2 gene_location: 12:75864908-76157700 ---
Set                          LogFC                LogCPM              P-val
T_Cells                      -5.10461906179975    3.61452527540908    0.00037397531130852
--- gene_id: ENSMUSG00000025512 gene_name: Chid1 gene_location: 7:141073049-141119770 ---
Set                          LogFC                LogCPM              P-val
T_Cells                      -5.66414077775387    2.600398408994      0.00273663070274176
--- gene_id: ENSMUSG00000002897 gene_name: Il17ra gene_location: 6:120440208-120464520 ---
Set                          LogFC                LogCPM              P-val
Interfollicular_Epidermis    -7.69742857962667    3.30126239693874    0.00903315399425511
--- gene_id: ENSMUSG00000040331 gene_name: Nsmce4a gene_location: 7:130134256-130174848 ---
Set                          LogFC                LogCPM              P-val
T_Cells                      -7.21487439158863    4.2547117319689     0.000765035700204082
--- gene_id: ENSMUSG00000061527 gene_name: Krt5 gene_location: 15:101615505-101621333 ---
Set                          LogFC                LogCPM              P-val
T_Cells                      -2.14546861804051    5.63032577309702    0.00223537304195952
--- gene_id: ENSMUSG00000034473 gene_name: Sec22a gene_location: 16:35131501-35184288 ---
Set                          LogFC                LogCPM              P-val
T_Cells                      -7.25233522585404    3.85189649865496    0.00491662815936797
--- gene_id: ENSMUSG00000039197 gene_name: Adk gene_location: 14:21102642-21498637 ---
Set                          LogFC                LogCPM              P-val
T_Cells                      4.54055942762685     3.35295313953179    0.0099743994514158
--- gene_id: ENSMUSG00000024896 gene_name: Minpp1 gene_location: 19:32463169-32492764 ---
```

```
Set                          LogFC                 LogCPM              P-val
T_Cells                      -6.9850388480003      3.03520165289732    0.00220643616803176
--- gene_id: ENSMUSG00000036169 gene_name: Sostdc1 gene_location: 12:36364138-36368451 ---
Set                          LogFC                 LogCPM              P-val
Interfollicular_Epidermis    5.51902374466285      6.24297596699878    2.49947034184867e-05
--- gene_id: ENSMUSG00000026208 gene_name: Des gene_location: 1:75336973-75345223 ---
Set                          LogFC                 LogCPM              P-val
Mixed_Cluster                -11.3648417921133     8.2043762509058     0.00113689005914623
--- gene_id: ENSMUSG00000017607 gene_name: Tns4 gene_location: 11:98956504-98980132 ---
Set                          LogFC                 LogCPM              P-val
T_Cells                      -4.83355041582004     3.27357324123791    0.00670740752629547
--- gene_id: ENSMUSG00000029338 gene_name: Antxr2 gene_location: 5:98030642-98178902 ---
Set                          LogFC                 LogCPM              P-val
T_Cells                      -5.81724880030316     3.07713894120401    0.00535554501396014
--- gene_id: ENSMUSG00000006519 gene_name: Cyba gene_location: 8:123151515-123159669 ---
Set                          LogFC                 LogCPM              P-val
Upper_Hair_Follicle          -5.40714275085145     3.52845327527975    0.000854523039260141
--- gene_id: ENSMUSG00000044017 gene_name: Adgrd1 gene_location: 5:129173814-129281663 ---
Set                          LogFC                 LogCPM              P-val
T_Cells                      -5.96684636009268     3.99204327775221    0.000204396891821324
--- gene_id: ENSMUSG00000074457 gene_name: S100a16 gene_location: 3:90444561-90450458 ---
Set                          LogFC                 LogCPM              P-val
T_Cells                      -3.61784325826456     3.98421993436422    3.02698619413588e-05
--- gene_id: ENSMUSG00000030156 gene_name: Cd69 gene_location: 6:129244288-129252399 ---
Set                          LogFC                 LogCPM              P-val
Upper_Hair_Follicle          -6.70806677510611     4.14083625071253    0.00248999293340354
--- gene_id: ENSMUSG00000118087 gene_name: 4833438C02Rik gene_location: 19:46292020-46294075 ---
Set                          LogFC                 LogCPM              P-val
Upper_Hair_Follicle          -8.23130060591946     2.95641637683675    0.00760042535172256
--- gene_id: ENSMUSG00000001285 gene_name: Myg1 gene_location: 15:102240144-102246574 ---
Set                          LogFC                 LogCPM              P-val
T_Cells                      -6.54895822041731     4.22489791113057    0.00188509981786506
--- gene_id: ENSMUSG00000002602 gene_name: Axl gene_location: 7:25456698-25488130 ---
Set                          LogFC                 LogCPM              P-val
T_Cells                      -3.84934134913833     2.7751493145445     0.000145786878534649
--- gene_id: ENSMUSG00000060586 gene_name: H2-Eb1 gene_location: 17:34524841-34535648 ---
Set                          LogFC                 LogCPM              P-val
Upper_Hair_Follicle          -4.10162996120618     4.19619849494462    0.00197825453944371
--- gene_id: ENSMUSG00000054252 gene_name: Fgfr3 gene_location: 5:33879018-33894412 ---
Set                          LogFC                 LogCPM              P-val
T_Cells                      -4.80003206859802     2.55517998842145    0.00434241283203604
--- gene_id: ENSMUSG00000016382 gene_name: Pls3 gene_location: X:74829260-74918788 ---
Set                          LogFC                 LogCPM              P-val
T_Cells                      -6.78143061132095     5.06397386141951    0.000112366462945766
--- gene_id: ENSMUSG00000022562 gene_name: Oplah gene_location: 15:76180801-76212215 ---
Set                          LogFC                 LogCPM              P-val
```

```
T_Cells                     -4.76963177026364    4.27909583506386    2.14996848196659e-05
--- gene_id: ENSMUSG00000116504 gene_name: I730030J21Rik gene_location: 15:100628349-100630630 ---
Set                         LogFC                LogCPM              P-val
Upper_Hair_Follicle         -10.0936340583415    4.83358146597161    0.00951291899025052
--- gene_id: ENSMUSG00000031838 gene_name: Ifi30 gene_location: 8:71215419-71219307 ---
Set                         LogFC                LogCPM              P-val
T_Cells                     3.85292402948464     3.49515020444635    0.00284526573091308
--- gene_id: ENSMUSG00000025218 gene_name: Poll gene_location: 19:45540714-45548970 ---
Set                         LogFC                LogCPM              P-val
T_Cells                     -6.96844167686104    3.55935043694902    0.00635371181808078
--- gene_id: ENSMUSG00000089996 gene_name: Tmsb15b2 gene_location: X:135856014-135858774 ---
Set                         LogFC                LogCPM              P-val
Interfollicular_Epidermis   5.3401938724622      1.6656724435773     0.00215740439409542
--- gene_id: ENSMUSG00000025145 gene_name: Lrrc45 gene_location: 11:120604751-120611954 ---
Set                         LogFC                LogCPM              P-val
T_Cells                     -6.06651573414914    3.43611743454098    0.00553914193853242
--- gene_id: ENSMUSG00000079018 gene_name: Ly6c1 gene_location: 15:74915867-74920679 ---
Set                         LogFC                LogCPM              P-val
T_Cells                     -6.89630130498954    4.96815249227005    3.02698619413588e-05
--- gene_id: ENSMUSG00000038279 gene_name: Nop2 gene_location: 6:125108872-125121716 ---
Set                         LogFC                LogCPM              P-val
T_Cells                     -7.11462349453647    4.1666416931956     0.0085896489673039
--- gene_id: ENSMUSG00000031816 gene_name: Mthfsd gene_location: 8:121818367-121835131 ---
Set                         LogFC                LogCPM              P-val
Lower_Hair_Follicle         -6.97914091256748    3.00185689255761    0.00103962849089551
--- gene_id: ENSMUSG00000052031 gene_name: Tagap1 gene_location: 17:7222410-7228555 ---
Set                         LogFC                LogCPM              P-val
T_Cells                     -5.41015659859404    2.22850084526306    0.0085896489673039
--- gene_id: ENSMUSG00000030732 gene_name: Chrdl2 gene_location: 7:99655379-99683935 ---
Set                         LogFC                LogCPM              P-val
Mixed_Cluster               -11.7398317427097    7.79726207008379    0.00113689005914623
--- gene_id: ENSMUSG00000032068 gene_name: Plet1 gene_location: 9:50405825-50416782 ---
Set                         LogFC                LogCPM              P-val
Mixed_Cluster               9.64873995287722     9.27004837113207    0.00733415238455528
--- gene_id: ENSMUSG00000047501 gene_name: Cldn4 gene_location: 5:134973973-134975788 ---
Set                         LogFC                LogCPM              P-val
Mixed_Cluster               12.5999350311977     9.63377414387313    0.00113689005914623
--- gene_id: ENSMUSG00000031137 gene_name: Fgf13 gene_location: X:58107505-58613431 ---
Set                         LogFC                LogCPM              P-val
T_Cells                     -7.0208369591969     3.2102959246833     0.00573991064209263
--- gene_id: ENSMUSG00000032719 gene_name: Sbspon gene_location: 1:15924086-15962946 ---
Set                         LogFC                LogCPM              P-val
Mixed_Cluster               -10.6578946442006    6.73272463769593    0.00636243947550316
--- gene_id: ENSMUSG00000059898 gene_name: Dsc3 gene_location: 18:20093987-20135408 ---
Set                         LogFC                LogCPM              P-val
T_Cells                     -4.35251793037509    3.73091390087134    0.000112366462945766
```

```
--- gene_id: ENSMUSG00000008373 gene_name: Prpf31 gene_location: 7:3632984-3645485 ---
Set                          LogFC                 LogCPM              P-val
Upper_Hair_Follicle          -6.00230417974332     2.47073244029142    0.00548035665553508
--- gene_id: ENSMUSG00000053522 gene_name: Lgals7 gene_location: 7:28563278-28565709 ---
Set                          LogFC                 LogCPM              P-val
T_Cells                      -3.11826818470116     5.4078252821101     0.000172601698813066
--- gene_id: ENSMUSG00000022414 gene_name: Tab1 gene_location: 15:80017328-80045908 ---
Set                          LogFC                 LogCPM              P-val
T_Cells                      7.13364885014578      4.14418348414916    0.00995169199349372
--- gene_id: ENSMUSG00000034855 gene_name: Cxcl10 gene_location: 5:92494497-92496748 ---
Set                          LogFC                 LogCPM              P-val
Mixed_Cluster                -10.2273188436034     7.58248969912311    0.00636243947550316
--- gene_id: ENSMUSG00000021876 gene_name: Rnase4 gene_location: 14:51328534-51343608 ---
Set                          LogFC                 LogCPM              P-val
T_Cells                      -4.49923494056607     4.76978298916255    4.31302793881498e-08
--- gene_id: ENSMUSG00000031304 gene_name: Il2rg gene_location: X:100307984-100311861 ---
Set                          LogFC                 LogCPM              P-val
Upper_Hair_Follicle          -6.18943658484635     5.42737997554055    1.75939701258799e-08
--- gene_id: ENSMUSG00000066148 gene_name: Prpf4 gene_location: 4:62327034-62345227 ---
Set                          LogFC                 LogCPM              P-val
T_Cells                      -6.76403698272727     3.45261427362871    0.00390702964183957
--- gene_id: ENSMUSG00000054293 gene_name: P2ry10b gene_location: X:106192533-106217267 ---
Set                          LogFC                 LogCPM              P-val
T_Cells                      4.11437534780873      1.83679464179904    0.00320147436747132
--- gene_id: ENSMUSG00000032180 gene_name: Tmed1 gene_location: 9:21418849-21421548 ---
Set                          LogFC                 LogCPM              P-val
T_Cells                      5.69030964200577      4.05957765205708    0.00409426293677322
--- gene_id: ENSMUSG00000027962 gene_name: Vcam1 gene_location: 3:115903598-115923337 ---
Set                          LogFC                 LogCPM              P-val
Mixed_Cluster                -10.1059707949847     7.21929816745812    0.00636243947550316
--- gene_id: ENSMUSG00000033253 gene_name: Szt2 gene_location: 4:118219940-118266470 ---
Set                          LogFC                 LogCPM              P-val
T_Cells                      8.01468660300831      4.18874067681193    0.00507793256535871
--- gene_id: ENSMUSG00000020241 gene_name: Col6a2 gene_location: 10:76431596-76459464 ---
Set                          LogFC                 LogCPM              P-val
T_Cells                      -4.43302476830022     2.73244106251605    0.00188509981786506
--- gene_id: ENSMUSG00000023571 gene_name: C1qtnf12 gene_location: 4:156046775-156051086 ---
Set                          LogFC                 LogCPM              P-val
T_Cells                      -3.39996440479255     3.312491203244      0.000505099190929517
--- gene_id: ENSMUSG00000066026 gene_name: Dhrs3 gene_location: 4:144619397-144654779 ---
Set                          LogFC                 LogCPM              P-val
T_Cells                      -4.75018385428031     3.19110942482708    0.00810879239872489
--- gene_id: ENSMUSG00000020844 gene_name: Nxn gene_location: 11:76148024-76289966 ---
Set                          LogFC                 LogCPM              P-val
T_Cells                      -4.9672414258943      3.10760655461496    0.00888140771656999
--- gene_id: ENSMUSG00000017466 gene_name: Timp2 gene_location: 11:118191887-118246566 ---
```

| Set | LogFC | LogCPM | P-val |
|---|---|---|---|
| T_Cells | -3.11336396540572 | 2.40943785747312 | 0.000984844988950389 |

--- gene_id: ENSMUSG00000040564 gene_name: Apoc1 gene_location: 7:19423406-19426585 ---

| Set | LogFC | LogCPM | P-val |
|---|---|---|---|
| T_Cells | -2.79070745412549 | 3.2649056330258 | 0.00273663070274176 |

--- gene_id: ENSMUSG00000005413 gene_name: Hmox1 gene_location: 8:75820249-75827217 ---

| Set | LogFC | LogCPM | P-val |
|---|---|---|---|
| T_Cells | -4.86789102088371 | 2.7536237421628 | 0.0057767422642682 |

--- gene_id: ENSMUSG00000001240 gene_name: Ramp2 gene_location: 11:101136854-101150372 ---

| Set | LogFC | LogCPM | P-val |
|---|---|---|---|
| T_Cells | -4.40943739777169 | 2.43264645173938 | 0.00321714219669433 |

--- gene_id: ENSMUSG00000020941 gene_name: Map3k14 gene_location: 11:103110588-103158298 ---

| Set | LogFC | LogCPM | P-val |
|---|---|---|---|
| Interfollicular_Epidermis | -7.39970613393895 | 2.55220313115913 | 0.00818214317425175 |

--- gene_id: ENSMUSG00000081769 gene_name: Gm12216 gene_location: 11:53674244-53750082 ---

| Set | LogFC | LogCPM | P-val |
|---|---|---|---|
| Upper_Hair_Follicle | -10.5714636057679 | 5.09204173716633 | 0.000523831735251934 |

--- gene_id: ENSMUSG00000058756 gene_name: Thra gene_location: 11:98631464-98659832 ---

| Set | LogFC | LogCPM | P-val |
|---|---|---|---|
| T_Cells | -5.58136969263394 | 2.78897277702339 | 0.00502474583660132 |

--- gene_id: ENSMUSG00000025473 gene_name: Adam8 gene_location: 7:139558845-139572475 ---

| Set | LogFC | LogCPM | P-val |
|---|---|---|---|
| Upper_Hair_Follicle | -7.96180288046102 | 4.34664174858769 | 0.00190609679133284 |

--- gene_id: ENSMUSG00000034783 gene_name: Cd207 gene_location: 6:83648197-83654839 ---

| Set | LogFC | LogCPM | P-val |
|---|---|---|---|
| Mixed_Cluster | -11.2441076993593 | 8.08510054804822 | 0.0020666444079557 |

--- gene_id: ENSMUSG00000022474 gene_name: Pmm1 gene_location: 15:81835309-81845131 ---

| Set | LogFC | LogCPM | P-val |
|---|---|---|---|
| T_Cells | -6.20393573491955 | 4.27584097287394 | 0.000113304361482114 |

--- gene_id: ENSMUSG00000018425 gene_name: Dhx40 gene_location: 11:86659672-86698572 ---

| Set | LogFC | LogCPM | P-val |
|---|---|---|---|
| Interfollicular_Epidermis | -6.2861967229875 | 2.82746145101967 | 0.00759656213586384 |

--- gene_id: ENSMUSG00000022037 gene_name: Clu gene_location: 14:66205932-66218996 ---

| Set | LogFC | LogCPM | P-val |
|---|---|---|---|
| T_Cells | -5.50313606081124 | 4.5612551379072 | 2.63624364573524e-06 |

--- gene_id: ENSMUSG00000027555 gene_name: Car13 gene_location: 3:14706787-14728062 ---

| Set | LogFC | LogCPM | P-val |
|---|---|---|---|
| T_Cells | -4.27478750697735 | 2.6182976797424 | 0.00248584639952393 |

--- gene_id: ENSMUSG00000036570 gene_name: Fxyd1 gene_location: 7:30751103-30756624 ---

| Set | LogFC | LogCPM | P-val |
|---|---|---|---|
| Mixed_Cluster | -9.36313204825313 | 7.73657681117416 | 0.00636243947550316 |

--- gene_id: ENSMUSG00000027800 gene_name: Tm4sf1 gene_location: 3:57193032-57209409 ---

| Set | LogFC | LogCPM | P-val |
|---|---|---|---|
| T_Cells | -8.13844455597713 | 6.21225783370694 | 1.84661201406147e-05 |

--- gene_id: ENSMUSG00000058914 gene_name: C1qtnf3 gene_location: 15:10952418-10980236 ---

| Set | LogFC | LogCPM | P-val |
|---|---|---|---|

```
Mixed_Cluster                -10.4286192354152    7.53898235352359    0.00485094649090404
--- gene_id: ENSMUSG00000007655 gene_name: Cav1 gene_location: 6:17306334-17341451 ---
Set                          LogFC                LogCPM              P-val
T_Cells                      -3.76684051546542    3.93309464442338    3.02698619413588e-05
--- gene_id: ENSMUSG00000030805 gene_name: Stx4a gene_location: 7:127423466-127448191 ---
Set                          LogFC                LogCPM              P-val
T_Cells                      -6.12607589648961    3.59540086874209    0.00434241283203604
--- gene_id: ENSMUSG00000026167 gene_name: Wnt10a gene_location: 1:74830675-74843338 ---
Set                          LogFC                LogCPM              P-val
T_Cells                      -3.82621248101254    2.46446694687397    0.000403511848310349
--- gene_id: ENSMUSG00000022661 gene_name: Cd200 gene_location: 16:45202498-45229416 ---
Set                          LogFC                LogCPM              P-val
T_Cells                      -3.23148485065781    2.2283453066284     0.0085896489673039
--- gene_id: ENSMUSG00000023908 gene_name: Pkmyt1 gene_location: 17:23945310-23955709 ---
Set                          LogFC                LogCPM              P-val
T_Cells                      -6.56179918421963    3.70313450229453    0.00396444388666427
--- gene_id: ENSMUSG00000049421 gene_name: Zfp260 gene_location: 7:29794202-29807047 ---
Set                          LogFC                LogCPM              P-val
Interfollicular_Epidermis    -5.60415974671341    2.89075542943857    0.00275517953788891
--- gene_id: ENSMUSG00000026509 gene_name: Capn2 gene_location: 1:182294825-182345173 ---
Set                          LogFC                LogCPM              P-val
Interfollicular_Epidermis    -7.10028836429716    3.69793794268896    0.00275517953788891
--- gene_id: ENSMUSG00000040723 gene_name: Rcsd1 gene_location: 1:165474085-165537326 ---
Set                          LogFC                LogCPM              P-val
Upper_Hair_Follicle          -8.95514442293489    5.10350075214759    0.00190609679133284
--- gene_id: ENSMUSG00000061353 gene_name: Cxcl12 gene_location: 6:117145496-117158328 ---
Set                          LogFC                LogCPM              P-val
T_Cells                      -5.76671840934043    4.98991109999481    1.80286561342313e-05
--- gene_id: ENSMUSG00000024610 gene_name: Cd74 gene_location: 18:60936920-60945724 ---
Set                          LogFC                LogCPM              P-val
Upper_Hair_Follicle          -3.84553030366661    6.36130124550291    0.00345394546539992
--- gene_id: ENSMUSG00000042286 gene_name: Stab1 gene_location: 14:30860970-30890598 ---
Set                          LogFC                LogCPM              P-val
Macrophages                  -12.4531514496104    6.85511525245544    6.90143015496872e-05
--- gene_id: ENSMUSG00000028494 gene_name: Plin2 gene_location: 4:86566623-86588297 ---
Set                          LogFC                LogCPM              P-val
T_Cells                      -4.05932284884203    3.2608611367198     0.00396444388666427
--- gene_id: ENSMUSG00000083367 gene_name: Gm8806 gene_location: X:79577351-79578757 ---
Set                          LogFC                LogCPM              P-val
Interfollicular_Epidermis    3.08727147740493     1.1632372723604     0.00525924324302293
--- gene_id: ENSMUSG00000070473 gene_name: Cldn3 gene_location: 5:135015068-135016326 ---
Set                          LogFC                LogCPM              P-val
Mixed_Cluster                10.8621444285369     7.30299126251542    0.00938932261884825
--- gene_id: ENSMUSG00000001323 gene_name: Srr gene_location: 11:74797185-74816774 ---
Set                          LogFC                LogCPM              P-val
T_Cells                      -5.85643852927219    2.49241867456735    0.00810879239872489
```

```
--- gene_id: ENSMUSG00000048332 gene_name: Lhfp gene_location: 3:52948949-53169100 ---
Set                          LogFC                 LogCPM              P-val
T_Cells                      7.31434145754034      4.27146687307873    0.00470101880546735
--- gene_id: ENSMUSG00000031207 gene_name: Msn gene_location: X:95139648-95212158 ---
Set                          LogFC                 LogCPM              P-val
Interfollicular_Epidermis    -4.26864325460883     4.41209909280081    0.00255220896026439
--- gene_id: ENSMUSG00000035783 gene_name: Acta2 gene_location: 19:34218490-34232990 ---
Set                          LogFC                 LogCPM              P-val
Mixed_Cluster                -9.42476634615917     9.30076501232727    0.0046277826535293
--- gene_id: ENSMUSG00000025158 gene_name: Rfng gene_location: 11:120671572-120675033 ---
Set                          LogFC                 LogCPM              P-val
T_Cells                      -6.68616340706524     3.58884580634277    0.00647194152102878
--- gene_id: ENSMUSG00000036256 gene_name: Igfbp7 gene_location: 5:77497087-77555888 ---
Set                          LogFC                 LogCPM              P-val
T_Cells                      -4.45223107574431     3.11270371479519    0.000300954737527096
--- gene_id: ENSMUSG00000054146 gene_name: Krt15 gene_location: 11:100022584-100026754 ---
Set                          LogFC                 LogCPM              P-val
T_Cells                      -2.03978095924208     6.51492476374021    0.00223537304195952
--- gene_id: ENSMUSG00000031841 gene_name: Cdh13 gene_location: 8:119010472-120051660 ---
Set                          LogFC                 LogCPM              P-val
T_Cells                      -3.35045881331732     3.05373657696852    0.000642118849834969
--- gene_id: ENSMUSG00000027303 gene_name: Ptpra gene_location: 2:130292198-130398044 ---
Set                          LogFC                 LogCPM              P-val
Interfollicular_Epidermis    -5.98864991724374     2.86956407322357    0.00525924324302293
--- gene_id: ENSMUSG00000021981 gene_name: Cab39l gene_location: 14:59678421-59823213 ---
Set                          LogFC                 LogCPM              P-val
T_Cells                      5.54992717228157      4.55509257924975    0.00223537304195952
--- gene_id: ENSMUSG00000022512 gene_name: Cldn1 gene_location: 16:26175392-26190591 ---
Set                          LogFC                 LogCPM              P-val
T_Cells                      -3.69815184451498     3.45585152564582    0.000192681630347959
--- gene_id: ENSMUSG00000104788 gene_name: Gm36448 gene_location: 14:52776109-52778333 ---
Set                          LogFC                 LogCPM              P-val
T_Cells                      5.01487678924219      1.93690705555212    1.86237346033295e-06
--- gene_id: ENSMUSG00000022672 gene_name: Prkdc gene_location: 16:15455730-15660099 ---
Set                          LogFC                 LogCPM              P-val
T_Cells                      -7.59072733126699     3.88572020956506    0.00689813563966043
```
