## Supplemental Table 3 for "Single-cell X-chromosome inactivation analysis links biased chimerism to differential gene expression and epigenetic erosion"

| Sample:003C | Disease:Control | | | Sex:F | | Age:67 | UMI1:3460 | UMI2:965 | total1:0 | total2:114 |
|---|---|---|---|---|---|---|---|---|---|---|
| Region_Celltype | X1 | X2 | Both | Unknown | LowCoverage | Expected1 | Expected2 | P_value | Direction | |
| Macrophage | 0 | 55 | 3 | 154 | 209 | 0 | 55 | 1 | NS | |
| Macrophage_Alveolar | 0 | 32 | 2 | 90 | 160 | 0 | 32 | 1 | NS | |
| cMonocyte | 0 | 2 | 1 | 34 | 42 | 0 | 2 | 1 | NS | |
| cDC2 | 0 | 14 | 1 | 12 | 33 | 0 | 14 | 1 | NS | |
| pDC | 0 | 0 | 0 | 26 | 1 | 0 | 0 | 1 | NS | |
| cDC1 | 0 | 3 | 1 | 7 | 5 | 0 | 3 | 1 | NS | |
| ncMonocyte | 0 | 0 | 0 | 4 | 8 | 0 | 0 | 1 | NS | |
| T_Cytotoxic | 0 | 2 | 0 | 2 | 8 | 0 | 2 | 1 | NS | |
| Multiplet | 0 | 2 | 0 | 2 | 3 | 0 | 2 | 1 | NS | |
| DC_Mature | 0 | 1 | 0 | 4 | 2 | 0 | 1 | 1 | NS | |
| T | 0 | 0 | 0 | 1 | 5 | 0 | 0 | 1 | NS | |
| B | 0 | 0 | 0 | 4 | 1 | 0 | 0 | 1 | NS | |
| Mast | 0 | 3 | 0 | 0 | 1 | 0 | 3 | 1 | NS | |
| VE_Capillary_B | 0 | 0 | 0 | 4 | 0 | 0 | 0 | 1 | NS | |
| NK | 0 | 0 | 0 | 2 | 2 | 0 | 0 | 1 | NS | |
| Ciliated | 0 | 0 | 0 | 2 | 1 | 0 | 0 | 1 | NS | |
| ILC_A | 0 | 0 | 0 | 0 | 2 | 0 | 0 | 1 | NS | |
| VE_Capillary_A | 0 | 0 | 0 | 0 | 2 | 0 | 0 | 1 | NS | |
| Lymphatic | 0 | 0 | 0 | 0 | 2 | 0 | 0 | 1 | NS | |
| B_Plasma | 0 | 0 | 0 | 0 | 2 | 0 | 0 | 1 | NS | |
| Club | 0 | 0 | 0 | 1 | 0 | 0 | 0 | 1 | NS | |
| ATI | 0 | 0 | 0 | 0 | 1 | 0 | 0 | 1 | NS | |
| ATII | 0 | 0 | 0 | 0 | 1 | 0 | 0 | 1 | NS | |
| DC_Langerhans | 0 | 0 | 0 | 0 | 1 | 0 | 0 | 1 | NS | |
| Fibroblast | 0 | 0 | 0 | 0 | 1 | 0 | 0 | 1 | NS | |
| VE_Arterial | 0 | 0 | 0 | 1 | 0 | 0 | 0 | 1 | NS | |

| Sample:052CO | Disease:COPD | | | Sex:F | Age:62 | UMI1:3482 | UMI2:2080 | total1:302 | total2:344 |
|---|---|---|---|---|---|---|---|---|---|
| Region_Celltype | X1 | X2 | Both | Unknown | LowCoverage | Expected1 | Expected2 | P_value | Direction |
| T_Cytotoxic | 26 | 73 | 12 | 215 | 217 | 46.282 | 52.718 | 0.0027552 | Outlier |
| T | 18 | 33 | 8 | 119 | 169 | 23.842 | 27.158 | 0.23958 | NS |
| cMonocyte | 72 | 56 | 3 | 15 | 72 | 59.839 | 68.161 | 0.12831 | NS |
| Macrophage_Alveolar | 39 | 45 | 2 | 34 | 57 | 39.269 | 44.731 | 0.96677 | NS |
| NK | 11 | 9 | 1 | 53 | 53 | 9.35 | 10.65 | 0.60174 | NS |
| ncMonocyte | 50 | 40 | 1 | 5 | 24 | 42.074 | 47.926 | 0.23728 | NS |
| Macrophage | 13 | 15 | 2 | 28 | 27 | 13.09 | 14.91 | 0.98081 | NS |
| Multiplet | 18 | 13 | 5 | 14 | 17 | 14.492 | 16.508 | 0.37239 | NS |
| cDC2 | 11 | 23 | 0 | 11 | 21 | 15.895 | 18.105 | 0.22477 | NS |
| ATII | 3 | 6 | 2 | 18 | 18 | 4.207 | 4.793 | 0.56136 | NS |
| ATI | 1 | 3 | 2 | 15 | 23 | 1.87 | 2.13 | 0.52134 | NS |
| B | 3 | 1 | 1 | 16 | 17 | 1.87 | 2.13 | 0.41299 | NS |

| | | | | | | | | | |
|---|---|---|---|---|---|---|---|---|---|
| VE_Capillary_B | 4 | 3 | 0 | 4 | 21 | 3.272 | 3.728 | 0.69714 | NS |
| Myofibroblast | 4 | 5 | 0 | 1 | 13 | 4.207 | 4.793 | 0.9218 | NS |
| Fibroblast | 7 | 10 | 0 | 1 | 2 | 7.947 | 9.053 | 0.74341 | NS |
| cDC1 | 5 | 0 | 2 | 5 | 5 | 2.337 | 2.663 | 0.056792 | NS |
| VE_Capillary_A | 2 | 1 | 0 | 4 | 10 | 1.402 | 1.598 | 0.62249 | NS |
| Lymphatic | 2 | 2 | 0 | 1 | 9 | 1.87 | 2.13 | 0.9267 | NS |
| B_Plasma | 1 | 1 | 0 | 2 | 10 | 0.935 | 1.065 | 0.94813 | NS |
| Mast | 1 | 1 | 0 | 0 | 7 | 0.935 | 1.065 | 0.94813 | NS |
| VE_Arterial | 2 | 2 | 0 | 1 | 3 | 1.87 | 2.13 | 0.9267 | NS |
| Ciliated | 0 | 0 | 0 | 6 | 1 | 0 | 0 | 1 | NS |
| T_Regulatory | 0 | 0 | 1 | 1 | 4 | 0 | 0 | 1 | NS |
| SMC | 0 | 0 | 0 | 0 | 6 | 0 | 0 | 1 | NS |
| Mesothelial | 5 | 0 | 0 | 0 | 1 | 2.337 | 2.663 | 0.056792 | NS |
| Pericyte | 2 | 0 | 0 | 0 | 2 | 0.935 | 1.065 | 0.22829 | NS |
| Club | 0 | 0 | 1 | 0 | 3 | 0 | 0 | 1 | NS |
| pDC | 0 | 0 | 0 | 1 | 2 | 0 | 0 | 1 | NS |
| ILC_A | 0 | 0 | 0 | 2 | 1 | 0 | 0 | 1 | NS |
| ILC_B | 0 | 0 | 0 | 1 | 1 | 0 | 0 | 1 | NS |
| Basal | 1 | 1 | 0 | 0 | 0 | 0.935 | 1.065 | 0.94813 | NS |
| Goblet | 0 | 1 | 0 | 0 | 1 | 0.467 | 0.533 | 0.43475 | NS |
| DC_Mature | 0 | 0 | 0 | 1 | 0 | 0 | 0 | 1 | NS |
| VE_Venous | 1 | 0 | 0 | 0 | 0 | 0.467 | 0.533 | 0.39427 | NS |

| Sample:056CO | Disease:COPD | | | Sex:F | Age:57 | UMI1:4382 | UMI2:2193 | total1:106 | total2:254 |
|---|---|---|---|---|---|---|---|---|---|
| Region_Celltype | X1 | X2 | Both | Unknown | LowCoverage | Expected1 | Expected2 | P_value | Direction |
| Macrophage_Alveolar | 35 | 76 | 8 | 361 | 196 | 32.683 | 78.317 | 0.73555 | NS |
| Macrophage | 23 | 44 | 10 | 104 | 116 | 19.728 | 47.272 | 0.54415 | NS |
| T | 12 | 6 | 2 | 146 | 75 | 5.3 | 12.7 | 0.025416 | NS |
| T_Cytotoxic | 9 | 7 | 2 | 119 | 65 | 4.711 | 11.289 | 0.1255 | NS |
| cMonocyte | 0 | 17 | 0 | 45 | 59 | 5.006 | 11.994 | 0.015404 | NS |
| ncMonocyte | 1 | 21 | 2 | 36 | 48 | 6.478 | 15.522 | 0.027899 | NS |
| NK | 2 | 8 | 0 | 57 | 26 | 2.944 | 7.056 | 0.62446 | NS |
| cDC2 | 10 | 19 | 4 | 25 | 23 | 8.539 | 20.461 | 0.68077 | NS |
| B | 2 | 9 | 1 | 33 | 14 | 3.239 | 7.761 | 0.53518 | NS |
| Multiplet | 4 | 9 | 1 | 24 | 15 | 3.828 | 9.172 | 0.94131 | NS |
| B_Plasma | 0 | 3 | 0 | 23 | 18 | 0.883 | 2.117 | 0.30879 | NS |
| Ciliated | 1 | 3 | 2 | 20 | 15 | 1.178 | 2.822 | 0.8877 | NS |
| ATII | 1 | 2 | 0 | 15 | 10 | 0.883 | 2.117 | 0.91825 | NS |
| DC_Mature | 2 | 5 | 4 | 7 | 3 | 2.061 | 4.939 | 0.97129 | NS |
| cDC1 | 1 | 7 | 0 | 8 | 3 | 2.356 | 5.644 | 0.40517 | NS |
| Mast | 0 | 1 | 0 | 9 | 6 | 0.294 | 0.706 | 0.5568 | NS |
| Myofibroblast | 2 | 4 | 0 | 3 | 6 | 1.767 | 4.233 | 0.8846 | NS |
| Lymphatic | 0 | 1 | 0 | 7 | 4 | 0.294 | 0.706 | 0.5568 | NS |

| | | | | | | | | | |
|---|---|---|---|---|---|---|---|---|---|
| pDC | 0 | 0 | 0 | 9 | 2 | 0 | 0 | 1 | NS |
| VE_Capillary_B | 0 | 0 | 0 | 5 | 5 | 0 | 0 | 1 | NS |
| Mesothelial | 0 | 5 | 0 | 3 | 1 | 1.472 | 3.528 | 0.18887 | NS |
| Fibroblast | 0 | 3 | 0 | 2 | 2 | 0.883 | 2.117 | 0.30879 | NS |
| Club | 0 | 1 | 0 | 5 | 1 | 0.294 | 0.706 | 0.5568 | NS |
| ILC_A | 1 | 0 | 0 | 2 | 3 | 0.294 | 0.706 | 0.29644 | NS |
| ATI | 0 | 1 | 0 | 2 | 1 | 0.294 | 0.706 | 0.5568 | NS |
| T_Regulatory | 0 | 1 | 0 | 2 | 0 | 0.294 | 0.706 | 0.5568 | NS |
| VE_Arterial | 0 | 0 | 0 | 3 | 0 | 0 | 0 | 1 | NS |
| Basal | 0 | 0 | 0 | 2 | 0 | 0 | 0 | 1 | NS |
| DC_Langerhans | 0 | 0 | 0 | 1 | 0 | 0 | 0 | 1 | NS |
| SMC | 0 | 0 | 0 | 0 | 1 | 0 | 0 | 1 | NS |
| VE_Venous | 0 | 1 | 0 | 0 | 0 | 0.294 | 0.706 | 0.5568 | NS |
| ILC_B | 0 | 0 | 0 | 1 | 0 | 0 | 0 | 1 | NS |

Sample:098C Disease:Control Sex:F Age:41 UMI1:9240 UMI2:5955 total1:1216 total2:2071

| Region_Celltype | X1 | X2 | Both | Unknown | LowCoverage | Expected1 | Expected2 | P_value | Direction |
|---|---|---|---|---|---|---|---|---|---|
| Macrophage_Alveolar | 911 | 1558 | 166 | 212 | 672 | 913.387 | 1555.613 | 0.94389 | NS |
| Macrophage | 235 | 375 | 49 | 114 | 286 | 225.665 | 384.335 | 0.58142 | NS |
| cMonocyte | 23 | 43 | 4 | 90 | 111 | 24.416 | 41.584 | 0.79724 | NS |
| T_Cytotoxic | 3 | 4 | 2 | 88 | 94 | 2.59 | 4.41 | 0.82279 | NS |
| cDC2 | 9 | 23 | 1 | 61 | 59 | 11.838 | 20.162 | 0.449 | NS |
| NK | 3 | 12 | 0 | 71 | 52 | 5.549 | 9.451 | 0.30253 | NS |
| Multiplet | 10 | 15 | 8 | 18 | 19 | 9.249 | 15.751 | 0.82712 | NS |
| T | 1 | 4 | 1 | 37 | 27 | 1.85 | 3.15 | 0.55167 | NS |
| Ciliated | 0 | 1 | 0 | 39 | 22 | 0.37 | 0.63 | 0.50049 | NS |
| Fibroblast | 3 | 3 | 1 | 34 | 18 | 2.22 | 3.78 | 0.64955 | NS |
| B | 3 | 1 | 4 | 30 | 21 | 1.48 | 2.52 | 0.2789 | NS |
| ncMonocyte | 7 | 11 | 2 | 17 | 20 | 6.659 | 11.341 | 0.90675 | NS |
| ATII | 2 | 4 | 1 | 17 | 25 | 2.22 | 3.78 | 0.89435 | NS |
| Lymphatic | 1 | 2 | 0 | 18 | 19 | 1.11 | 1.89 | 0.92519 | NS |
| T_Regulatory | 0 | 4 | 1 | 5 | 12 | 1.48 | 2.52 | 0.17784 | NS |
| VE_Arterial | 1 | 3 | 0 | 8 | 8 | 1.48 | 2.52 | 0.71379 | NS |
| Basal | 0 | 2 | 0 | 6 | 10 | 0.74 | 1.26 | 0.3407 | NS |
| cDC1 | 0 | 1 | 0 | 11 | 5 | 0.37 | 0.63 | 0.50049 | NS |
| ILC_A | 0 | 0 | 0 | 9 | 5 | 0 | 0 | 1 | NS |
| Goblet | 1 | 0 | 0 | 5 | 4 | 0.37 | 0.63 | 0.33752 | NS |
| DC_Mature | 0 | 1 | 0 | 5 | 4 | 0.37 | 0.63 | 0.50049 | NS |
| VE_Capillary_B | 0 | 1 | 0 | 4 | 3 | 0.37 | 0.63 | 0.50049 | NS |
| VE_Capillary_A | 0 | 0 | 0 | 5 | 1 | 0 | 0 | 1 | NS |
| ATI | 0 | 0 | 0 | 3 | 3 | 0 | 0 | 1 | NS |
| VE_Venous | 0 | 1 | 0 | 1 | 3 | 0.37 | 0.63 | 0.50049 | NS |
| Myofibroblast | 1 | 0 | 0 | 2 | 2 | 0.37 | 0.63 | 0.33752 | NS |

| | | | | | | | | | |
|---|---|---|---|---|---|---|---|---|---|
| Mesothelial | 0 | 1 | 0 | 2 | 0 | 0.37 | 0.63 | 0.50049 | NS |
| B_Plasma | 0 | 0 | 0 | 1 | 2 | 0 | 0 | 1 | NS |
| VE_Peribronchial | 1 | 0 | 0 | 1 | 1 | 0.37 | 0.63 | 0.33752 | NS |
| Mast | 0 | 0 | 0 | 1 | 1 | 0 | 0 | 1 | NS |
| ILC_B | 1 | 0 | 0 | 0 | 1 | 0.37 | 0.63 | 0.33752 | NS |
| PNEC | 0 | 0 | 0 | 1 | 0 | 0 | 0 | 1 | NS |
| SMC | 0 | 1 | 0 | 0 | 0 | 0.37 | 0.63 | 0.50049 | NS |
| DC_Langerhans | 0 | 0 | 0 | 0 | 1 | 0 | 0 | 1 | NS |
| Club | 0 | 0 | 0 | 1 | 0 | 0 | 0 | 1 | NS |

Sample:133C Disease:Control Sex:F Age:32 UMI1:8566 UMI2:6112 total1:1175 total2:2132

| Region_Celltype | X1 | X2 | Both | Unknown | LowCoverage | Expected1 | Expected2 | P_value | Direction |
|---|---|---|---|---|---|---|---|---|---|
| Macrophage | 566 | 981 | 66 | 316 | 1207 | 549.66 | 997.34 | 0.54068 | NS |
| ATII | 250 | 567 | 35 | 35 | 330 | 290.286 | 526.714 | 0.034139 | NS |
| Macrophage_Alveolar | 248 | 333 | 43 | 41 | 265 | 206.433 | 374.567 | 0.012462 | NS |
| cMonocyte | 35 | 78 | 6 | 15 | 92 | 40.15 | 72.85 | 0.46716 | NS |
| Multiplet | 28 | 59 | 41 | 10 | 41 | 30.912 | 56.088 | 0.64089 | NS |
| cDC2 | 26 | 38 | 1 | 18 | 25 | 22.74 | 41.26 | 0.55286 | NS |
| T_Cytotoxic | 4 | 6 | 1 | 12 | 38 | 3.553 | 6.447 | 0.83668 | NS |
| Ciliated | 2 | 27 | 2 | 3 | 20 | 10.304 | 18.696 | 0.0076515 | Outlier |
| ATI | 0 | 6 | 1 | 2 | 30 | 2.132 | 3.868 | 0.10738 | NS |
| T | 1 | 1 | 1 | 5 | 18 | 0.711 | 1.289 | 0.76993 | NS |
| NK | 0 | 2 | 0 | 2 | 15 | 0.711 | 1.289 | 0.35259 | NS |
| cDC1 | 2 | 4 | 0 | 3 | 3 | 2.132 | 3.868 | 0.93616 | NS |
| DC_Mature | 3 | 4 | 1 | 3 | 1 | 2.487 | 4.513 | 0.77889 | NS |
| VE_Venous | 1 | 2 | 0 | 0 | 7 | 1.066 | 1.934 | 0.95483 | NS |
| VE_Arterial | 1 | 2 | 0 | 2 | 5 | 1.066 | 1.934 | 0.95483 | NS |
| Goblet | 0 | 8 | 0 | 0 | 1 | 2.842 | 5.158 | 0.063003 | NS |
| B | 2 | 2 | 0 | 1 | 4 | 1.421 | 2.579 | 0.67916 | NS |
| T_Regulatory | 1 | 0 | 0 | 1 | 6 | 0.355 | 0.645 | 0.32937 | NS |
| Mast | 1 | 2 | 0 | 0 | 5 | 1.066 | 1.934 | 0.95483 | NS |
| B_Plasma | 1 | 1 | 1 | 1 | 3 | 0.711 | 1.289 | 0.76993 | NS |
| Lymphatic | 0 | 0 | 0 | 3 | 4 | 0 | 0 | 1 | NS |
| Club | 0 | 4 | 0 | 0 | 2 | 1.421 | 2.579 | 0.18863 | NS |
| ncMonocyte | 3 | 2 | 0 | 0 | 1 | 1.777 | 3.223 | 0.4386 | NS |
| ILC_B | 0 | 0 | 0 | 2 | 2 | 0 | 0 | 1 | NS |
| ILC_A | 0 | 0 | 0 | 0 | 2 | 0 | 0 | 1 | NS |
| DC_Langerhans | 0 | 2 | 0 | 0 | 0 | 0.711 | 1.289 | 0.35259 | NS |
| pDC | 0 | 0 | 0 | 2 | 0 | 0 | 0 | 1 | NS |
| Basal | 0 | 1 | 0 | 0 | 1 | 0.355 | 0.645 | 0.51098 | NS |
| Ionocyte | 0 | 0 | 0 | 1 | 0 | 0 | 0 | 1 | NS |

Sample:1372C Disease:Control Sex:F Age:21 UMI1:4595 UMI2:3493 total1:572 total2:594

| Region_Celltype | X1 | X2 | Both | Unknown | LowCoverage | Expected1 | Expected2 | P_value | Direction |
|---|---|---|---|---|---|---|---|---|---|
| Macrophage | 221 | 203 | 15 | 467 | 451 | 208 | 216 | 0.37191 | NS |
| Macrophage_Alveolar | 138 | 145 | 40 | 169 | 223 | 138.83 | 144.17 | 0.94435 | NS |
| cMonocyte | 61 | 72 | 4 | 156 | 172 | 65.245 | 67.755 | 0.60218 | NS |
| Fibroblast | 36 | 48 | 5 | 25 | 46 | 41.208 | 42.792 | 0.42014 | NS |
| Lymphatic | 21 | 24 | 0 | 40 | 47 | 22.075 | 22.925 | 0.82047 | NS |
| ATII | 22 | 8 | 2 | 30 | 32 | 14.717 | 15.283 | 0.053675 | NS |
| cDC2 | 16 | 18 | 0 | 24 | 36 | 16.679 | 17.321 | 0.86905 | NS |
| T_Cytotoxic | 4 | 6 | 1 | 41 | 22 | 4.906 | 5.094 | 0.68366 | NS |
| Ciliated | 3 | 3 | 0 | 37 | 18 | 2.943 | 3.057 | 0.97393 | NS |
| VE_Venous | 16 | 17 | 2 | 4 | 17 | 16.189 | 16.811 | 0.96294 | NS |
| Multiplet | 6 | 9 | 7 | 8 | 16 | 7.358 | 7.642 | 0.61775 | NS |
| VE_Arterial | 11 | 13 | 3 | 4 | 12 | 11.774 | 12.226 | 0.82306 | NS |
| T | 2 | 0 | 0 | 20 | 9 | 0.981 | 1.019 | 0.24231 | NS |
| VE_Peribronchial | 4 | 6 | 3 | 5 | 4 | 4.906 | 5.094 | 0.68366 | NS |
| Club | 3 | 3 | 0 | 7 | 8 | 2.943 | 3.057 | 0.97393 | NS |
| Basal | 1 | 1 | 2 | 10 | 6 | 0.981 | 1.019 | 0.98495 | NS |
| ncMonocyte | 3 | 7 | 0 | 7 | 3 | 4.906 | 5.094 | 0.38345 | NS |
| ATI | 1 | 1 | 0 | 8 | 8 | 0.981 | 1.019 | 0.98495 | NS |
| NK | 0 | 1 | 0 | 11 | 5 | 0.491 | 0.509 | 0.42011 | NS |
| DC_Mature | 0 | 0 | 0 | 6 | 3 | 0 | 0 | 1 | NS |
| Goblet | 1 | 2 | 0 | 2 | 4 | 1.472 | 1.528 | 0.69561 | NS |
| VE_Capillary_B | 1 | 3 | 0 | 2 | 2 | 1.962 | 2.038 | 0.48109 | NS |
| Myofibroblast | 0 | 2 | 0 | 1 | 2 | 0.981 | 1.019 | 0.25421 | NS |
| ILC_B | 1 | 0 | 0 | 1 | 3 | 0.491 | 0.509 | 0.40837 | NS |
| B | 0 | 1 | 0 | 0 | 2 | 0.491 | 0.509 | 0.42011 | NS |
| cDC1 | 0 | 0 | 0 | 1 | 2 | 0 | 0 | 1 | NS |
| ILC_A | 0 | 0 | 0 | 2 | 0 | 0 | 0 | 1 | NS |
| B_Plasma | 0 | 0 | 0 | 0 | 1 | 0 | 0 | 1 | NS |
| VE_Capillary_A | 0 | 1 | 0 | 0 | 0 | 0.491 | 0.509 | 0.42011 | NS |
| SMC | 0 | 0 | 0 | 1 | 0 | 0 | 0 | 1 | NS |

Sample:157I Disease:IPF Sex:F Age:66 UMI1:7063 UMI2:5050 total1:990 total2:994

| Region_Celltype | X1 | X2 | Both | Unknown | LowCoverage | Expected1 | Expected2 | P_value | Direction |
|---|---|---|---|---|---|---|---|---|---|
| Macrophage | 544 | 540 | 55 | 440 | 876 | 540.907 | 543.093 | 0.89432 | NS |
| Macrophage_Alveolar | 148 | 134 | 29 | 99 | 220 | 140.716 | 141.284 | 0.53947 | NS |
| cDC2 | 59 | 77 | 14 | 48 | 68 | 67.863 | 68.137 | 0.28138 | NS |
| B | 24 | 19 | 6 | 72 | 111 | 21.457 | 21.543 | 0.58272 | NS |
| Ciliated | 36 | 24 | 5 | 65 | 99 | 29.94 | 30.06 | 0.26616 | NS |
| T_Cytotoxic | 11 | 16 | 6 | 47 | 81 | 13.473 | 13.527 | 0.49906 | NS |
| T | 9 | 18 | 4 | 47 | 74 | 13.473 | 13.527 | 0.2169 | NS |
| Club | 26 | 25 | 1 | 29 | 47 | 25.449 | 25.551 | 0.91304 | NS |
| NK | 8 | 11 | 4 | 36 | 56 | 9.481 | 9.519 | 0.62981 | NS |

| | | | | | | | | | |
|---|---|---|---|---|---|---|---|---|---|
| Multiplet | 14 | 14 | 11 | 16 | 35 | 13.972 | 14.028 | 0.99398 | NS |
| cMonocyte | 19 | 19 | 1 | 14 | 26 | 18.962 | 19.038 | 0.99299 | NS |
| ncMonocyte | 24 | 20 | 3 | 2 | 27 | 21.956 | 22.044 | 0.66263 | NS |
| Goblet | 4 | 6 | 2 | 20 | 23 | 4.99 | 5.01 | 0.65633 | NS |
| DC_Mature | 6 | 2 | 0 | 17 | 17 | 3.992 | 4.008 | 0.29988 | NS |
| Basal | 7 | 6 | 1 | 9 | 14 | 6.487 | 6.513 | 0.84039 | NS |
| T_Regulatory | 2 | 5 | 0 | 13 | 14 | 3.493 | 3.507 | 0.41383 | NS |
| VE_Peribronchial | 6 | 7 | 0 | 6 | 12 | 6.487 | 6.513 | 0.84843 | NS |
| pDC | 3 | 3 | 1 | 12 | 8 | 2.994 | 3.006 | 0.99721 | NS |
| ATII | 4 | 2 | 0 | 9 | 10 | 2.994 | 3.006 | 0.55587 | NS |
| Mast | 4 | 5 | 1 | 4 | 11 | 4.491 | 4.509 | 0.8167 | NS |
| Aberrant_Basaloid | 3 | 5 | 1 | 3 | 11 | 3.992 | 4.008 | 0.61711 | NS |
| Fibroblast | 10 | 9 | 1 | 0 | 3 | 9.481 | 9.519 | 0.8662 | NS |
| cDC1 | 5 | 4 | 3 | 6 | 1 | 4.491 | 4.509 | 0.81007 | NS |
| DC_Langerhans | 4 | 8 | 1 | 1 | 3 | 5.988 | 6.012 | 0.41039 | NS |
| Lymphatic | 3 | 3 | 4 | 3 | 3 | 2.994 | 3.006 | 0.99721 | NS |
| B_Plasma | 3 | 2 | 2 | 1 | 5 | 2.495 | 2.505 | 0.74822 | NS |
| VE_Venous | 2 | 2 | 2 | 2 | 5 | 1.996 | 2.004 | 0.99773 | NS |
| ILC_A | 0 | 1 | 0 | 2 | 7 | 0.499 | 0.501 | 0.41484 | NS |
| Myofibroblast | 0 | 4 | 1 | 2 | 2 | 1.996 | 2.004 | 0.10293 | NS |
| VE_Capillary_B | 2 | 1 | 0 | 2 | 0 | 1.497 | 1.503 | 0.67706 | NS |
| ILC_B | 0 | 0 | 0 | 1 | 3 | 0 | 0 | 1 | NS |
| Pericyte | 0 | 0 | 0 | 1 | 2 | 0 | 0 | 1 | NS |
| ATI | 0 | 0 | 0 | 1 | 1 | 0 | 0 | 1 | NS |
| Mesothelial | 0 | 1 | 0 | 0 | 1 | 0.499 | 0.501 | 0.41484 | NS |
| SMC | 0 | 1 | 0 | 0 | 0 | 0.499 | 0.501 | 0.41484 | NS |
| VE_Arterial | 0 | 0 | 0 | 0 | 1 | 0 | 0 | 1 | NS |

| Sample:174I | Disease:IPF | | | Sex:F | Age:67 | UMI1:5876 | UMI2:3077 | total1:228 | total2:526 |
|---|---|---|---|---|---|---|---|---|---|
| Region_Celltype | X1 | X2 | Both | Unknown | LowCoverage | Expected1 | Expected2 | P_value | Direction |
| Macrophage_Alveolar | 89 | 240 | 28 | 134 | 434 | 99.485 | 229.515 | 0.36592 | NS |
| Macrophage | 75 | 147 | 12 | 251 | 421 | 67.13 | 154.87 | 0.42336 | NS |
| cDC2 | 19 | 49 | 5 | 121 | 92 | 20.562 | 47.438 | 0.76802 | NS |
| B | 6 | 10 | 0 | 49 | 94 | 4.838 | 11.162 | 0.66431 | NS |
| cMonocyte | 2 | 8 | 1 | 54 | 40 | 3.024 | 6.976 | 0.59758 | NS |
| Ciliated | 6 | 10 | 0 | 28 | 48 | 4.838 | 11.162 | 0.66431 | NS |
| Multiplet | 3 | 17 | 3 | 25 | 43 | 6.048 | 13.952 | 0.24939 | NS |
| T | 4 | 2 | 0 | 32 | 44 | 1.814 | 4.186 | 0.20677 | NS |
| pDC | 2 | 1 | 1 | 21 | 14 | 0.907 | 2.093 | 0.372 | NS |
| NK | 2 | 3 | 0 | 7 | 27 | 1.512 | 3.488 | 0.74645 | NS |
| T_Cytotoxic | 0 | 2 | 0 | 12 | 24 | 0.605 | 1.395 | 0.39862 | NS |
| ncMonocyte | 3 | 2 | 0 | 9 | 19 | 1.512 | 3.488 | 0.34433 | NS |
| DC_Mature | 3 | 2 | 1 | 17 | 8 | 1.512 | 3.488 | 0.34433 | NS |

| | | | | | | | | | |
|---|---|---|---|---|---|---|---|---|---|
| Club | 1 | 2 | 0 | 12 | 14 | 0.907 | 2.093 | 0.93513 | NS |
| Basal | 0 | 2 | 0 | 13 | 14 | 0.605 | 1.395 | 0.39862 | NS |
| Aberrant_Basaloid | 2 | 10 | 2 | 1 | 10 | 3.629 | 8.371 | 0.43268 | NS |
| Mast | 2 | 2 | 0 | 3 | 15 | 1.21 | 2.79 | 0.56856 | NS |
| Fibroblast | 2 | 2 | 0 | 2 | 15 | 1.21 | 2.79 | 0.56856 | NS |
| cDC1 | 1 | 3 | 0 | 11 | 3 | 1.21 | 2.79 | 0.86839 | NS |
| Goblet | 0 | 4 | 0 | 4 | 9 | 1.21 | 2.79 | 0.23258 | NS |
| T_Regulatory | 1 | 1 | 0 | 2 | 13 | 0.605 | 1.395 | 0.68682 | NS |
| DC_Langerhans | 0 | 2 | 0 | 5 | 7 | 0.605 | 1.395 | 0.39862 | NS |
| B_Plasma | 3 | 4 | 0 | 0 | 7 | 2.117 | 4.883 | 0.62398 | NS |
| Lymphatic | 0 | 0 | 2 | 1 | 9 | 0 | 0 | 1 | NS |
| VE_Peribronchial | 0 | 0 | 0 | 0 | 8 | 0 | 0 | 1 | NS |
| Myofibroblast | 1 | 1 | 0 | 2 | 3 | 0.605 | 1.395 | 0.68682 | NS |
| VE_Venous | 1 | 0 | 0 | 1 | 3 | 0.302 | 0.698 | 0.30066 | NS |
| VE_Arterial | 0 | 0 | 0 | 2 | 3 | 0 | 0 | 1 | NS |
| ILC_A | 0 | 0 | 0 | 1 | 2 | 0 | 0 | 1 | NS |
| Pericyte | 0 | 0 | 0 | 1 | 1 | 0 | 0 | 1 | NS |
| ATII | 0 | 0 | 0 | 0 | 1 | 0 | 0 | 1 | NS |
| VE_Capillary_B | 0 | 0 | 0 | 0 | 1 | 0 | 0 | 1 | NS |
| ILC_B | 0 | 0 | 0 | 0 | 1 | 0 | 0 | 1 | NS |

| Sample:178CO | Disease:COPD | | | Sex:F | Age:58 | UMI1:8879 | UMI2:2934 | total1:238 | total2:259 |
|---|---|---|---|---|---|---|---|---|---|
| Region_Celltype | X1 | X2 | Both | Unknown | LowCoverage | Expected1 | Expected2 | P_value | Direction |
| Macrophage_Alveolar | 176 | 180 | 14 | 1171 | 353 | 170.479 | 185.521 | 0.67889 | NS |
| Macrophage | 15 | 15 | 3 | 131 | 28 | 14.366 | 15.634 | 0.86998 | NS |
| T_Cytotoxic | 6 | 7 | 0 | 117 | 15 | 6.225 | 6.775 | 0.92944 | NS |
| Multiplet | 8 | 3 | 2 | 95 | 24 | 5.268 | 5.732 | 0.23378 | NS |
| cDC1 | 5 | 10 | 1 | 32 | 21 | 7.183 | 7.817 | 0.41702 | NS |
| cMonocyte | 5 | 8 | 0 | 46 | 7 | 6.225 | 6.775 | 0.62756 | NS |
| cDC2 | 6 | 10 | 0 | 35 | 15 | 7.662 | 8.338 | 0.55253 | NS |
| ncMonocyte | 1 | 2 | 0 | 49 | 11 | 1.437 | 1.563 | 0.71664 | NS |
| T | 4 | 5 | 0 | 48 | 6 | 4.31 | 4.69 | 0.88353 | NS |
| NK | 5 | 4 | 0 | 35 | 8 | 4.31 | 4.69 | 0.74478 | NS |
| B | 0 | 2 | 0 | 25 | 8 | 0.958 | 1.042 | 0.26179 | NS |
| B_Plasma | 0 | 4 | 1 | 13 | 7 | 1.915 | 2.085 | 0.11252 | NS |
| pDC | 1 | 1 | 0 | 17 | 5 | 0.958 | 1.042 | 0.96629 | NS |
| DC_Mature | 0 | 4 | 0 | 11 | 5 | 1.915 | 2.085 | 0.11252 | NS |
| Mast | 0 | 0 | 0 | 8 | 4 | 0 | 0 | 1 | NS |
| Club | 0 | 1 | 0 | 6 | 5 | 0.479 | 0.521 | 0.42749 | NS |
| Mesothelial | 1 | 1 | 0 | 7 | 3 | 0.958 | 1.042 | 0.96629 | NS |
| Goblet | 1 | 0 | 0 | 7 | 2 | 0.479 | 0.521 | 0.40119 | NS |
| ATII | 2 | 0 | 0 | 7 | 0 | 0.958 | 1.042 | 0.23514 | NS |
| T_Regulatory | 0 | 1 | 0 | 3 | 1 | 0.479 | 0.521 | 0.42749 | NS |

| | | | | | | | | | |
|---|---|---|---|---|---|---|---|---|---|
| DC_Langerhans | 1 | 1 | 0 | 1 | 2 | 0.958 | 1.042 | 0.96629 | NS |
| Ciliated | 0 | 0 | 0 | 5 | 0 | 0 | 0 | 1 | NS |
| Fibroblast | 0 | 0 | 0 | 4 | 0 | 0 | 0 | 1 | NS |
| Lymphatic | 0 | 0 | 0 | 3 | 0 | 0 | 0 | 1 | NS |
| Basal | 0 | 0 | 0 | 2 | 1 | 0 | 0 | 1 | NS |
| ILC_A | 0 | 0 | 0 | 2 | 1 | 0 | 0 | 1 | NS |
| ATI | 1 | 0 | 0 | 2 | 0 | 0.479 | 0.521 | 0.40119 | NS |
| VE_Arterial | 0 | 0 | 0 | 1 | 0 | 0 | 0 | 1 | NS |
| PNEC | 0 | 0 | 0 | 1 | 0 | 0 | 0 | 1 | NS |

| Sample:184CO | Disease:COPD | | | Sex:F | Age:55 | UMI1:6514 | UMI2:4576 | total1:1011 | total2:1831 |
|---|---|---|---|---|---|---|---|---|---|
| Region_Celltype | X1 | X2 | Both | Unknown | LowCoverage | Expected1 | Expected2 | P_value | Direction |
| Macrophage_Alveolar | 219 | 365 | 68 | 40 | 187 | 207.749 | 376.251 | 0.4942 | NS |
| Macrophage | 171 | 360 | 26 | 24 | 114 | 188.895 | 342.105 | 0.24598 | NS |
| cMonocyte | 180 | 377 | 15 | 12 | 111 | 198.145 | 358.855 | 0.25094 | NS |
| ncMonocyte | 171 | 330 | 8 | 4 | 49 | 178.223 | 322.777 | 0.63201 | NS |
| cDC2 | 91 | 192 | 7 | 17 | 44 | 100.673 | 182.327 | 0.39026 | NS |
| T | 17 | 33 | 5 | 114 | 160 | 17.787 | 32.213 | 0.8688 | NS |
| NK | 4 | 22 | 3 | 96 | 129 | 9.249 | 16.751 | 0.094815 | NS |
| T_Cytotoxic | 17 | 8 | 0 | 79 | 99 | 8.893 | 16.107 | 0.02177 | NS |
| Multiplet | 26 | 37 | 21 | 7 | 26 | 22.411 | 40.589 | 0.51101 | NS |
| Ciliated | 25 | 7 | 3 | 10 | 18 | 11.384 | 20.616 | 0.00058925 | Uniform |
| cDC1 | 15 | 27 | 2 | 2 | 6 | 14.941 | 27.059 | 0.98926 | NS |
| B | 2 | 6 | 4 | 16 | 18 | 2.846 | 5.154 | 0.64536 | NS |
| DC_Mature | 11 | 19 | 0 | 4 | 8 | 10.672 | 19.328 | 0.92977 | NS |
| T_Regulatory | 3 | 2 | 1 | 8 | 23 | 1.779 | 3.221 | 0.43941 | NS |
| ILC_A | 2 | 2 | 0 | 11 | 14 | 1.423 | 2.577 | 0.68008 | NS |
| Mast | 3 | 6 | 2 | 3 | 10 | 3.202 | 5.798 | 0.92034 | NS |
| ATII | 7 | 5 | 0 | 1 | 9 | 4.269 | 7.731 | 0.26396 | NS |
| Club | 7 | 2 | 3 | 1 | 7 | 3.202 | 5.798 | 0.070801 | NS |
| Fibroblast | 9 | 6 | 0 | 0 | 4 | 5.336 | 9.664 | 0.1805 | NS |
| VE_Arterial | 3 | 1 | 1 | 1 | 10 | 1.423 | 2.577 | 0.2621 | NS |
| Goblet | 7 | 4 | 0 | 0 | 3 | 3.913 | 7.087 | 0.18807 | NS |
| ILC_B | 0 | 2 | 0 | 7 | 5 | 0.711 | 1.289 | 0.35223 | NS |
| Lymphatic | 2 | 0 | 0 | 2 | 9 | 0.711 | 1.289 | 0.16798 | NS |
| PNEC | 2 | 0 | 0 | 3 | 7 | 0.711 | 1.289 | 0.16798 | NS |
| B_Plasma | 1 | 3 | 0 | 1 | 5 | 1.423 | 2.577 | 0.74486 | NS |
| pDC | 1 | 2 | 0 | 4 | 2 | 1.067 | 1.933 | 0.95396 | NS |
| Myofibroblast | 4 | 2 | 0 | 0 | 3 | 2.134 | 3.866 | 0.28132 | NS |
| Mesothelial | 3 | 3 | 0 | 0 | 2 | 2.134 | 3.866 | 0.61354 | NS |
| Basal | 2 | 1 | 0 | 0 | 5 | 1.067 | 1.933 | 0.44617 | NS |
| DC_Langerhans | 1 | 3 | 0 | 1 | 1 | 1.423 | 2.577 | 0.74486 | NS |
| VE_Venous | 2 | 1 | 0 | 0 | 2 | 1.067 | 1.933 | 0.44617 | NS |

| | | | | | | | | | |
|---|---|---|---|---|---|---|---|---|---|
| ATI | 2 | 1 | 0 | 0 | 1 | 1.067 | 1.933 | 0.44617 | NS |
| VE_Capillary_B | 1 | 1 | 0 | 1 | 1 | 0.711 | 1.289 | 0.77061 | NS |
| VE_Peribronchial | 0 | 1 | 1 | 2 | 0 | 0.356 | 0.644 | 0.51067 | NS |
| Pericyte | 0 | 0 | 0 | 0 | 1 | 0 | 0 | 1 | NS |

Sample:207CO    Disease:COPD    Sex:F    Age:60    UMI1:3165    UMI2:1218    total1:62    total2:146

| Region_Celltype | X1 | X2 | Both | Unknown | LowCoverage | Expected1 | Expected2 | P_value | Direction |
|---|---|---|---|---|---|---|---|---|---|
| Macrophage | 14 | 19 | 1 | 141 | 23 | 9.837 | 23.163 | 0.286 | NS |
| B | 13 | 17 | 2 | 108 | 26 | 8.942 | 21.058 | 0.27675 | NS |
| cMonocyte | 4 | 13 | 1 | 97 | 26 | 5.067 | 11.933 | 0.67894 | NS |
| Macrophage_Alveolar | 7 | 2 | 0 | 82 | 28 | 2.683 | 6.317 | 0.041243 | NS |
| NK | 2 | 24 | 1 | 69 | 10 | 7.75 | 18.25 | 0.041059 | NS |
| T | 6 | 16 | 1 | 53 | 21 | 6.558 | 15.442 | 0.85231 | NS |
| T_Cytotoxic | 2 | 17 | 0 | 47 | 14 | 5.663 | 13.337 | 0.13858 | NS |
| Multiplet | 1 | 17 | 2 | 36 | 19 | 5.365 | 12.635 | 0.056514 | NS |
| ncMonocyte | 1 | 2 | 0 | 26 | 11 | 0.894 | 2.106 | 0.92598 | NS |
| cDC2 | 2 | 9 | 2 | 21 | 5 | 3.279 | 7.721 | 0.52318 | NS |
| B_Plasma | 0 | 3 | 0 | 17 | 5 | 0.894 | 2.106 | 0.30531 | NS |
| ATII | 5 | 1 | 0 | 12 | 4 | 1.788 | 4.212 | 0.061428 | NS |
| pDC | 1 | 3 | 0 | 8 | 1 | 1.192 | 2.808 | 0.87884 | NS |
| Club | 0 | 0 | 0 | 9 | 4 | 0 | 0 | 1 | NS |
| Mesothelial | 0 | 0 | 0 | 11 | 2 | 0 | 0 | 1 | NS |
| Fibroblast | 0 | 0 | 0 | 11 | 1 | 0 | 0 | 1 | NS |
| cDC1 | 0 | 0 | 1 | 7 | 4 | 0 | 0 | 1 | NS |
| Ciliated | 1 | 0 | 0 | 7 | 1 | 0.298 | 0.702 | 0.29837 | NS |
| T_Regulatory | 0 | 0 | 0 | 5 | 3 | 0 | 0 | 1 | NS |
| VE_Capillary_B | 0 | 0 | 0 | 4 | 1 | 0 | 0 | 1 | NS |
| Myofibroblast | 0 | 1 | 0 | 4 | 0 | 0.298 | 0.702 | 0.55395 | NS |
| DC_Mature | 1 | 0 | 0 | 0 | 2 | 0.298 | 0.702 | 0.29837 | NS |
| ILC_A | 1 | 1 | 0 | 0 | 1 | 0.596 | 1.404 | 0.68009 | NS |
| VE_Peribronchial | 0 | 0 | 0 | 2 | 0 | 0 | 0 | 1 | NS |
| ILC_B | 0 | 0 | 0 | 1 | 1 | 0 | 0 | 1 | NS |
| Lymphatic | 0 | 0 | 0 | 1 | 1 | 0 | 0 | 1 | NS |
| VE_Venous | 1 | 0 | 0 | 1 | 0 | 0.298 | 0.702 | 0.29837 | NS |
| Pericyte | 0 | 0 | 0 | 0 | 1 | 0 | 0 | 1 | NS |
| Mast | 0 | 0 | 0 | 1 | 0 | 0 | 0 | 1 | NS |
| Basal | 0 | 0 | 0 | 1 | 0 | 0 | 0 | 1 | NS |
| SMC | 0 | 0 | 0 | 0 | 1 | 0 | 0 | 1 | NS |
| PNEC | 0 | 0 | 0 | 1 | 0 | 0 | 0 | 1 | NS |
| Aberrant_Basaloid | 0 | 1 | 0 | 0 | 0 | 0.298 | 0.702 | 0.55395 | NS |

Sample:209I    Disease:IPF    Sex:F    Age:65    UMI1:4592    UMI2:2894    total1:440    total2:742

| Region_Celltype | X1 | X2 | Both | Unknown | LowCoverage | Expected1 | Expected2 | P_value | Direction |
|---|---|---|---|---|---|---|---|---|---|

| | | | | | | | | | |
|---|---|---|---|---|---|---|---|---|---|
| Macrophage | 254 | 436 | 37 | 275 | 538 | 256.853 | 433.147 | 0.87364 | NS |
| Macrophage_Alveolar | 93 | 150 | 19 | 57 | 145 | 90.457 | 152.543 | 0.8119 | NS |
| Ciliated | 16 | 29 | 3 | 125 | 93 | 16.751 | 28.249 | 0.86926 | NS |
| cMonocyte | 8 | 15 | 1 | 27 | 54 | 8.562 | 14.438 | 0.863 | NS |
| cDC2 | 16 | 11 | 2 | 13 | 22 | 10.051 | 16.949 | 0.10519 | NS |
| B | 4 | 17 | 0 | 14 | 28 | 7.817 | 13.183 | 0.19023 | NS |
| Aberrant_Basaloid | 13 | 22 | 4 | 3 | 20 | 13.029 | 21.971 | 0.99432 | NS |
| Multiplet | 8 | 17 | 4 | 9 | 21 | 9.306 | 15.694 | 0.69778 | NS |
| T_Cytotoxic | 7 | 5 | 2 | 12 | 24 | 4.467 | 7.533 | 0.30062 | NS |
| Goblet | 5 | 10 | 0 | 7 | 12 | 5.584 | 9.416 | 0.8235 | NS |
| NK | 3 | 3 | 0 | 7 | 16 | 2.234 | 3.766 | 0.65546 | NS |
| T | 1 | 1 | 0 | 6 | 14 | 0.745 | 1.255 | 0.79671 | NS |
| VE_Peribronchial | 1 | 4 | 0 | 6 | 10 | 1.861 | 3.139 | 0.54676 | NS |
| Basal | 2 | 3 | 0 | 6 | 8 | 1.861 | 3.139 | 0.92819 | NS |
| Club | 1 | 4 | 0 | 4 | 10 | 1.861 | 3.139 | 0.54676 | NS |
| Mesothelial | 1 | 2 | 0 | 7 | 6 | 1.117 | 1.883 | 0.92054 | NS |
| ncMonocyte | 0 | 2 | 0 | 5 | 5 | 0.745 | 1.255 | 0.33885 | NS |
| Fibroblast | 1 | 3 | 0 | 0 | 3 | 1.489 | 2.511 | 0.70882 | NS |
| Myofibroblast | 2 | 1 | 1 | 0 | 2 | 1.117 | 1.883 | 0.47047 | NS |
| Mast | 1 | 0 | 0 | 0 | 5 | 0.372 | 0.628 | 0.33881 | NS |
| Pericyte | 0 | 2 | 0 | 2 | 2 | 0.745 | 1.255 | 0.33885 | NS |
| DC_Langerhans | 3 | 1 | 0 | 0 | 0 | 1.489 | 2.511 | 0.2817 | NS |
| DC_Mature | 0 | 0 | 0 | 3 | 1 | 0 | 0 | 1 | NS |
| T_Regulatory | 0 | 0 | 0 | 1 | 3 | 0 | 0 | 1 | NS |
| VE_Venous | 0 | 1 | 0 | 1 | 1 | 0.372 | 0.628 | 0.49885 | NS |
| SMC | 0 | 0 | 0 | 2 | 1 | 0 | 0 | 1 | NS |
| VE_Arterial | 0 | 1 | 1 | 0 | 0 | 0.372 | 0.628 | 0.49885 | NS |
| Lymphatic | 0 | 1 | 0 | 1 | 0 | 0.372 | 0.628 | 0.49885 | NS |
| cDC1 | 0 | 1 | 0 | 0 | 0 | 0.372 | 0.628 | 0.49885 | NS |
| ILC_A | 0 | 0 | 0 | 0 | 1 | 0 | 0 | 1 | NS |

Sample:217CO Disease:COPD Sex:F Age:70 UMI1:2573 UMI2:1474 total1:116 total2:136

| Region_Celltype | X1 | X2 | Both | Unknown | LowCoverage | Expected1 | Expected2 | P_value | Direction |
|---|---|---|---|---|---|---|---|---|---|
| cMonocyte | 34 | 55 | 5 | 371 | 81 | 40.968 | 48.032 | 0.29014 | NS |
| Macrophage_Alveolar | 26 | 15 | 1 | 59 | 45 | 18.873 | 22.127 | 0.11384 | NS |
| NK | 16 | 6 | 5 | 73 | 39 | 10.127 | 11.873 | 0.071422 | NS |
| Macrophage | 14 | 13 | 1 | 73 | 30 | 12.429 | 14.571 | 0.66881 | NS |
| T_Cytotoxic | 2 | 5 | 0 | 50 | 49 | 3.222 | 3.778 | 0.49939 | NS |
| T | 7 | 1 | 1 | 45 | 35 | 3.683 | 4.317 | 0.078295 | NS |
| ncMonocyte | 0 | 6 | 0 | 42 | 14 | 2.762 | 3.238 | 0.058211 | NS |
| Multiplet | 4 | 7 | 0 | 20 | 12 | 5.063 | 5.937 | 0.64504 | NS |
| cDC2 | 2 | 7 | 8 | 13 | 7 | 4.143 | 4.857 | 0.28676 | NS |
| Ciliated | 4 | 2 | 0 | 11 | 16 | 2.762 | 3.238 | 0.47113 | NS |

| | | | | | | | | | |
|---|---|---|---|---|---|---|---|---|---|
| B | 0 | 4 | 0 | 15 | 13 | 1.841 | 2.159 | 0.12198 | NS |
| Club | 0 | 2 | 0 | 10 | 7 | 0.921 | 1.079 | 0.27415 | NS |
| ATII | 1 | 2 | 0 | 6 | 3 | 1.381 | 1.619 | 0.75057 | NS |
| Lymphatic | 0 | 1 | 0 | 4 | 5 | 0.46 | 0.54 | 0.43937 | NS |
| Mesothelial | 0 | 2 | 0 | 3 | 2 | 0.921 | 1.079 | 0.27415 | NS |
| Basal | 0 | 0 | 0 | 4 | 3 | 0 | 0 | 1 | NS |
| VE_Capillary_B | 1 | 0 | 0 | 3 | 3 | 0.46 | 0.54 | 0.38994 | NS |
| Goblet | 2 | 0 | 0 | 3 | 1 | 0.921 | 1.079 | 0.22405 | NS |
| ATI | 0 | 1 | 0 | 1 | 4 | 0.46 | 0.54 | 0.43937 | NS |
| SMC | 0 | 1 | 0 | 4 | 0 | 0.46 | 0.54 | 0.43937 | NS |
| Myofibroblast | 0 | 0 | 0 | 3 | 2 | 0 | 0 | 1 | NS |
| Mast | 2 | 2 | 1 | 0 | 0 | 1.841 | 2.159 | 0.91056 | NS |
| Fibroblast | 0 | 0 | 0 | 2 | 2 | 0 | 0 | 1 | NS |
| VE_Venous | 0 | 1 | 0 | 2 | 0 | 0.46 | 0.54 | 0.43937 | NS |
| VE_Arterial | 1 | 1 | 0 | 1 | 0 | 0.921 | 1.079 | 0.93669 | NS |
| cDC1 | 0 | 0 | 0 | 1 | 1 | 0 | 0 | 1 | NS |
| VE_Capillary_A | 0 | 1 | 0 | 1 | 0 | 0.46 | 0.54 | 0.43937 | NS |
| Pericyte | 0 | 1 | 0 | 0 | 1 | 0.46 | 0.54 | 0.43937 | NS |
| PNEC | 0 | 0 | 0 | 0 | 1 | 0 | 0 | 1 | NS |
| ILC_A | 0 | 0 | 0 | 1 | 0 | 0 | 0 | 1 | NS |
| ILC_B | 0 | 0 | 0 | 0 | 1 | 0 | 0 | 1 | NS |
| T_Regulatory | 0 | 0 | 0 | 1 | 0 | 0 | 0 | 1 | NS |
| pDC | 0 | 0 | 0 | 0 | 1 | 0 | 0 | 1 | NS |

| Sample:221I | Disease:IPF | | | Sex:F | Age:67 | UMI1:5330 | UMI2:2989 | total1:45 | total2:86 |
|---|---|---|---|---|---|---|---|---|---|
| Region_Celltype | X1 | X2 | Both | Unknown | LowCoverage | Expected1 | Expected2 | P_value | Direction |
| Macrophage | 24 | 48 | 2 | 1404 | 376 | 24.733 | 47.267 | 0.89731 | NS |
| Macrophage_Alveolar | 9 | 13 | 2 | 232 | 71 | 7.557 | 14.443 | 0.65346 | NS |
| Ciliated | 4 | 2 | 0 | 62 | 48 | 2.061 | 3.939 | 0.26293 | NS |
| cMonocyte | 0 | 6 | 0 | 84 | 19 | 2.061 | 3.939 | 0.11468 | NS |
| Multiplet | 1 | 5 | 0 | 77 | 16 | 2.061 | 3.939 | 0.48226 | NS |
| NK | 0 | 2 | 0 | 71 | 11 | 0.687 | 1.313 | 0.36242 | NS |
| cDC2 | 5 | 7 | 2 | 49 | 21 | 4.122 | 7.878 | 0.71201 | NS |
| B | 0 | 0 | 0 | 64 | 11 | 0 | 0 | 1 | NS |
| ncMonocyte | 0 | 1 | 0 | 57 | 10 | 0.344 | 0.656 | 0.51957 | NS |
| T | 0 | 0 | 0 | 25 | 4 | 0 | 0 | 1 | NS |
| Club | 0 | 0 | 0 | 11 | 10 | 0 | 0 | 1 | NS |
| T_Cytotoxic | 0 | 0 | 0 | 16 | 4 | 0 | 0 | 1 | NS |
| Goblet | 0 | 0 | 0 | 13 | 7 | 0 | 0 | 1 | NS |
| Basal | 0 | 0 | 0 | 7 | 7 | 0 | 0 | 1 | NS |
| DC_Langerhans | 0 | 1 | 0 | 7 | 5 | 0.344 | 0.656 | 0.51957 | NS |
| Fibroblast | 0 | 0 | 0 | 9 | 1 | 0 | 0 | 1 | NS |
| pDC | 1 | 0 | 0 | 3 | 2 | 0.344 | 0.656 | 0.32287 | NS |

| | | | | | | | | | |
|---|---|---|---|---|---|---|---|---|---|
| Mast | 1 | 0 | 0 | 4 | 0 | 0.344 | 0.656 | 0.32287 | NS |
| T_Regulatory | 0 | 0 | 0 | 5 | 0 | 0 | 0 | 1 | NS |
| Lymphatic | 0 | 0 | 0 | 4 | 1 | 0 | 0 | 1 | NS |
| DC_Mature | 0 | 0 | 0 | 0 | 5 | 0 | 0 | 1 | NS |
| VE_Peribronchial | 0 | 0 | 0 | 3 | 1 | 0 | 0 | 1 | NS |
| cDC1 | 0 | 1 | 0 | 2 | 1 | 0.344 | 0.656 | 0.51957 | NS |
| ILC_A | 0 | 0 | 0 | 1 | 1 | 0 | 0 | 1 | NS |
| Aberrant_Basaloid | 0 | 0 | 0 | 2 | 0 | 0 | 0 | 1 | NS |
| Pericyte | 0 | 0 | 0 | 1 | 0 | 0 | 0 | 1 | NS |
| Ionocyte | 0 | 0 | 0 | 1 | 0 | 0 | 0 | 1 | NS |
| VE_Arterial | 0 | 0 | 0 | 1 | 0 | 0 | 0 | 1 | NS |
| Myofibroblast | 0 | 0 | 0 | 1 | 0 | 0 | 0 | 1 | NS |
| ILC_B | 0 | 0 | 0 | 1 | 0 | 0 | 0 | 1 | NS |
| SMC | 0 | 0 | 0 | 1 | 0 | 0 | 0 | 1 | NS |
| VE_Venous | 0 | 0 | 0 | 1 | 0 | 0 | 0 | 1 | NS |
| B_Plasma | 0 | 0 | 0 | 1 | 0 | 0 | 0 | 1 | NS |

| Sample:225I | Disease:IPF | | | Sex:F | Age:70 | UMI1:12085 | UMI2:8032 | total1:549 | total2:890 |
|---|---|---|---|---|---|---|---|---|---|
| Region_Celltype | X1 | X2 | Both | Unknown | LowCoverage | Expected1 | Expected2 | P_value | Direction |
| Macrophage | 185 | 318 | 32 | 1818 | 1020 | 191.902 | 311.098 | 0.65302 | NS |
| Macrophage_Alveolar | 178 | 354 | 42 | 826 | 784 | 202.966 | 329.034 | 0.11039 | NS |
| Myofibroblast | 45 | 36 | 3 | 160 | 139 | 30.903 | 50.097 | 0.026448 | NS |
| Ciliated | 55 | 32 | 5 | 131 | 111 | 33.192 | 53.808 | 0.00094348 | Uniform |
| cDC2 | 14 | 23 | 1 | 188 | 85 | 14.116 | 22.884 | 0.97783 | NS |
| Multiplet | 15 | 23 | 17 | 84 | 79 | 14.498 | 23.502 | 0.90586 | NS |
| cMonocyte | 5 | 7 | 1 | 169 | 32 | 4.578 | 7.422 | 0.86043 | NS |
| VE_Peribronchial | 7 | 30 | 2 | 51 | 40 | 14.116 | 22.884 | 0.066976 | NS |
| NK | 4 | 10 | 1 | 52 | 52 | 5.341 | 8.659 | 0.59088 | NS |
| ncMonocyte | 6 | 4 | 1 | 69 | 32 | 3.815 | 6.185 | 0.32844 | NS |
| T | 5 | 3 | 2 | 32 | 25 | 3.052 | 4.948 | 0.33008 | NS |
| Pericyte | 5 | 4 | 1 | 26 | 25 | 3.434 | 5.566 | 0.45939 | NS |
| VE_Capillary_B | 0 | 9 | 2 | 28 | 20 | 3.434 | 5.566 | 0.039412 | NS |
| SMC | 5 | 4 | 0 | 20 | 30 | 3.434 | 5.566 | 0.45939 | NS |
| T_Cytotoxic | 2 | 1 | 3 | 26 | 23 | 1.145 | 1.855 | 0.48437 | NS |
| VE_Venous | 3 | 6 | 1 | 19 | 11 | 3.434 | 5.566 | 0.83111 | NS |
| Mast | 0 | 2 | 0 | 21 | 12 | 0.763 | 1.237 | 0.33153 | NS |
| B | 2 | 2 | 0 | 21 | 6 | 1.526 | 2.474 | 0.73574 | NS |
| pDC | 3 | 4 | 0 | 19 | 4 | 2.671 | 4.329 | 0.85768 | NS |
| DC_Langerhans | 2 | 2 | 1 | 16 | 9 | 1.526 | 2.474 | 0.73574 | NS |
| VE_Arterial | 0 | 6 | 1 | 11 | 6 | 2.289 | 3.711 | 0.092594 | NS |
| Fibroblast | 0 | 1 | 0 | 12 | 8 | 0.382 | 0.618 | 0.49232 | NS |
| Lymphatic | 2 | 0 | 1 | 9 | 7 | 0.763 | 1.237 | 0.18084 | NS |
| VE_Capillary_A | 0 | 0 | 0 | 11 | 4 | 0 | 0 | 1 | NS |

| | | | | | | | | | |
|---|---|---|---|---|---|---|---|---|---|
| DC_Mature | 3 | 0 | 0 | 5 | 6 | 1.145 | 1.855 | 0.10123 | NS |
| cDC1 | 0 | 0 | 0 | 7 | 7 | 0 | 0 | 1 | NS |
| ATII | 0 | 0 | 0 | 11 | 3 | 0 | 0 | 1 | NS |
| Basal | 0 | 3 | 0 | 3 | 5 | 1.145 | 1.855 | 0.23434 | NS |
| T_Regulatory | 1 | 0 | 0 | 5 | 4 | 0.382 | 0.618 | 0.34403 | NS |
| Goblet | 0 | 2 | 0 | 7 | 1 | 0.763 | 1.237 | 0.33153 | NS |
| ILC_B | 0 | 1 | 0 | 5 | 2 | 0.382 | 0.618 | 0.49232 | NS |
| ILC_A | 0 | 0 | 0 | 4 | 3 | 0 | 0 | 1 | NS |
| Club | 1 | 0 | 0 | 2 | 3 | 0.382 | 0.618 | 0.34403 | NS |
| Aberrant_Basaloid | 1 | 2 | 1 | 0 | 2 | 1.145 | 1.855 | 0.902 | NS |
| Mesothelial | 0 | 1 | 0 | 2 | 2 | 0.382 | 0.618 | 0.49232 | NS |
| ATI | 0 | 0 | 1 | 1 | 1 | 0 | 0 | 1 | NS |

Sample:235CO Disease:COPD Sex:F Age:61 UMI1:4176 UMI2:2094 total1:366 total2:401

| Region_Celltype | X1 | X2 | Both | Unknown | LowCoverage | Expected1 | Expected2 | P_value | Direction |
|---|---|---|---|---|---|---|---|---|---|
| Macrophage_Alveolar | 192 | 192 | 22 | 83 | 554 | 183.239 | 200.761 | 0.52708 | NS |
| Macrophage | 52 | 59 | 5 | 21 | 118 | 52.967 | 58.033 | 0.89653 | NS |
| cMonocyte | 25 | 31 | 1 | 2 | 29 | 26.722 | 29.278 | 0.74409 | NS |
| Multiplet | 7 | 12 | 15 | 4 | 28 | 9.066 | 9.934 | 0.49739 | NS |
| ATII | 14 | 7 | 0 | 9 | 34 | 10.021 | 10.979 | 0.21464 | NS |
| B | 2 | 3 | 0 | 6 | 53 | 2.386 | 2.614 | 0.80573 | NS |
| Mast | 7 | 9 | 0 | 5 | 29 | 7.635 | 8.365 | 0.82174 | NS |
| cDC2 | 11 | 12 | 0 | 5 | 17 | 10.975 | 12.025 | 0.99417 | NS |
| Club | 7 | 7 | 0 | 4 | 25 | 6.681 | 7.319 | 0.90388 | NS |
| Ciliated | 8 | 7 | 4 | 4 | 19 | 7.158 | 7.842 | 0.75842 | NS |
| NK | 2 | 3 | 1 | 2 | 34 | 2.386 | 2.614 | 0.80573 | NS |
| T_Cytotoxic | 2 | 5 | 0 | 6 | 27 | 3.34 | 3.66 | 0.46085 | NS |
| ncMonocyte | 13 | 8 | 0 | 1 | 16 | 10.021 | 10.979 | 0.35566 | NS |
| B_Plasma | 4 | 4 | 2 | 1 | 24 | 3.817 | 4.183 | 0.92726 | NS |
| Myofibroblast | 3 | 9 | 0 | 2 | 20 | 5.726 | 6.274 | 0.24733 | NS |
| T | 0 | 2 | 0 | 3 | 21 | 0.954 | 1.046 | 0.2629 | NS |
| cDC1 | 3 | 6 | 0 | 0 | 8 | 4.295 | 4.705 | 0.53423 | NS |
| Basal | 1 | 4 | 1 | 0 | 7 | 2.386 | 2.614 | 0.35439 | NS |
| Fibroblast | 2 | 0 | 1 | 0 | 9 | 0.954 | 1.046 | 0.23411 | NS |
| DC_Mature | 1 | 6 | 0 | 4 | 0 | 3.34 | 3.66 | 0.17626 | NS |
| VE_Peribronchial | 4 | 1 | 0 | 1 | 5 | 2.386 | 2.614 | 0.28802 | NS |
| Lymphatic | 0 | 2 | 0 | 2 | 6 | 0.954 | 1.046 | 0.2629 | NS |
| Goblet | 2 | 2 | 0 | 1 | 2 | 1.909 | 2.091 | 0.94853 | NS |
| Aberrant_Basaloid | 2 | 0 | 0 | 2 | 1 | 0.954 | 1.046 | 0.23411 | NS |
| Pericyte | 1 | 2 | 0 | 0 | 2 | 1.432 | 1.568 | 0.7197 | NS |
| T_Regulatory | 0 | 1 | 0 | 1 | 3 | 0.477 | 0.523 | 0.42856 | NS |
| ILC_B | 0 | 1 | 1 | 0 | 3 | 0.477 | 0.523 | 0.42856 | NS |
| VE_Arterial | 0 | 1 | 1 | 0 | 2 | 0.477 | 0.523 | 0.42856 | NS |

| | | | | | | | | | |
|---|---|---|---|---|---|---|---|---|---|
| Mesothelial | 1 | 2 | 0 | 0 | 0 | 1.432 | 1.568 | 0.7197 | NS |
| ILC_A | 0 | 1 | 0 | 1 | 1 | 0.477 | 0.523 | 0.42856 | NS |
| VE_Capillary_B | 0 | 1 | 0 | 0 | 2 | 0.477 | 0.523 | 0.42856 | NS |
| SMC | 0 | 0 | 0 | 0 | 2 | 0 | 0 | 1 | NS |
| VE_Venous | 0 | 0 | 0 | 0 | 2 | 0 | 0 | 1 | NS |
| DC_Langerhans | 0 | 1 | 0 | 0 | 0 | 0.477 | 0.523 | 0.42856 | NS |

| Sample:237CO | Disease:COPD | | | Sex:F | Age:57 | UMI1:3566 | UMI2:1888 | total1:106 | total2:351 |
|---|---|---|---|---|---|---|---|---|---|
| Region_Celltype | X1 | X2 | Both | Unknown | LowCoverage | Expected1 | Expected2 | P_value | Direction |
| Macrophage | 20 | 103 | 12 | 188 | 157 | 28.53 | 94.47 | 0.17175 | NS |
| B | 22 | 42 | 0 | 87 | 82 | 14.845 | 49.155 | 0.16245 | NS |
| cMonocyte | 3 | 39 | 5 | 101 | 82 | 9.742 | 32.258 | 0.040305 | NS |
| cDC2 | 3 | 39 | 10 | 52 | 49 | 9.742 | 32.258 | 0.040305 | NS |
| ncMonocyte | 3 | 22 | 2 | 55 | 59 | 5.799 | 19.201 | 0.29863 | NS |
| Macrophage_Alveolar | 7 | 24 | 2 | 32 | 26 | 7.19 | 23.81 | 0.95411 | NS |
| Mast | 6 | 18 | 2 | 29 | 33 | 5.567 | 18.433 | 0.88375 | NS |
| NK | 4 | 6 | 1 | 35 | 16 | 2.319 | 7.681 | 0.41892 | NS |
| T | 2 | 3 | 1 | 33 | 19 | 1.16 | 3.84 | 0.56763 | NS |
| cDC1 | 4 | 12 | 6 | 11 | 11 | 3.711 | 12.289 | 0.90497 | NS |
| B_Plasma | 1 | 3 | 0 | 17 | 13 | 0.928 | 3.072 | 0.9524 | NS |
| ATII | 6 | 5 | 0 | 14 | 9 | 2.551 | 8.449 | 0.13147 | NS |
| Multiplet | 3 | 3 | 1 | 12 | 11 | 1.392 | 4.608 | 0.33513 | NS |
| T_Cytotoxic | 3 | 3 | 0 | 11 | 12 | 1.392 | 4.608 | 0.33513 | NS |
| Ciliated | 3 | 9 | 0 | 6 | 7 | 2.783 | 9.217 | 0.91765 | NS |
| Mesothelial | 1 | 4 | 1 | 5 | 7 | 1.16 | 3.84 | 0.9023 | NS |
| Lymphatic | 0 | 2 | 0 | 7 | 5 | 0.464 | 1.536 | 0.46882 | NS |
| Fibroblast | 0 | 0 | 0 | 9 | 5 | 0 | 0 | 1 | NS |
| VE_Capillary_B | 4 | 1 | 1 | 3 | 3 | 1.16 | 3.84 | 0.072295 | NS |
| Myofibroblast | 2 | 1 | 0 | 3 | 5 | 0.696 | 2.304 | 0.28446 | NS |
| pDC | 0 | 3 | 1 | 2 | 4 | 0.696 | 2.304 | 0.37497 | NS |
| VE_Peribronchial | 3 | 1 | 0 | 3 | 1 | 0.928 | 3.072 | 0.14278 | NS |
| ATI | 2 | 0 | 0 | 3 | 3 | 0.464 | 1.536 | 0.1143 | NS |
| VE_Capillary_A | 1 | 1 | 0 | 3 | 3 | 0.464 | 1.536 | 0.57789 | NS |
| ILC_A | 0 | 0 | 0 | 4 | 3 | 0 | 0 | 1 | NS |
| DC_Mature | 1 | 2 | 1 | 1 | 1 | 0.696 | 2.304 | 0.78273 | NS |
| Pericyte | 1 | 1 | 0 | 1 | 3 | 0.464 | 1.536 | 0.57789 | NS |
| SMC | 0 | 1 | 0 | 3 | 2 | 0.232 | 0.768 | 0.60849 | NS |
| Basal | 0 | 0 | 1 | 1 | 3 | 0 | 0 | 1 | NS |
| VE_Arterial | 1 | 0 | 0 | 3 | 1 | 0.232 | 0.768 | 0.26415 | NS |
| T_Regulatory | 0 | 1 | 0 | 4 | 0 | 0.232 | 0.768 | 0.60849 | NS |
| ILC_B | 0 | 1 | 0 | 2 | 1 | 0.232 | 0.768 | 0.60849 | NS |
| Club | 0 | 0 | 0 | 3 | 1 | 0 | 0 | 1 | NS |
| VE_Venous | 0 | 1 | 0 | 3 | 0 | 0.232 | 0.768 | 0.60849 | NS |

| | | | | | | | | | |
|---|---|---|---|---|---|---|---|---|---|
| PNEC | 0 | 0 | 0 | 0 | 1 | 0 | 0 | 1 | NS |

Sample:253C Disease:Control Sex:F Age:66 UMI1:945 UMI2:783 total1:27 total2:249

| Region_Celltype | X1 | X2 | Both | Unknown | LowCoverage | Expected1 | Expected2 | P_value | Direction |
|---|---|---|---|---|---|---|---|---|---|
| cMonocyte | 4 | 39 | 2 | 119 | 93 | 4.207 | 38.793 | 0.93958 | NS |
| Macrophage | 4 | 79 | 8 | 59 | 40 | 8.12 | 74.88 | 0.21905 | NS |
| Macrophage_Alveolar | 9 | 72 | 1 | 9 | 18 | 7.924 | 73.076 | 0.78223 | NS |
| T_Cytotoxic | 0 | 4 | 0 | 30 | 3 | 0.391 | 3.609 | 0.52125 | NS |
| cDC2 | 0 | 16 | 0 | 8 | 8 | 1.565 | 14.435 | 0.19954 | NS |
| NK | 0 | 1 | 0 | 13 | 6 | 0.098 | 0.902 | 0.74843 | NS |
| ncMonocyte | 1 | 5 | 0 | 6 | 7 | 0.587 | 5.413 | 0.72485 | NS |
| Lymphatic | 1 | 3 | 2 | 4 | 6 | 0.391 | 3.609 | 0.57019 | NS |
| cDC1 | 0 | 3 | 0 | 5 | 4 | 0.293 | 2.707 | 0.57856 | NS |
| VE_Arterial | 3 | 6 | 0 | 1 | 2 | 0.88 | 8.12 | 0.22441 | NS |
| VE_Capillary_B | 2 | 4 | 0 | 5 | 1 | 0.587 | 5.413 | 0.32123 | NS |
| Fibroblast | 2 | 3 | 0 | 3 | 2 | 0.489 | 4.511 | 0.26916 | NS |
| Myofibroblast | 0 | 2 | 0 | 3 | 5 | 0.196 | 1.804 | 0.65015 | NS |
| VE_Venous | 1 | 4 | 0 | 0 | 3 | 0.489 | 4.511 | 0.64998 | NS |
| T | 0 | 1 | 0 | 4 | 1 | 0.098 | 0.902 | 0.74843 | NS |
| DC_Mature | 0 | 2 | 0 | 2 | 0 | 0.196 | 1.804 | 0.65015 | NS |
| Multiplet | 0 | 1 | 1 | 2 | 0 | 0.098 | 0.902 | 0.74843 | NS |
| Ciliated | 0 | 2 | 0 | 1 | 0 | 0.196 | 1.804 | 0.65015 | NS |
| ILC_A | 0 | 0 | 0 | 3 | 0 | 0 | 0 | 1 | NS |
| PNEC | 0 | 1 | 0 | 1 | 1 | 0.098 | 0.902 | 0.74843 | NS |
| VE_Capillary_A | 0 | 0 | 0 | 3 | 0 | 0 | 0 | 1 | NS |
| B | 0 | 0 | 0 | 2 | 1 | 0 | 0 | 1 | NS |
| T_Regulatory | 0 | 0 | 0 | 2 | 1 | 0 | 0 | 1 | NS |
| Mast | 0 | 0 | 0 | 1 | 1 | 0 | 0 | 1 | NS |
| ATI | 0 | 0 | 0 | 0 | 1 | 0 | 0 | 1 | NS |
| ATII | 0 | 1 | 0 | 0 | 0 | 0.098 | 0.902 | 0.74843 | NS |
| SMC | 0 | 0 | 0 | 0 | 1 | 0 | 0 | 1 | NS |
| Goblet | 0 | 0 | 0 | 0 | 1 | 0 | 0 | 1 | NS |
| DC_Langerhans | 0 | 0 | 1 | 0 | 0 | 0 | 0 | 1 | NS |

Sample:296C Disease:Control Sex:F Age:80 UMI1:3303 UMI2:2274 total1:39 total2:1235

| Region_Celltype | X1 | X2 | Both | Unknown | LowCoverage | Expected1 | Expected2 | P_value | Direction |
|---|---|---|---|---|---|---|---|---|---|
| Macrophage_Alveolar | 10 | 516 | 19 | 91 | 304 | 16.102 | 509.898 | 0.22648 | NS |
| Macrophage | 9 | 358 | 20 | 28 | 113 | 11.235 | 355.765 | 0.61442 | NS |
| cMonocyte | 10 | 294 | 3 | 29 | 130 | 9.306 | 294.694 | 0.8725 | NS |
| VE_Capillary_B | 0 | 3 | 0 | 26 | 22 | 0.092 | 2.908 | 0.76007 | NS |
| Lymphatic | 1 | 2 | 0 | 27 | 17 | 0.092 | 2.908 | 0.33658 | NS |
| ATII | 4 | 4 | 2 | 7 | 21 | 0.245 | 7.755 | 0.033474 | NS |
| ncMonocyte | 1 | 23 | 0 | 2 | 4 | 0.735 | 23.265 | 0.83743 | NS |

| | | | | | | | | | |
|---|---|---|---|---|---|---|---|---|---|
| Multiplet | 0 | 14 | 2 | 4 | 7 | 0.429 | 13.571 | 0.50943 | NS |
| NK | 0 | 4 | 0 | 5 | 15 | 0.122 | 3.878 | 0.72436 | NS |
| T | 0 | 4 | 0 | 4 | 16 | 0.122 | 3.878 | 0.72436 | NS |
| Fibroblast | 0 | 1 | 0 | 6 | 6 | 0.031 | 0.969 | 0.86004 | NS |
| T_Cytotoxic | 0 | 1 | 0 | 4 | 7 | 0.031 | 0.969 | 0.86004 | NS |
| VE_Capillary_A | 1 | 0 | 0 | 8 | 2 | 0.031 | 0.969 | 0.1702 | NS |
| Club | 0 | 0 | 1 | 4 | 2 | 0 | 0 | 1 | NS |
| Ciliated | 0 | 1 | 0 | 1 | 4 | 0.031 | 0.969 | 0.86004 | NS |
| cDC2 | 0 | 5 | 0 | 0 | 1 | 0.153 | 4.847 | 0.69339 | NS |
| Myofibroblast | 2 | 0 | 0 | 1 | 3 | 0.061 | 1.939 | 0.052418 | NS |
| VE_Venous | 0 | 0 | 0 | 2 | 3 | 0 | 0 | 1 | NS |
| VE_Peribronchial | 0 | 0 | 2 | 1 | 2 | 0 | 0 | 1 | NS |
| SMC | 0 | 0 | 0 | 3 | 1 | 0 | 0 | 1 | NS |
| ATI | 1 | 0 | 0 | 1 | 2 | 0.031 | 0.969 | 0.1702 | NS |
| VE_Arterial | 0 | 1 | 0 | 1 | 2 | 0.031 | 0.969 | 0.86004 | NS |
| cDC1 | 0 | 2 | 0 | 1 | 0 | 0.061 | 1.939 | 0.80309 | NS |
| Mast | 0 | 1 | 0 | 0 | 2 | 0.031 | 0.969 | 0.86004 | NS |
| T_Regulatory | 0 | 0 | 1 | 1 | 1 | 0 | 0 | 1 | NS |
| Mesothelial | 0 | 0 | 0 | 0 | 2 | 0 | 0 | 1 | NS |
| B | 0 | 0 | 0 | 1 | 1 | 0 | 0 | 1 | NS |
| B_Plasma | 0 | 0 | 0 | 1 | 0 | 0 | 0 | 1 | NS |
| Goblet | 0 | 1 | 0 | 0 | 0 | 0.031 | 0.969 | 0.86004 | NS |
| DC_Mature | 0 | 0 | 0 | 1 | 0 | 0 | 0 | 1 | NS |

Sample:396C Disease:Control Sex:F Age:37 UMI1:1268 UMI2:739 total1:75 total2:179

| Region_Celltype | X1 | X2 | Both | Unknown | LowCoverage | Expected1 | Expected2 | P_value | Direction |
|---|---|---|---|---|---|---|---|---|---|
| Macrophage_Alveolar | 35 | 85 | 12 | 65 | 143 | 35.433 | 84.567 | 0.95105 | NS |
| Macrophage | 30 | 77 | 8 | 36 | 98 | 31.594 | 75.406 | 0.80975 | NS |
| ATII | 4 | 11 | 0 | 26 | 42 | 4.429 | 10.571 | 0.86162 | NS |
| cDC2 | 5 | 0 | 0 | 8 | 4 | 1.476 | 3.524 | 0.019672 | NS |
| Multiplet | 0 | 4 | 0 | 7 | 6 | 1.181 | 2.819 | 0.23914 | NS |
| Lymphatic | 0 | 0 | 0 | 6 | 1 | 0 | 0 | 1 | NS |
| cMonocyte | 0 | 1 | 0 | 3 | 1 | 0.295 | 0.705 | 0.55615 | NS |
| ATI | 0 | 0 | 0 | 1 | 2 | 0 | 0 | 1 | NS |
| Ciliated | 0 | 0 | 0 | 2 | 0 | 0 | 0 | 1 | NS |
| NK | 0 | 0 | 0 | 1 | 1 | 0 | 0 | 1 | NS |
| Club | 0 | 0 | 0 | 0 | 2 | 0 | 0 | 1 | NS |
| ncMonocyte | 1 | 0 | 0 | 1 | 0 | 0.295 | 0.705 | 0.29688 | NS |
| Fibroblast | 0 | 0 | 0 | 1 | 0 | 0 | 0 | 1 | NS |
| VE_Venous | 0 | 0 | 0 | 1 | 0 | 0 | 0 | 1 | NS |
| B_Plasma | 0 | 0 | 0 | 1 | 0 | 0 | 0 | 1 | NS |
| Mesothelial | 0 | 0 | 0 | 1 | 0 | 0 | 0 | 1 | NS |
| SMC | 0 | 0 | 0 | 1 | 0 | 0 | 0 | 1 | NS |

| | | | | | | | | | |
|---|---|---|---|---|---|---|---|---|---|
| Goblet | 0 | 0 | 0 | 0 | 1 | 0 | 0 | 1 | NS |
| DC_Langerhans | 0 | 0 | 0 | 1 | 0 | 0 | 0 | 1 | NS |
| Basal | 0 | 1 | 0 | 0 | 0 | 0.295 | 0.705 | 0.55615 | NS |
| VE_Arterial | 0 | 0 | 0 | 1 | 0 | 0 | 0 | 1 | NS |
| VE_Capillary_B | 0 | 0 | 0 | 1 | 0 | 0 | 0 | 1 | NS |

Sample:439C Disease:Control Sex:F Age:66 UMI1:7197 UMI2:3833 total1:226 total2:305

| Region_Celltype | X1 | X2 | Both | Unknown | LowCoverage | Expected1 | Expected2 | P_value | Direction |
|---|---|---|---|---|---|---|---|---|---|
| Macrophage_Alveolar | 96 | 164 | 12 | 1330 | 224 | 110.659 | 149.341 | 0.18897 | NS |
| Macrophage | 46 | 57 | 3 | 547 | 87 | 43.838 | 59.162 | 0.76132 | NS |
| T | 21 | 14 | 3 | 259 | 32 | 14.896 | 20.104 | 0.14442 | NS |
| ncMonocyte | 16 | 22 | 2 | 177 | 26 | 16.173 | 21.827 | 0.96791 | NS |
| cMonocyte | 11 | 15 | 0 | 166 | 20 | 11.066 | 14.934 | 0.98524 | NS |
| T_Cytotoxic | 13 | 7 | 1 | 118 | 22 | 8.512 | 11.488 | 0.15467 | NS |
| NK | 7 | 7 | 0 | 72 | 9 | 5.959 | 8.041 | 0.69305 | NS |
| cDC2 | 3 | 8 | 1 | 38 | 4 | 4.682 | 6.318 | 0.45197 | NS |
| Multiplet | 4 | 5 | 1 | 27 | 7 | 3.831 | 5.169 | 0.93577 | NS |
| Ciliated | 2 | 0 | 0 | 17 | 2 | 0.851 | 1.149 | 0.20426 | NS |
| ILC_A | 0 | 2 | 0 | 11 | 4 | 0.851 | 1.149 | 0.2984 | NS |
| B | 1 | 0 | 0 | 10 | 4 | 0.426 | 0.574 | 0.36936 | NS |
| ATII | 0 | 1 | 0 | 8 | 2 | 0.426 | 0.574 | 0.46216 | NS |
| cDC1 | 2 | 0 | 0 | 7 | 0 | 0.851 | 1.149 | 0.20426 | NS |
| Lymphatic | 0 | 0 | 0 | 7 | 1 | 0 | 0 | 1 | NS |
| Fibroblast | 0 | 1 | 0 | 6 | 0 | 0.426 | 0.574 | 0.46216 | NS |
| Club | 0 | 0 | 0 | 6 | 0 | 0 | 0 | 1 | NS |
| T_Regulatory | 2 | 1 | 0 | 1 | 1 | 1.277 | 1.723 | 0.55319 | NS |
| VE_Venous | 0 | 0 | 0 | 4 | 1 | 0 | 0 | 1 | NS |
| DC_Mature | 0 | 0 | 0 | 3 | 1 | 0 | 0 | 1 | NS |
| B_Plasma | 0 | 0 | 0 | 3 | 1 | 0 | 0 | 1 | NS |
| VE_Capillary_B | 0 | 1 | 0 | 2 | 0 | 0.426 | 0.574 | 0.46216 | NS |
| VE_Arterial | 0 | 0 | 0 | 2 | 1 | 0 | 0 | 1 | NS |
| ATI | 0 | 0 | 0 | 2 | 1 | 0 | 0 | 1 | NS |
| Myofibroblast | 1 | 0 | 0 | 1 | 0 | 0.426 | 0.574 | 0.36936 | NS |
| ILC_B | 0 | 0 | 0 | 2 | 0 | 0 | 0 | 1 | NS |
| VE_Capillary_A | 0 | 0 | 0 | 2 | 0 | 0 | 0 | 1 | NS |
| Mesothelial | 1 | 0 | 0 | 0 | 0 | 0.426 | 0.574 | 0.36936 | NS |
| Goblet | 0 | 0 | 0 | 0 | 1 | 0 | 0 | 1 | NS |

Sample:454C Disease:Control Sex:F Age:48 UMI1:1163 UMI2:540 total1:21 total2:32

| Region_Celltype | X1 | X2 | Both | Unknown | LowCoverage | Expected1 | Expected2 | P_value | Direction |
|---|---|---|---|---|---|---|---|---|---|
| Macrophage | 11 | 14 | 2 | 215 | 52 | 9.906 | 15.094 | 0.7537 | NS |
| cDC2 | 2 | 5 | 0 | 40 | 10 | 2.774 | 4.226 | 0.66273 | NS |
| ncMonocyte | 1 | 4 | 2 | 20 | 27 | 1.981 | 3.019 | 0.4976 | NS |

| | | | | | | | | | |
|---|---|---|---|---|---|---|---|---|---|
| cMonocyte | 1 | 5 | 1 | 20 | 14 | 2.377 | 3.623 | 0.37661 | NS |
| Macrophage_Alveolar | 2 | 0 | 0 | 18 | 7 | 0.792 | 1.208 | 0.18845 | NS |
| ATII | 1 | 2 | 0 | 5 | 6 | 1.189 | 1.811 | 0.87287 | NS |
| cDC1 | 1 | 1 | 0 | 6 | 3 | 0.792 | 1.208 | 0.8347 | NS |
| DC_Mature | 2 | 0 | 0 | 7 | 1 | 0.792 | 1.208 | 0.18845 | NS |
| Lymphatic | 0 | 0 | 0 | 2 | 5 | 0 | 0 | 1 | NS |
| Multiplet | 0 | 0 | 0 | 5 | 0 | 0 | 0 | 1 | NS |
| T_Cytotoxic | 0 | 0 | 0 | 2 | 2 | 0 | 0 | 1 | NS |
| ATI | 0 | 0 | 0 | 3 | 0 | 0 | 0 | 1 | NS |
| VE_Arterial | 0 | 0 | 0 | 0 | 2 | 0 | 0 | 1 | NS |
| NK | 0 | 0 | 0 | 2 | 0 | 0 | 0 | 1 | NS |
| Club | 0 | 0 | 0 | 0 | 2 | 0 | 0 | 1 | NS |
| pDC | 0 | 0 | 0 | 2 | 0 | 0 | 0 | 1 | NS |
| B_Plasma | 0 | 1 | 0 | 0 | 0 | 0.396 | 0.604 | 0.4821 | NS |
| VE_Venous | 0 | 0 | 0 | 0 | 1 | 0 | 0 | 1 | NS |
| B | 0 | 0 | 0 | 0 | 1 | 0 | 0 | 1 | NS |
| Myofibroblast | 0 | 0 | 0 | 1 | 0 | 0 | 0 | 1 | NS |
| T | 0 | 0 | 0 | 0 | 1 | 0 | 0 | 1 | NS |
