## Supplemental Table 4 for "Single-cell X-chromosome inactivation analysis links biased chimerism to differential gene expression and epigenetic erosion"

IB5431, haplotyped: 40961, With Celltype: 40961

| Region_Celltype | X1 | X2 | Both | Unknown | LowCoverage | Expected1 | Expected2 | P_value | Direction |
|---|---|---|---|---|---|---|---|---|---|
| OTC_Unassigned | 927 | 1072 | 414 | 1374 | 4841 | 878.39 | 1120.61 | 0.12239 | NS |
| CB_Unassigned | 194 | 354 | 156 | 604 | 2345 | 240.799 | 307.201 | 0.0038576 | Outlier |
| OC_astrocytes | 630 | 1134 | 117 | 399 | 1069 | 775.128 | 988.872 | 6.004e-07 | Outlier |
| CB_Bergmann1 | 326 | 576 | 65 | 513 | 1587 | 396.352 | 505.648 | 0.00072359 | Outlier |
| OTC_excitatory | 833 | 988 | 87 | 111 | 695 | 800.175 | 1020.825 | 0.27409 | NS |
| OTC_astrocytes | 684 | 837 | 121 | 243 | 731 | 668.35 | 852.65 | 0.56799 | NS |
| CB_Neurins_In_Purkinje | 615 | 877 | 94 | 173 | 770 | 655.607 | 836.393 | 0.13274 | NS |
| OC_Unassigned | 46 | 68 | 11 | 781 | 1605 | 50.093 | 63.907 | 0.58302 | NS |
| OTC_inhibitory | 641 | 601 | 72 | 164 | 917 | 545.753 | 696.247 | 0.00013029 | Uniform |
| OC_microglia | 147 | 73 | 23 | 769 | 1145 | 96.671 | 123.329 | 1.3883e-06 | Uniform |
| OTC_oligodendrocytes | 191 | 198 | 36 | 249 | 794 | 170.932 | 218.068 | 0.14919 | NS |
| OTC_microglia | 137 | 68 | 27 | 457 | 737 | 90.08 | 114.92 | 3.1385e-06 | Uniform |
| CB_Neurons_Ex | 71 | 101 | 25 | 80 | 426 | 75.579 | 96.421 | 0.61758 | NS |
| CB_Astrocytes | 61 | 97 | 8 | 86 | 231 | 69.428 | 88.572 | 0.33557 | NS |
| CB_Neurons_In | 93 | 142 | 13 | 30 | 172 | 103.262 | 131.738 | 0.33712 | NS |
| OC_oligodendrocytes | 77 | 130 | 21 | 68 | 151 | 90.959 | 116.041 | 0.16237 | NS |
| CB_Macrophages | 20 | 6 | 3 | 109 | 198 | 11.425 | 14.575 | 0.015022 | NS |
| OC_endothelial | 39 | 50 | 11 | 39 | 148 | 39.108 | 49.892 | 0.98699 | NS |
| OTC_OPCs | 26 | 38 | 8 | 38 | 138 | 28.123 | 35.877 | 0.70412 | NS |
| OC_pericytes | 35 | 36 | 13 | 30 | 117 | 31.198 | 39.802 | 0.5225 | NS |
| OTC_endothelial | 34 | 28 | 9 | 41 | 102 | 27.244 | 34.756 | 0.22492 | NS |
| CB_Neurons? | 79 | 71 | 3 | 9 | 34 | 65.912 | 84.088 | 0.1305 | NS |
| OC_neuron | 51 | 69 | 6 | 12 | 42 | 52.73 | 67.27 | 0.82167 | NS |
| CB_Oligodendrocytes | 15 | 17 | 2 | 27 | 88 | 14.061 | 17.939 | 0.81368 | NS |
| OTC_pericytes | 31 | 18 | 9 | 26 | 59 | 21.531 | 27.469 | 0.055118 | NS |
| CB_Fibroblasts | 6 | 13 | 0 | 21 | 71 | 8.349 | 10.651 | 0.43187 | NS |
| CB_Endothelial | 2 | 9 | 3 | 21 | 71 | 4.834 | 6.166 | 0.19172 | NS |
| OTC_neuron | 23 | 26 | 9 | 7 | 26 | 21.531 | 27.469 | 0.76574 | NS |
| CB_Mural | 1 | 4 | 1 | 9 | 36 | 2.197 | 2.803 | 0.41696 | NS |
| CB_Lymphocytes | 0 | 0 | 0 | 3 | 10 | 0 | 0 | 1 | NS |
| CB_OPCs | 3 | 1 | 0 | 1 | 2 | 1.758 | 2.242 | 0.37097 | NS |
| CB_Neurons | 0 | 1 | 0 | 0 | 1 | 0.439 | 0.561 | 0.453 | NS |

IB5508, haplotyped: 35393, With Celltype: 35393

| Region_Celltype | X1 | X2 | Both | Unknown | LowCoverage | Expected1 | Expected2 | P_value | Direction |
|---|---|---|---|---|---|---|---|---|---|
| OTC_Unassigned | 292 | 301 | 97 | 1390 | 4643 | 295.909 | 297.091 | 0.82042 | NS |
| CB_Unassigned | 330 | 369 | 234 | 573 | 2135 | 348.803 | 350.197 | 0.31432 | NS |
| OTC_astrocytes | 852 | 748 | 161 | 405 | 1130 | 798.404 | 801.596 | 0.057983 | NS |
| CB_Bergmann1 | 629 | 760 | 93 | 342 | 1308 | 693.115 | 695.885 | 0.014862 | NS |
| OC_Unassigned | 81 | 77 | 22 | 589 | 1850 | 78.842 | 79.158 | 0.80819 | NS |
| CB_Neurins_In_Purkinje | 725 | 721 | 85 | 114 | 483 | 721.558 | 724.442 | 0.89814 | NS |

| | | | | | | | | | |
|---|---|---|---|---|---|---|---|---|---|
| OC_astrocytes | 474 | 403 | 99 | 237 | 721 | 437.625 | 439.375 | 0.082139 | NS |
| OTC_inhibitory | 211 | 203 | 24 | 230 | 1087 | 206.587 | 207.413 | 0.75905 | NS |
| OTC_microglia | 102 | 53 | 25 | 502 | 778 | 77.345 | 77.655 | 0.0045716 | Uniform |
| OTC_oligodendrocytes | 135 | 216 | 57 | 224 | 794 | 175.15 | 175.85 | 0.0022773 | Outlier |
| OTC_excitatory | 186 | 205 | 14 | 124 | 882 | 195.11 | 195.89 | 0.51456 | NS |
| CB_Neurons_Ex | 155 | 180 | 25 | 164 | 643 | 167.166 | 167.834 | 0.34685 | NS |
| CB_Astrocytes | 194 | 236 | 18 | 103 | 448 | 214.571 | 215.429 | 0.16011 | NS |
| OC_microglia | 84 | 54 | 16 | 288 | 505 | 68.862 | 69.138 | 0.0668 | NS |
| CB_Neurons_In | 147 | 125 | 18 | 33 | 151 | 135.729 | 136.271 | 0.33342 | NS |
| OC_oligodendrocytes | 96 | 66 | 28 | 53 | 167 | 80.838 | 81.162 | 0.090698 | NS |
| CB_Macrophages | 22 | 15 | 4 | 98 | 175 | 18.463 | 18.537 | 0.40884 | NS |
| CB_Oligodendrocytes | 40 | 38 | 6 | 40 | 126 | 38.922 | 39.078 | 0.86297 | NS |
| OTC_pericytes | 22 | 44 | 12 | 42 | 125 | 32.934 | 33.066 | 0.053517 | NS |
| OC_neuron | 61 | 61 | 16 | 10 | 38 | 60.878 | 61.122 | 0.98757 | NS |
| OTC_endothelial | 18 | 13 | 5 | 32 | 78 | 15.469 | 15.531 | 0.51899 | NS |
| CB_Fibroblasts | 30 | 7 | 2 | 17 | 80 | 18.463 | 18.537 | 0.0047862 | Uniform |
| OC_pericytes | 17 | 29 | 7 | 20 | 63 | 22.954 | 23.046 | 0.21043 | NS |
| OTC_OPCs | 16 | 11 | 0 | 18 | 88 | 13.473 | 13.527 | 0.48979 | NS |
| OTC_neuron | 30 | 34 | 8 | 3 | 23 | 31.936 | 32.064 | 0.73201 | NS |
| OC_endothelial | 19 | 8 | 4 | 14 | 40 | 13.473 | 13.527 | 0.12451 | NS |
| CB_Neurons? | 25 | 39 | 3 | 1 | 1 | 31.936 | 32.064 | 0.21732 | NS |
| CB_Endothelial | 3 | 4 | 0 | 8 | 18 | 3.493 | 3.507 | 0.79161 | NS |
| CB_Mural | 5 | 3 | 0 | 5 | 8 | 3.992 | 4.008 | 0.61152 | NS |
| CB_Lymphocytes | 3 | 1 | 1 | 2 | 10 | 1.996 | 2.004 | 0.46355 | NS |
| CB_OPCs | 0 | 0 | 0 | 1 | 0 | 0 | 0 | 1 | NS |
| CB_Bergmann2 | 0 | 0 | 0 | 1 | 0 | 0 | 0 | 1 | NS |

IB5691, haplotyped: 41447, With Celltype: 41447

| Region_Celltype | X1 | X2 | Both | Unknown | LowCoverage | Expected1 | Expected2 | P_value | Direction |
|---|---|---|---|---|---|---|---|---|---|
| CB_Neurons_Ex | 259 | 621 | 141 | 3703 | 3293 | 320.564 | 559.436 | 0.0017928 | Outlier |
| CB_Unassigned | 51 | 89 | 10 | 4110 | 1596 | 50.999 | 89.001 | 0.99988 | NS |
| OC_Unassigned | 96 | 150 | 13 | 2420 | 2081 | 89.612 | 156.388 | 0.55241 | NS |
| OC_astrocytes | 314 | 505 | 62 | 785 | 1335 | 298.343 | 520.657 | 0.42395 | NS |
| OTC_astrocytes | 329 | 523 | 46 | 725 | 1139 | 310.364 | 541.636 | 0.35113 | NS |
| OTC_Unassigned | 65 | 66 | 7 | 1473 | 1139 | 47.72 | 83.28 | 0.03107 | NS |
| OTC_oligodendrocytes | 178 | 257 | 51 | 463 | 998 | 158.461 | 276.539 | 0.17375 | NS |
| OC_microglia | 73 | 142 | 18 | 993 | 707 | 78.32 | 136.68 | 0.59115 | NS |
| OTC_microglia | 69 | 114 | 15 | 951 | 640 | 66.663 | 116.337 | 0.80031 | NS |
| OC_oligodendrocytes | 121 | 204 | 35 | 319 | 659 | 118.39 | 206.61 | 0.83193 | NS |
| CB_Bergmann1 | 69 | 87 | 13 | 456 | 605 | 56.827 | 99.173 | 0.16008 | NS |
| CB_Neurins_In_Purkinje | 165 | 220 | 36 | 156 | 404 | 140.247 | 244.753 | 0.068206 | NS |
| OC_neuron | 126 | 316 | 34 | 131 | 245 | 161.011 | 280.989 | 0.011912 | NS |
| OC_endothelial | 24 | 68 | 9 | 365 | 311 | 33.514 | 58.486 | 0.13028 | NS |

| | | | | | | | | | |
|---|---|---|---|---|---|---|---|---|---|
| CB_Astrocytes | 28 | 47 | 14 | 171 | 260 | 27.321 | 47.679 | 0.90849 | NS |
| CB_Neurons_In | 39 | 71 | 21 | 122 | 194 | 40.071 | 69.929 | 0.88044 | NS |
| CB_Oligodendrocytes | 10 | 24 | 10 | 126 | 166 | 12.385 | 21.615 | 0.53817 | NS |
| OTC_endothelial | 10 | 14 | 3 | 147 | 137 | 8.743 | 15.257 | 0.70989 | NS |
| OTC_neuron | 51 | 127 | 11 | 37 | 82 | 64.841 | 113.159 | 0.11741 | NS |
| CB_Neurons | 20 | 26 | 13 | 103 | 132 | 16.757 | 29.243 | 0.48998 | NS |
| OC_pericytes | 9 | 17 | 6 | 98 | 115 | 9.471 | 16.529 | 0.89139 | NS |
| CB_Endothelial | 4 | 9 | 1 | 92 | 69 | 4.736 | 8.264 | 0.76004 | NS |
| CB_Macrophages | 3 | 15 | 5 | 88 | 62 | 6.557 | 11.443 | 0.17943 | NS |
| CB_OPCs | 24 | 30 | 7 | 33 | 53 | 19.671 | 34.329 | 0.396 | NS |
| CB_Fibroblasts | 9 | 5 | 1 | 56 | 72 | 5.1 | 8.9 | 0.14044 | NS |
| CB_Neurons? | 12 | 22 | 3 | 45 | 50 | 12.385 | 21.615 | 0.92236 | NS |
| OTC_pericytes | 6 | 4 | 0 | 43 | 32 | 3.643 | 6.357 | 0.29149 | NS |
| CB_Bergmann2 | 4 | 6 | 1 | 34 | 32 | 3.643 | 6.357 | 0.86942 | NS |
| CB_Mural | 2 | 7 | 0 | 24 | 19 | 3.278 | 5.722 | 0.50802 | NS |
| CB_Lymphocytes | 0 | 1 | 0 | 6 | 2 | 0.364 | 0.636 | 0.50453 | NS |

IB5869, haplotyped: 44551, With Celltype: 44551

| Region_Celltype | X1 | X2 | Both | Unknown | LowCoverage | Expected1 | Expected2 | P_value | Direction |
|---|---|---|---|---|---|---|---|---|---|
| OTC_Unassigned | 126 | 353 | 51 | 2090 | 3436 | 153.786 | 325.214 | 0.048347 | NS |
| CB_Unassigned | 191 | 399 | 119 | 1673 | 2611 | 189.423 | 400.577 | 0.92178 | NS |
| OC_Unassigned | 64 | 149 | 42 | 1590 | 2660 | 68.385 | 144.615 | 0.64619 | NS |
| CB_Bergmann1 | 364 | 619 | 102 | 934 | 1985 | 315.599 | 667.401 | 0.021718 | NS |
| OC_astrocytes | 607 | 742 | 133 | 801 | 1425 | 433.105 | 915.895 | 6.0512e-12 | Uniform |
| OTC_astrocytes | 399 | 1013 | 115 | 688 | 1384 | 453.332 | 958.668 | 0.025931 | NS |
| OC_microglia | 85 | 426 | 42 | 1095 | 1459 | 164.06 | 346.94 | 8.3885e-09 | Outlier |
| OTC_microglia | 55 | 260 | 30 | 1199 | 1415 | 101.133 | 213.867 | 2.0717e-05 | Outlier |
| CB_Neurins_In_Purkinje | 360 | 684 | 71 | 286 | 721 | 335.183 | 708.817 | 0.24914 | NS |
| OTC_excitatory | 97 | 333 | 12 | 351 | 930 | 138.054 | 291.946 | 0.0016822 | Outlier |
| CB_Astrocytes | 143 | 319 | 46 | 360 | 685 | 148.328 | 313.672 | 0.70598 | NS |
| OC_endothelial | 73 | 86 | 36 | 269 | 403 | 51.048 | 107.952 | 0.011611 | NS |
| CB_Macrophages | 19 | 88 | 8 | 276 | 380 | 34.353 | 72.647 | 0.015267 | NS |
| OTC_inhibitory | 32 | 103 | 6 | 177 | 418 | 43.343 | 91.657 | 0.1238 | NS |
| OC_oligodendrocytes | 52 | 83 | 20 | 151 | 299 | 43.343 | 91.657 | 0.2703 | NS |
| OC_neuron | 76 | 130 | 48 | 83 | 145 | 66.138 | 139.862 | 0.30672 | NS |
| CB_Neurons_In | 40 | 70 | 12 | 80 | 159 | 35.316 | 74.684 | 0.50573 | NS |
| OTC_neuron | 36 | 67 | 23 | 49 | 135 | 33.069 | 69.931 | 0.66531 | NS |
| OTC_endothelial | 35 | 18 | 9 | 94 | 153 | 17.016 | 35.984 | 0.00047562 | Uniform |
| OC_pericytes | 19 | 30 | 12 | 80 | 141 | 15.732 | 33.268 | 0.49008 | NS |
| CB_Fibroblasts | 9 | 38 | 5 | 66 | 149 | 15.09 | 31.91 | 0.15023 | NS |
| CB_Neurons_Ex | 11 | 29 | 26 | 63 | 136 | 12.842 | 27.158 | 0.65248 | NS |
| OTC_oligodendrocytes | 9 | 26 | 8 | 86 | 127 | 11.237 | 23.763 | 0.55534 | NS |
| CB_Endothelial | 7 | 27 | 3 | 73 | 126 | 10.916 | 23.084 | 0.28103 | NS |

| | | | | | | | | | |
|---|---|---|---|---|---|---|---|---|---|
| OTC_pericytes | 3 | 37 | 1 | 60 | 101 | 12.842 | 27.158 | 0.0057579 | Outlier |
| CB_Neurons? | 11 | 36 | 5 | 8 | 25 | 15.09 | 31.91 | 0.34619 | NS |
| CB_Mural | 3 | 10 | 0 | 11 | 45 | 4.174 | 8.826 | 0.60656 | NS |
| CB_Oligodendrocytes | 2 | 7 | 3 | 7 | 24 | 2.89 | 6.11 | 0.63739 | NS |
| CB_OPCs | 1 | 10 | 1 | 8 | 14 | 3.532 | 7.468 | 0.182 | NS |
| CB_Lymphocytes | 0 | 1 | 0 | 11 | 17 | 0.321 | 0.679 | 0.53629 | NS |
| OTC_OPCs | 0 | 1 | 0 | 4 | 6 | 0.321 | 0.679 | 0.53629 | NS |
| CB_Bergmann2? | 0 | 0 | 0 | 1 | 0 | 0 | 0 | 1 | NS |
| CB_Neurons | 0 | 0 | 0 | 0 | 1 | 0 | 0 | 1 | NS |

IB5888, haplotyped: 21471, With Celltype: 21471

| Region_Celltype | X1 | X2 | Both | Unknown | LowCoverage | Expected1 | Expected2 | P_value | Direction |
|---|---|---|---|---|---|---|---|---|---|
| OC_astrocytes | 234 | 222 | 23 | 893 | 1314 | 208.437 | 247.563 | 0.090327 | NS |
| OTC_astrocytes | 211 | 191 | 24 | 837 | 1262 | 183.754 | 218.246 | 0.054594 | NS |
| CB_Bergmann1 | 93 | 185 | 13 | 777 | 1194 | 127.074 | 150.926 | 0.0031272 | Outlier |
| OTC_microglia | 74 | 57 | 8 | 976 | 926 | 59.88 | 71.12 | 0.080969 | NS |
| CB_Unassigned | 58 | 116 | 21 | 689 | 941 | 79.535 | 94.465 | 0.018213 | NS |
| CB_Neurins_In_Purkinje | 175 | 292 | 17 | 416 | 846 | 213.465 | 253.535 | 0.010661 | NS |
| OTC_Unassigned | 28 | 43 | 6 | 705 | 800 | 32.454 | 38.546 | 0.44969 | NS |
| OC_Unassigned | 19 | 30 | 0 | 705 | 744 | 22.398 | 26.602 | 0.48713 | NS |
| OC_microglia | 49 | 27 | 3 | 745 | 655 | 34.74 | 41.26 | 0.020049 | NS |
| OC_oligodendrocytes | 88 | 67 | 14 | 211 | 457 | 70.85 | 84.15 | 0.051336 | NS |
| CB_Astrocytes | 21 | 36 | 1 | 184 | 224 | 26.055 | 30.945 | 0.33626 | NS |
| OTC_oligodendrocytes | 29 | 25 | 4 | 143 | 237 | 24.683 | 29.317 | 0.40612 | NS |
| CB_Macrophages | 8 | 8 | 2 | 167 | 164 | 7.314 | 8.686 | 0.80808 | NS |
| CB_Neurons_Ex | 10 | 4 | 3 | 100 | 145 | 6.399 | 7.601 | 0.16717 | NS |
| CB_Neurons_In | 21 | 22 | 1 | 86 | 118 | 19.655 | 23.345 | 0.77148 | NS |
| OC_neuron | 39 | 34 | 5 | 55 | 85 | 33.368 | 39.632 | 0.35123 | NS |
| OTC_endothelial | 5 | 9 | 2 | 70 | 104 | 6.399 | 7.601 | 0.59038 | NS |
| OTC_neuron | 20 | 12 | 4 | 50 | 71 | 14.627 | 17.373 | 0.17774 | NS |
| OC_pericytes | 0 | 11 | 1 | 62 | 72 | 5.028 | 5.972 | 0.010681 | NS |
| OC_endothelial | 6 | 7 | 2 | 52 | 62 | 5.942 | 7.058 | 0.98188 | NS |
| OTC_pericytes | 3 | 4 | 0 | 35 | 51 | 3.2 | 3.8 | 0.91443 | NS |
| CB_Neurons? | 7 | 15 | 0 | 27 | 43 | 10.056 | 11.944 | 0.34432 | NS |
| CB_Oligodendrocytes | 4 | 4 | 2 | 20 | 45 | 3.657 | 4.343 | 0.86363 | NS |
| CB_Fibroblasts | 1 | 1 | 0 | 13 | 21 | 0.914 | 1.086 | 0.93156 | NS |
| CB_Endothelial | 1 | 1 | 0 | 13 | 20 | 0.914 | 1.086 | 0.93156 | NS |
| CB_OPCs | 0 | 5 | 0 | 9 | 18 | 2.285 | 2.715 | 0.085211 | NS |
| CB_Mural | 0 | 0 | 0 | 8 | 11 | 0 | 0 | 1 | NS |
| CB_Lymphocytes | 0 | 1 | 0 | 0 | 3 | 0.457 | 0.543 | 0.44145 | NS |
| CB_Bergmann2 | 0 | 1 | 0 | 0 | 0 | 0.457 | 0.543 | 0.44145 | NS |

IB6001, haplotyped: 46118, With Celltype: 46118

| Region_Celltype | X1 | X2 | Both | Unknown | LowCoverage | Expected1 | Expected2 | P_value | Direction |
|---|---|---|---|---|---|---|---|---|---|
| CB_Unassigned | 1340 | 1502 | 387 | 3132 | 6151 | 1238.645 | 1603.355 | 0.0069265 | Uniform |
| OTC_microglia | 392 | 307 | 73 | 3140 | 2345 | 304.649 | 394.351 | 2.976e-06 | Uniform |
| OC_Unassigned | 96 | 128 | 19 | 2174 | 2200 | 97.627 | 126.373 | 0.87667 | NS |
| OTC_Unassigned | 94 | 98 | 14 | 2397 | 1849 | 83.68 | 108.32 | 0.29089 | NS |
| OC_astrocytes | 389 | 940 | 107 | 847 | 1700 | 579.226 | 749.774 | 1.7562e-14 | Outlier |
| OC_microglia | 196 | 143 | 30 | 1846 | 1465 | 147.748 | 191.252 | 0.0002101 | Uniform |
| OTC_endothelial | 222 | 239 | 48 | 888 | 1142 | 200.92 | 260.08 | 0.16356 | NS |
| OTC_astrocytes | 265 | 496 | 43 | 256 | 548 | 331.671 | 429.329 | 0.00046441 | Outlier |
| OC_endothelial | 100 | 150 | 31 | 494 | 752 | 108.959 | 141.041 | 0.4166 | NS |
| CB_Macrophages | 38 | 34 | 8 | 509 | 393 | 31.38 | 40.62 | 0.26958 | NS |
| CB_Neurins_In_Purkinje | 162 | 207 | 16 | 94 | 302 | 160.823 | 208.177 | 0.93042 | NS |
| CB_Astrocytes | 64 | 83 | 11 | 105 | 228 | 64.068 | 82.932 | 0.99363 | NS |
| CB_Bergmann1 | 52 | 63 | 5 | 97 | 231 | 50.121 | 64.879 | 0.80309 | NS |
| CB_Neurons_Ex | 27 | 42 | 34 | 96 | 192 | 30.073 | 38.927 | 0.59534 | NS |
| OTC_pericytes | 31 | 34 | 11 | 134 | 166 | 28.329 | 36.671 | 0.63817 | NS |
| OC_pericytes | 19 | 35 | 7 | 129 | 171 | 23.535 | 30.465 | 0.37178 | NS |
| OTC_oligodendrocytes | 29 | 46 | 7 | 119 | 153 | 32.688 | 42.312 | 0.5406 | NS |
| CB_Endothelial | 12 | 7 | 1 | 61 | 95 | 8.281 | 10.719 | 0.22651 | NS |
| CB_Fibroblasts | 23 | 4 | 1 | 34 | 84 | 11.768 | 15.232 | 0.0014128 | Uniform |
| OC_neuron | 24 | 59 | 10 | 12 | 27 | 36.174 | 46.826 | 0.049343 | NS |
| OTC_neuron | 32 | 42 | 13 | 9 | 13 | 32.252 | 41.748 | 0.96669 | NS |
| OC_oligodendrocytes | 3 | 11 | 1 | 22 | 54 | 6.102 | 7.898 | 0.21078 | NS |
| CB_Mural | 0 | 3 | 0 | 12 | 28 | 1.308 | 1.692 | 0.19601 | NS |
| CB_Lymphocytes | 0 | 1 | 0 | 14 | 15 | 0.436 | 0.564 | 0.45536 | NS |
| CB_Neurons_In | 6 | 6 | 0 | 5 | 11 | 5.23 | 6.77 | 0.75277 | NS |
| CB_Oligodendrocytes | 0 | 1 | 0 | 0 | 1 | 0.436 | 0.564 | 0.45536 | NS |
| CB_OPCs | 1 | 1 | 0 | 0 | 0 | 0.872 | 1.128 | 0.89768 | NS |

IB6125, haplotyped: 40302, With Celltype: 40302

| Region_Celltype | X1 | X2 | Both | Unknown | LowCoverage | Expected1 | Expected2 | P_value | Direction |
|---|---|---|---|---|---|---|---|---|---|
| CB_Unassigned | 956 | 1122 | 251 | 5531 | 5819 | 965.724 | 1112.276 | 0.76224 | NS |
| OTC_Unassigned | 94 | 106 | 12 | 3105 | 2494 | 92.947 | 107.053 | 0.916 | NS |
| OTC_microglia | 179 | 120 | 16 | 1979 | 1366 | 138.956 | 160.044 | 0.0010322 | Uniform |
| OC_Unassigned | 35 | 35 | 3 | 1408 | 1106 | 32.532 | 37.468 | 0.67632 | NS |
| OC_microglia | 74 | 31 | 6 | 1111 | 688 | 48.797 | 56.203 | 0.00041658 | Uniform |
| OC_oligodendrocytes | 150 | 215 | 31 | 537 | 815 | 169.629 | 195.371 | 0.14309 | NS |
| OTC_oligodendrocytes | 115 | 178 | 29 | 383 | 634 | 136.168 | 156.832 | 0.07723 | NS |
| OTC_astrocytes | 127 | 178 | 30 | 429 | 550 | 141.745 | 163.255 | 0.22916 | NS |
| CB_Neurins_In_Purkinje | 311 | 312 | 28 | 137 | 340 | 289.531 | 333.469 | 0.22353 | NS |
| OTC_endothelial | 54 | 66 | 8 | 477 | 413 | 55.768 | 64.232 | 0.81876 | NS |
| OTC_excitatory | 48 | 39 | 4 | 419 | 487 | 40.432 | 46.568 | 0.25114 | NS |
| CB_Neurons_Ex | 25 | 62 | 15 | 297 | 321 | 40.432 | 46.568 | 0.015726 | NS |

| | | | | | | | | | |
|---|---|---|---|---|---|---|---|---|---|
| CB_Macrophages | 18 | 34 | 3 | 366 | 275 | 24.166 | 27.834 | 0.21812 | NS |
| OTC_inhibitory | 21 | 23 | 0 | 241 | 338 | 20.448 | 23.552 | 0.90623 | NS |
| CB_Astrocytes | 47 | 67 | 8 | 199 | 236 | 52.98 | 61.02 | 0.42479 | NS |
| OC_astrocytes | 17 | 26 | 7 | 188 | 195 | 19.984 | 23.016 | 0.51577 | NS |
| OTC_pericytes | 11 | 21 | 0 | 158 | 155 | 14.872 | 17.128 | 0.32406 | NS |
| CB_Bergmann1 | 22 | 27 | 4 | 125 | 153 | 22.772 | 26.228 | 0.87558 | NS |
| CB_Fibroblasts | 8 | 23 | 2 | 144 | 128 | 14.407 | 16.593 | 0.090319 | NS |
| OC_endothelial | 4 | 6 | 0 | 80 | 76 | 4.647 | 5.353 | 0.77013 | NS |
| OC_neuron | 23 | 21 | 8 | 56 | 53 | 20.448 | 23.552 | 0.58642 | NS |
| OC_pericytes | 5 | 5 | 0 | 70 | 51 | 4.647 | 5.353 | 0.87462 | NS |
| CB_Neurons? | 22 | 16 | 3 | 38 | 39 | 17.66 | 20.34 | 0.31895 | NS |
| CB_Endothelial | 4 | 3 | 0 | 48 | 61 | 3.253 | 3.747 | 0.68955 | NS |
| OTC_neuron | 26 | 30 | 4 | 16 | 35 | 26.025 | 29.975 | 0.99618 | NS |
| CB_Oligodendrocytes | 8 | 8 | 1 | 36 | 45 | 7.436 | 8.564 | 0.84179 | NS |
| CB_Neurons_In | 18 | 13 | 3 | 23 | 31 | 14.407 | 16.593 | 0.36093 | NS |
| CB_Mural | 3 | 6 | 0 | 30 | 32 | 4.183 | 4.817 | 0.5692 | NS |
| OTC_OPCs | 0 | 0 | 0 | 22 | 17 | 0 | 0 | 1 | NS |
| CB_Lymphocytes | 0 | 0 | 0 | 0 | 2 | 0 | 0 | 1 | NS |

CTR018, haplotyped: 40699, With Celltype: 40699

| Region_Celltype | X1 | X2 | Both | Unknown | LowCoverage | Expected1 | Expected2 | P_value | Direction |
|---|---|---|---|---|---|---|---|---|---|
| FC_Unassigned | 423 | 811 | 264 | 3427 | 5156 | 455.864 | 778.136 | 0.16713 | NS |
| FC_Astrocytes1 | 1563 | 3250 | 456 | 460 | 1260 | 1778.016 | 3034.984 | 4.1516e-06 | Outlier |
| TC_Unassigned | 133 | 196 | 70 | 1612 | 1678 | 121.539 | 207.461 | 0.35893 | NS |
| FC_Microglia | 248 | 311 | 110 | 1418 | 1574 | 206.505 | 352.495 | 0.01152 | NS |
| TC_Microglia | 418 | 123 | 53 | 1326 | 1244 | 199.856 | 341.144 | 6.0911e-41 | Uniform |
| OC_Unassigned | 130 | 172 | 34 | 1322 | 1329 | 111.565 | 190.435 | 0.12572 | NS |
| OC_Astrocytes1 | 311 | 569 | 35 | 694 | 1097 | 325.089 | 554.911 | 0.48451 | NS |
| TC_Astrocytes1 | 405 | 935 | 85 | 241 | 517 | 495.022 | 844.978 | 0.00023142 | Outlier |
| OC_Microglia | 88 | 46 | 11 | 451 | 382 | 49.502 | 84.498 | 2.5404e-06 | Uniform |
| FC_Astrocytes2 | 127 | 274 | 51 | 160 | 274 | 148.137 | 252.863 | 0.1159 | NS |
| TC_Astrocytes2 | 72 | 144 | 11 | 116 | 158 | 79.795 | 136.205 | 0.43214 | NS |
| TC_CAMs | 61 | 13 | 12 | 216 | 198 | 27.337 | 46.663 | 1.6902e-08 | Uniform |
| OC_CAMs | 49 | 10 | 5 | 186 | 161 | 21.796 | 37.204 | 3.1891e-07 | Uniform |
| OC_Astrocytes2 | 14 | 55 | 1 | 96 | 84 | 25.49 | 43.51 | 0.030458 | NS |
| FC_Endothelial | 12 | 32 | 2 | 62 | 90 | 16.254 | 27.746 | 0.33136 | NS |
| OC_Oligodendrocytes | 16 | 14 | 2 | 58 | 84 | 11.083 | 18.917 | 0.20205 | NS |
| FC_Pericytes | 9 | 17 | 1 | 51 | 59 | 9.605 | 16.395 | 0.86108 | NS |
| OC_Neurons | 22 | 45 | 5 | 26 | 37 | 24.751 | 42.249 | 0.61804 | NS |
| FC_Oligodendrocytes | 14 | 21 | 4 | 28 | 58 | 12.93 | 22.07 | 0.7926 | NS |
| FC_CAMs | 9 | 5 | 1 | 45 | 48 | 5.172 | 8.828 | 0.1479 | NS |
| FC_Lymphocytes | 6 | 4 | 3 | 47 | 46 | 3.694 | 6.306 | 0.30223 | NS |
| TC_Oligodendrocytes | 8 | 16 | 1 | 28 | 44 | 8.866 | 15.134 | 0.79344 | NS |

| | | | | | | | | | |
|---|---|---|---|---|---|---|---|---|---|
| TC_Endothelial | 10 | 7 | 1 | 33 | 36 | 6.28 | 10.72 | 0.20158 | NS |
| FC_Fibroblasts | 9 | 9 | 1 | 26 | 40 | 6.65 | 11.35 | 0.42938 | NS |
| TC_Pericytes | 7 | 7 | 2 | 24 | 34 | 5.172 | 8.828 | 0.48584 | NS |
| OC_Endothelial | 6 | 7 | 2 | 24 | 21 | 4.802 | 8.198 | 0.63366 | NS |
| TC_Neurons | 3 | 19 | 3 | 18 | 13 | 8.127 | 13.873 | 0.07536 | NS |
| OC_Pericytes | 0 | 9 | 1 | 18 | 19 | 3.325 | 5.675 | 0.043444 | NS |
| OC_Fibroblasts | 2 | 6 | 0 | 17 | 22 | 2.955 | 5.045 | 0.60547 | NS |
| FC_Neurons | 3 | 11 | 3 | 9 | 17 | 5.172 | 8.828 | 0.36661 | NS |
| TC_Lymphocytes | 4 | 1 | 0 | 12 | 18 | 1.847 | 3.153 | 0.1671 | NS |
| TC_Fibroblasts | 1 | 4 | 0 | 9 | 16 | 1.847 | 3.153 | 0.55278 | NS |
| OC_Lymphocytes | 2 | 0 | 0 | 12 | 7 | 0.739 | 1.261 | 0.17473 | NS |
| FC_SMCs | 0 | 3 | 0 | 8 | 9 | 1.108 | 1.892 | 0.24365 | NS |
| TC_SMCs | 1 | 1 | 0 | 6 | 10 | 0.739 | 1.261 | 0.79223 | NS |
| OC_SMCs | 0 | 0 | 0 | 2 | 2 | 0 | 0 | 1 | NS |
| FC_OPCs | 0 | 0 | 1 | 1 | 1 | 0 | 0 | 1 | NS |
| TC_OPCs | 1 | 0 | 0 | 1 | 0 | 0.369 | 0.631 | 0.33723 | NS |
| OC_OPCs | 0 | 0 | 0 | 1 | 0 | 0 | 0 | 1 | NS |

CTR081, haplotyped: 25149, With Celltype: 25149

| Region_Celltype | X1 | X2 | Both | Unknown | LowCoverage | Expected1 | Expected2 | P_value | Direction |
|---|---|---|---|---|---|---|---|---|---|
| FC_Astrocytes1 | 1066 | 1631 | 186 | 550 | 2158 | 914.145 | 1782.855 | 1.7901e-05 | Uniform |
| TC_Astrocytes1 | 517 | 1201 | 107 | 278 | 1287 | 582.314 | 1135.686 | 0.016906 | NS |
| FC_Unassigned | 128 | 221 | 46 | 594 | 1843 | 118.293 | 230.707 | 0.44197 | NS |
| TC_Microglia | 46 | 336 | 16 | 618 | 1524 | 129.478 | 252.522 | 6.9683e-13 | Outlier |
| OC_Astrocytes2 | 347 | 546 | 45 | 231 | 1242 | 302.681 | 590.319 | 0.029268 | NS |
| TC_Unassigned | 109 | 293 | 76 | 385 | 1321 | 136.257 | 265.743 | 0.036813 | NS |
| OC_Unassigned | 76 | 158 | 27 | 322 | 1204 | 79.314 | 154.686 | 0.74494 | NS |
| TC_Astrocytes2 | 132 | 279 | 17 | 106 | 499 | 139.308 | 271.692 | 0.58778 | NS |
| FC_Microglia | 15 | 80 | 3 | 192 | 376 | 32.2 | 62.8 | 0.003879 | Outlier |
| OC_Microglia | 27 | 75 | 5 | 114 | 335 | 34.573 | 67.427 | 0.24809 | NS |
| FC_Astrocytes2 | 78 | 113 | 13 | 81 | 255 | 64.739 | 126.261 | 0.16077 | NS |
| OC_Astrocytes1 | 40 | 84 | 3 | 29 | 153 | 42.03 | 81.97 | 0.78414 | NS |
| TC_Oligodendrocytes | 12 | 29 | 6 | 11 | 67 | 13.897 | 27.103 | 0.65225 | NS |
| OC_Endothelial | 5 | 13 | 3 | 31 | 62 | 6.101 | 11.899 | 0.69109 | NS |
| FC_Endothelial | 11 | 7 | 3 | 16 | 65 | 6.101 | 11.899 | 0.10205 | NS |
| FC_Pericytes | 7 | 9 | 0 | 22 | 54 | 5.423 | 10.577 | 0.56735 | NS |
| FC_CAMs | 1 | 8 | 0 | 25 | 51 | 3.051 | 5.949 | 0.24712 | NS |
| OC_Pericytes | 3 | 16 | 1 | 17 | 42 | 6.44 | 12.56 | 0.19654 | NS |
| OC_Oligodendrocytes | 6 | 21 | 1 | 10 | 38 | 9.152 | 17.848 | 0.33979 | NS |
| TC_CAMs | 1 | 8 | 0 | 9 | 44 | 3.051 | 5.949 | 0.24712 | NS |
| FC_Fibroblasts | 8 | 9 | 0 | 7 | 31 | 5.762 | 11.238 | 0.43428 | NS |
| FC_Oligodendrocytes | 4 | 10 | 0 | 13 | 26 | 4.745 | 9.255 | 0.7612 | NS |
| OC_Fibroblasts | 3 | 9 | 2 | 2 | 34 | 4.067 | 7.933 | 0.63264 | NS |

| | | | | | | | | | |
|---|---|---|---|---|---|---|---|---|---|
| FC_Neurons | 17 | 19 | 4 | 2 | 8 | 12.202 | 23.798 | 0.24949 | NS |
| TC_Pericytes | 1 | 6 | 0 | 11 | 30 | 2.373 | 4.627 | 0.39096 | NS |
| TC_Endothelial | 0 | 6 | 1 | 15 | 25 | 2.034 | 3.966 | 0.11762 | NS |
| FC_Lymphocytes | 1 | 1 | 0 | 15 | 29 | 0.678 | 1.322 | 0.74415 | NS |
| TC_Lymphocytes | 2 | 3 | 0 | 6 | 29 | 1.695 | 3.305 | 0.84148 | NS |
| OC_Lymphocytes | 2 | 3 | 0 | 6 | 28 | 1.695 | 3.305 | 0.84148 | NS |
| OC_CAMs | 1 | 6 | 0 | 12 | 18 | 2.373 | 4.627 | 0.39096 | NS |
| TC_Fibroblasts | 1 | 5 | 0 | 3 | 18 | 2.034 | 3.966 | 0.49235 | NS |
| TC_Neurons | 4 | 8 | 5 | 0 | 6 | 4.067 | 7.933 | 0.97677 | NS |
| OC_SMCs | 0 | 1 | 0 | 3 | 10 | 0.339 | 0.661 | 0.52293 | NS |
| FC_Mesenchymal | 0 | 1 | 1 | 1 | 7 | 0.339 | 0.661 | 0.52293 | NS |
| TC_OPCs | 2 | 1 | 0 | 2 | 5 | 1.017 | 1.983 | 0.42212 | NS |
| FC_SMCs | 0 | 0 | 1 | 1 | 7 | 0 | 0 | 1 | NS |
| OC_Neurons | 2 | 2 | 1 | 1 | 3 | 1.356 | 2.644 | 0.64441 | NS |
| OC_OPCs | 1 | 0 | 0 | 0 | 2 | 0.339 | 0.661 | 0.32037 | NS |
| FC_OPCs | 0 | 1 | 0 | 2 | 0 | 0.339 | 0.661 | 0.52293 | NS |
| TC_SMCs | 0 | 0 | 0 | 1 | 1 | 0 | 0 | 1 | NS |

CTR148, haplotyped: 20916, With Celltype: 20916

| Region_Celltype | X1 | X2 | Both | Unknown | LowCoverage | Expected1 | Expected2 | P_value | Direction |
|---|---|---|---|---|---|---|---|---|---|
| FC_Unassigned | 217 | 244 | 87 | 1214 | 2288 | 196.74 | 264.26 | 0.17975 | NS |
| FC_Astrocytes1 | 1011 | 1488 | 176 | 274 | 1048 | 1066.494 | 1432.506 | 0.11122 | NS |
| OC_Unassigned | 95 | 143 | 24 | 820 | 1316 | 101.571 | 136.429 | 0.54075 | NS |
| FC_Microglia | 225 | 75 | 31 | 914 | 1019 | 128.03 | 171.97 | 8.6806e-16 | Uniform |
| TC_Unassigned | 74 | 90 | 13 | 548 | 1082 | 69.99 | 94.01 | 0.65548 | NS |
| OC_Astrocytes2 | 180 | 216 | 21 | 341 | 710 | 169 | 227 | 0.43111 | NS |
| OC_Microglia | 49 | 119 | 10 | 580 | 530 | 71.697 | 96.303 | 0.0098552 | Outlier |
| TC_Astrocytes2 | 67 | 104 | 21 | 237 | 643 | 72.977 | 98.023 | 0.51096 | NS |
| TC_Microglia | 21 | 24 | 0 | 285 | 257 | 19.205 | 25.795 | 0.70344 | NS |
| FC_Astrocytes2 | 65 | 106 | 9 | 44 | 118 | 72.977 | 98.023 | 0.37925 | NS |
| OC_Astrocytes1 | 26 | 36 | 5 | 30 | 99 | 26.46 | 35.54 | 0.93342 | NS |
| OC_Oligodendrocytes | 20 | 23 | 2 | 53 | 94 | 18.351 | 24.649 | 0.72055 | NS |
| TC_Astrocytes1 | 10 | 36 | 2 | 32 | 83 | 19.631 | 26.369 | 0.03164 | NS |
| OC_Pericytes | 7 | 27 | 0 | 52 | 66 | 14.51 | 19.49 | 0.050184 | NS |
| FC_Neurons | 19 | 34 | 16 | 13 | 30 | 22.619 | 30.381 | 0.47168 | NS |
| OC_Endothelial | 3 | 9 | 1 | 44 | 49 | 5.121 | 6.879 | 0.36014 | NS |
| FC_CAMs | 5 | 5 | 1 | 39 | 42 | 4.268 | 5.732 | 0.74262 | NS |
| TC_Oligodendrocytes | 4 | 9 | 1 | 22 | 44 | 5.548 | 7.452 | 0.52884 | NS |
| FC_Pericytes | 1 | 15 | 1 | 28 | 35 | 6.828 | 9.172 | 0.01654 | NS |
| FC_Endothelial | 1 | 4 | 0 | 25 | 40 | 2.134 | 2.866 | 0.43955 | NS |
| OC_CAMs | 6 | 5 | 0 | 23 | 34 | 4.694 | 6.306 | 0.57759 | NS |
| FC_Oligodendrocytes | 4 | 13 | 2 | 14 | 22 | 7.255 | 9.745 | 0.23552 | NS |
| OC_Neurons | 5 | 7 | 1 | 7 | 20 | 5.121 | 6.879 | 0.96004 | NS |

| | | | | | | | | | |
|---|---|---|---|---|---|---|---|---|---|
| OC_Lymphocytes | 1 | 2 | 0 | 20 | 14 | 1.28 | 1.72 | 0.81362 | NS |
| FC_Fibroblasts | 3 | 7 | 1 | 11 | 14 | 4.268 | 5.732 | 0.55563 | NS |
| OC_Fibroblasts | 1 | 4 | 0 | 8 | 19 | 2.134 | 2.866 | 0.43955 | NS |
| FC_Lymphocytes | 1 | 0 | 0 | 10 | 19 | 0.427 | 0.573 | 0.37004 | NS |
| TC_Neurons | 3 | 9 | 1 | 2 | 6 | 5.121 | 6.879 | 0.36014 | NS |
| TC_Pericytes | 2 | 2 | 0 | 6 | 8 | 1.707 | 2.293 | 0.83547 | NS |
| TC_Fibroblasts | 2 | 0 | 0 | 2 | 11 | 0.854 | 1.146 | 0.2049 | NS |
| OC_SMCs | 1 | 1 | 0 | 4 | 6 | 0.854 | 1.146 | 0.88324 | NS |
| FC_OPCs | 1 | 2 | 1 | 2 | 5 | 1.28 | 1.72 | 0.81362 | NS |
| TC_CAMs | 0 | 0 | 0 | 3 | 6 | 0 | 0 | 1 | NS |
| TC_Endothelial | 0 | 1 | 0 | 2 | 1 | 0.427 | 0.573 | 0.46138 | NS |
| FC_SMCs | 0 | 1 | 0 | 1 | 2 | 0.427 | 0.573 | 0.46138 | NS |
| TC_OPCs | 0 | 0 | 0 | 0 | 3 | 0 | 0 | 1 | NS |
| OC_Mesenchymal | 0 | 0 | 0 | 0 | 2 | 0 | 0 | 1 | NS |
| TC_Lymphocytes | 0 | 0 | 0 | 1 | 1 | 0 | 0 | 1 | NS |
| OC_OPCs | 0 | 0 | 0 | 0 | 1 | 0 | 0 | 1 | NS |

FTD014, haplotyped: 21143, With Celltype: 21143

| Region_Celltype | X1 | X2 | Both | Unknown | LowCoverage | Expected1 | Expected2 | P_value | Direction |
|---|---|---|---|---|---|---|---|---|---|
| TC_Unassigned | 176 | 317 | 107 | 1197 | 1745 | 193.657 | 299.343 | 0.2454 | NS |
| TC_Microglia | 176 | 563 | 53 | 1011 | 1484 | 290.29 | 448.71 | 1.5828e-10 | Outlier |
| TC_Astrocytes1 | 523 | 806 | 74 | 271 | 689 | 522.05 | 806.95 | 0.96991 | NS |
| FC_Astrocytes1 | 394 | 463 | 53 | 361 | 848 | 336.642 | 520.358 | 0.0050863 | Uniform |
| FC_Unassigned | 54 | 98 | 32 | 583 | 997 | 59.708 | 92.292 | 0.4987 | NS |
| TC_Astrocytes2 | 365 | 423 | 36 | 98 | 278 | 309.538 | 478.462 | 0.004749 | Uniform |
| OC_Unassigned | 32 | 40 | 9 | 481 | 617 | 28.283 | 43.717 | 0.53005 | NS |
| FC_Microglia | 20 | 95 | 12 | 430 | 553 | 45.174 | 69.826 | 0.00023006 | Outlier |
| FC_Astrocytes2 | 228 | 241 | 31 | 135 | 395 | 184.23 | 284.77 | 0.0039835 | Uniform |
| OC_Microglia | 29 | 48 | 4 | 362 | 371 | 30.247 | 46.753 | 0.83641 | NS |
| TC_Neurons | 68 | 107 | 7 | 112 | 314 | 68.742 | 106.258 | 0.93517 | NS |
| TC_Oligodendrocytes | 35 | 49 | 7 | 37 | 85 | 32.996 | 51.004 | 0.75281 | NS |
| OC_Astrocytes1 | 35 | 45 | 1 | 44 | 87 | 31.425 | 48.575 | 0.56627 | NS |
| FC_Oligodendrocytes | 21 | 33 | 3 | 44 | 110 | 21.212 | 32.788 | 0.96666 | NS |
| TC_CAMs | 12 | 26 | 3 | 60 | 89 | 14.927 | 23.073 | 0.48271 | NS |
| OC_Oligodendrocytes | 24 | 20 | 4 | 40 | 97 | 17.284 | 26.716 | 0.15139 | NS |
| OC_Neurons | 11 | 24 | 0 | 42 | 104 | 13.748 | 21.252 | 0.49199 | NS |
| OC_Astrocytes2 | 23 | 34 | 6 | 36 | 80 | 22.39 | 34.61 | 0.90715 | NS |
| FC_Neurons | 14 | 24 | 5 | 34 | 87 | 14.927 | 23.073 | 0.82666 | NS |
| TC_Lymphocytes | 4 | 13 | 2 | 36 | 54 | 6.678 | 10.322 | 0.32244 | NS |
| TC_Fibroblasts | 9 | 17 | 0 | 20 | 26 | 10.213 | 15.787 | 0.72742 | NS |
| OC_CAMs | 3 | 2 | 0 | 29 | 31 | 1.964 | 3.036 | 0.51234 | NS |
| OC_Lymphocytes | 1 | 1 | 0 | 17 | 22 | 0.786 | 1.214 | 0.82929 | NS |
| FC_CAMs | 2 | 5 | 0 | 18 | 13 | 2.75 | 4.25 | 0.67215 | NS |

| | | | | | | | | | |
|---|---|---|---|---|---|---|---|---|---|
| TC_Pericytes | 2 | 2 | 2 | 10 | 16 | 1.571 | 2.429 | 0.76042 | NS |
| OC_Endothelial | 1 | 2 | 0 | 11 | 18 | 1.178 | 1.822 | 0.87959 | NS |
| OC_Fibroblasts | 2 | 4 | 0 | 14 | 12 | 2.357 | 3.643 | 0.83036 | NS |
| FC_Pericytes | 2 | 3 | 0 | 8 | 14 | 1.964 | 3.036 | 0.98147 | NS |
| OC_Pericytes | 2 | 6 | 0 | 7 | 10 | 3.143 | 4.857 | 0.5408 | NS |
| TC_SMCs | 1 | 2 | 2 | 6 | 13 | 1.178 | 1.822 | 0.87959 | NS |
| TC_OPCs | 9 | 6 | 0 | 1 | 4 | 5.892 | 9.108 | 0.25645 | NS |
| FC_Lymphocytes | 0 | 1 | 0 | 7 | 8 | 0.393 | 0.607 | 0.48445 | NS |
| TC_Endothelial | 1 | 2 | 0 | 5 | 7 | 1.178 | 1.822 | 0.87959 | NS |
| FC_Endothelial | 0 | 3 | 0 | 4 | 4 | 1.178 | 1.822 | 0.2259 | NS |
| FC_OPCs | 3 | 1 | 0 | 2 | 5 | 1.571 | 2.429 | 0.30738 | NS |
| OC_Mesenchymal | 0 | 3 | 0 | 7 | 1 | 1.178 | 1.822 | 0.2259 | NS |
| FC_Fibroblasts | 2 | 3 | 0 | 1 | 1 | 1.964 | 3.036 | 0.98147 | NS |
| OC_OPCs | 0 | 0 | 0 | 1 | 2 | 0 | 0 | 1 | NS |
| FC_SMCs | 1 | 0 | 0 | 0 | 0 | 0.393 | 0.607 | 0.35043 | NS |

FTD024, haplotyped: 27188, With Celltype: 27188

| Region_Celltype | X1 | X2 | Both | Unknown | LowCoverage | Expected1 | Expected2 | P_value | Direction |
|---|---|---|---|---|---|---|---|---|---|
| TC_Astrocytes1 | 934 | 984 | 124 | 493 | 1710 | 916.191 | 1001.809 | 0.56499 | NS |
| FC_Astrocytes1 | 840 | 997 | 138 | 480 | 1702 | 877.499 | 959.501 | 0.215 | NS |
| FC_Unassigned | 215 | 242 | 75 | 719 | 1799 | 218.3 | 238.7 | 0.82697 | NS |
| TC_Unassigned | 197 | 193 | 96 | 617 | 1455 | 186.295 | 203.705 | 0.44326 | NS |
| TC_Microglia | 211 | 87 | 19 | 522 | 967 | 142.349 | 155.651 | 1.0421e-08 | Uniform |
| OC_Unassigned | 101 | 149 | 56 | 508 | 975 | 119.42 | 130.58 | 0.097077 | NS |
| OC_Microglia | 217 | 103 | 18 | 509 | 882 | 152.858 | 167.142 | 2.8427e-07 | Uniform |
| OC_Astrocytes1 | 199 | 411 | 32 | 221 | 667 | 291.385 | 318.615 | 6.8646e-08 | Outlier |
| FC_Microglia | 145 | 116 | 26 | 415 | 656 | 124.675 | 136.325 | 0.075039 | NS |
| FC_Astrocytes2 | 250 | 263 | 27 | 165 | 531 | 245.05 | 267.95 | 0.75712 | NS |
| OC_Astrocytes2 | 177 | 288 | 21 | 142 | 447 | 222.121 | 242.879 | 0.002796 | Outlier |
| TC_Astrocytes2 | 207 | 191 | 20 | 125 | 409 | 190.117 | 207.883 | 0.23137 | NS |
| OC_Endothelial | 28 | 30 | 2 | 95 | 152 | 27.705 | 30.295 | 0.95635 | NS |
| TC_Fibroblasts | 18 | 21 | 1 | 29 | 74 | 18.63 | 20.37 | 0.88643 | NS |
| TC_CAMs | 7 | 18 | 1 | 41 | 75 | 11.942 | 13.058 | 0.14966 | NS |
| OC_Oligodendrocytes | 13 | 28 | 9 | 11 | 45 | 19.585 | 21.415 | 0.13728 | NS |
| FC_Fibroblasts | 13 | 12 | 0 | 11 | 65 | 11.942 | 13.058 | 0.76475 | NS |
| OC_Pericytes | 14 | 6 | 0 | 16 | 52 | 9.554 | 10.446 | 0.15306 | NS |
| TC_Oligodendrocytes | 18 | 8 | 7 | 14 | 41 | 12.42 | 13.58 | 0.11629 | NS |
| FC_CAMs | 4 | 11 | 3 | 23 | 42 | 7.165 | 7.835 | 0.23189 | NS |
| TC_Lymphocytes | 4 | 3 | 1 | 23 | 43 | 3.344 | 3.656 | 0.72544 | NS |
| FC_Oligodendrocytes | 2 | 10 | 3 | 18 | 32 | 5.732 | 6.268 | 0.10305 | NS |
| OC_CAMs | 2 | 13 | 0 | 12 | 29 | 7.165 | 7.835 | 0.040628 | NS |
| OC_Fibroblasts | 11 | 4 | 0 | 11 | 27 | 7.165 | 7.835 | 0.15199 | NS |
| TC_Endothelial | 4 | 1 | 0 | 15 | 33 | 2.388 | 2.612 | 0.28869 | NS |

| | | | | | | | | | |
|---|---|---|---|---|---|---|---|---|---|
| OC_Lymphocytes | 5 | 5 | 1 | 14 | 25 | 4.777 | 5.223 | 0.92047 | NS |
| TC_Pericytes | 2 | 4 | 1 | 11 | 31 | 2.866 | 3.134 | 0.61061 | NS |
| TC_Mesenchymal | 9 | 2 | 0 | 7 | 30 | 5.254 | 5.746 | 0.094535 | NS |
| FC_Pericytes | 1 | 4 | 1 | 10 | 21 | 2.388 | 2.612 | 0.35361 | NS |
| FC_Endothelial | 1 | 1 | 0 | 11 | 23 | 0.955 | 1.045 | 0.96439 | NS |
| FC_Neurons | 2 | 8 | 2 | 7 | 7 | 4.777 | 5.223 | 0.18958 | NS |
| FC_Lymphocytes | 0 | 1 | 1 | 7 | 16 | 0.478 | 0.522 | 0.42825 | NS |
| TC_Neurons | 7 | 2 | 2 | 1 | 7 | 4.299 | 4.701 | 0.18787 | NS |
| FC_SMCs | 3 | 1 | 0 | 3 | 7 | 1.911 | 2.089 | 0.42894 | NS |
| TC_OPCs | 1 | 1 | 0 | 2 | 6 | 0.955 | 1.045 | 0.96439 | NS |
| TC_SMCs | 0 | 0 | 0 | 2 | 7 | 0 | 0 | 1 | NS |
| OC_Neurons | 1 | 4 | 0 | 0 | 4 | 2.388 | 2.612 | 0.35361 | NS |
| FC_OPCs | 0 | 1 | 0 | 0 | 4 | 0.478 | 0.522 | 0.42825 | NS |
| OC_SMCs | 0 | 1 | 0 | 0 | 3 | 0.478 | 0.522 | 0.42825 | NS |
| OC_OPCs | 0 | 0 | 0 | 2 | 0 | 0 | 0 | 1 | NS |
| OC_Mesenchymal | 0 | 0 | 1 | 0 | 0 | 0 | 0 | 1 | NS |

FTD038, haplotyped: 30507, With Celltype: 30507

| Region_Celltype | X1 | X2 | Both | Unknown | LowCoverage | Expected1 | Expected2 | P_value | Direction |
|---|---|---|---|---|---|---|---|---|---|
| OC_Unassigned | 132 | 518 | 137 | 1095 | 1872 | 190.649 | 459.351 | 0.00016609 | Outlier |
| TC_Unassigned | 133 | 267 | 106 | 1290 | 1822 | 117.323 | 282.677 | 0.23193 | NS |
| FC_Unassigned | 156 | 232 | 54 | 1162 | 2003 | 113.803 | 274.197 | 0.001469 | Uniform |
| OC_Astrocytes1 | 222 | 1008 | 94 | 641 | 1480 | 360.767 | 869.233 | 4.6937e-11 | Outlier |
| TC_Microglia | 330 | 380 | 61 | 898 | 1437 | 208.248 | 501.752 | 2.7434e-11 | Uniform |
| OC_Microglia | 220 | 437 | 45 | 642 | 1131 | 192.703 | 464.297 | 0.10471 | NS |
| TC_Astrocytes1 | 185 | 623 | 71 | 210 | 466 | 236.992 | 571.008 | 0.0032355 | Outlier |
| FC_Astrocytes1 | 133 | 371 | 32 | 265 | 750 | 147.827 | 356.173 | 0.29756 | NS |
| FC_Microglia | 150 | 77 | 18 | 340 | 797 | 66.581 | 160.419 | 4.5636e-15 | Uniform |
| OC_Astrocytes2 | 102 | 397 | 25 | 227 | 529 | 146.36 | 352.64 | 0.001163 | Outlier |
| TC_Fibroblasts | 161 | 275 | 35 | 207 | 416 | 127.882 | 308.118 | 0.017182 | NS |
| FC_Astrocytes2 | 53 | 153 | 17 | 164 | 355 | 60.421 | 145.579 | 0.41304 | NS |
| TC_Astrocytes2 | 51 | 227 | 17 | 102 | 191 | 81.539 | 196.461 | 0.0023689 | Outlier |
| TC_CAMs | 23 | 37 | 3 | 111 | 146 | 17.598 | 42.402 | 0.29733 | NS |
| OC_Endothelial | 9 | 71 | 7 | 89 | 136 | 23.465 | 56.535 | 0.0044629 | Outlier |
| OC_Fibroblasts | 35 | 53 | 6 | 65 | 143 | 25.811 | 62.189 | 0.14524 | NS |
| TC_Mesenchymal | 15 | 28 | 3 | 44 | 80 | 12.612 | 30.388 | 0.5813 | NS |
| FC_Fibroblasts | 14 | 20 | 2 | 21 | 90 | 9.972 | 24.028 | 0.30664 | NS |
| OC_CAMs | 21 | 18 | 3 | 35 | 66 | 11.439 | 27.561 | 0.02806 | NS |
| OC_Lymphocytes | 9 | 10 | 0 | 39 | 68 | 5.573 | 13.427 | 0.25288 | NS |
| OC_Pericytes | 8 | 19 | 1 | 19 | 51 | 7.919 | 19.081 | 0.98078 | NS |
| TC_Endothelial | 4 | 5 | 2 | 23 | 49 | 2.64 | 6.36 | 0.50638 | NS |
| TC_Oligodendrocytes | 1 | 19 | 3 | 16 | 32 | 5.866 | 14.134 | 0.041307 | NS |
| TC_Lymphocytes | 8 | 9 | 0 | 14 | 30 | 4.986 | 12.014 | 0.28742 | NS |

| | | | | | | | | | |
|---|---|---|---|---|---|---|---|---|---|
| FC_CAMs | 5 | 5 | 1 | 11 | 37 | 2.933 | 7.067 | 0.34478 | NS |
| FC_Pericytes | 0 | 5 | 1 | 9 | 41 | 1.467 | 3.533 | 0.18988 | NS |
| TC_Pericytes | 5 | 13 | 1 | 12 | 16 | 5.28 | 12.72 | 0.91785 | NS |
| FC_Oligodendrocytes | 4 | 11 | 3 | 4 | 24 | 4.4 | 10.6 | 0.87092 | NS |
| FC_Mesenchymal | 7 | 4 | 1 | 10 | 14 | 3.226 | 7.774 | 0.10673 | NS |
| OC_Oligodendrocytes | 3 | 7 | 1 | 8 | 15 | 2.933 | 7.067 | 0.97386 | NS |
| FC_Endothelial | 0 | 3 | 0 | 9 | 17 | 0.88 | 2.12 | 0.30989 | NS |
| OC_SMCs | 4 | 3 | 1 | 5 | 16 | 2.053 | 4.947 | 0.29359 | NS |
| OC_Mesenchymal | 4 | 2 | 1 | 7 | 12 | 1.76 | 4.24 | 0.19553 | NS |
| TC_SMCs | 0 | 0 | 2 | 11 | 12 | 0 | 0 | 1 | NS |
| OC_Neurons | 1 | 11 | 2 | 2 | 9 | 3.52 | 8.48 | 0.18833 | NS |
| FC_Lymphocytes | 2 | 1 | 1 | 8 | 10 | 0.88 | 2.12 | 0.36005 | NS |
| TC_Neurons | 2 | 6 | 1 | 3 | 7 | 2.346 | 5.654 | 0.84561 | NS |
| FC_Neurons | 1 | 2 | 0 | 2 | 7 | 0.88 | 2.12 | 0.91583 | NS |
| OC_OPCs | 0 | 3 | 1 | 0 | 2 | 0.88 | 2.12 | 0.30989 | NS |
| FC_SMCs | 0 | 0 | 0 | 1 | 2 | 0 | 0 | 1 | NS |
| TC_OPCs | 0 | 2 | 0 | 0 | 1 | 0.587 | 1.413 | 0.40704 | NS |

FTD073, haplotyped: 35357, With Celltype: 35357

| Region_Celltype | X1 | X2 | Both | Unknown | LowCoverage | Expected1 | Expected2 | P_value | Direction |
|---|---|---|---|---|---|---|---|---|---|
| FC_Unassigned | 117 | 762 | 147 | 1054 | 3016 | 132.376 | 746.624 | 0.29322 | NS |
| FC_Astrocytes1 | 376 | 1896 | 120 | 411 | 1672 | 342.16 | 1929.84 | 0.16876 | NS |
| TC_Unassigned | 71 | 559 | 102 | 1019 | 2538 | 94.877 | 535.123 | 0.046649 | NS |
| TC_Microglia | 37 | 664 | 50 | 875 | 1771 | 105.57 | 595.43 | 1.3693e-09 | Outlier |
| FC_Astrocytes2 | 200 | 949 | 71 | 283 | 1009 | 173.038 | 975.962 | 0.12719 | NS |
| OC_Microglia | 51 | 398 | 29 | 653 | 1295 | 67.619 | 381.381 | 0.10145 | NS |
| OC_Astrocytes1 | 142 | 935 | 26 | 331 | 968 | 162.195 | 914.805 | 0.2115 | NS |
| TC_Astrocytes2 | 212 | 856 | 34 | 281 | 878 | 160.839 | 907.161 | 0.0035421 | Uniform |
| TC_Astrocytes1 | 188 | 965 | 73 | 117 | 537 | 173.64 | 979.36 | 0.41088 | NS |
| OC_Unassigned | 46 | 304 | 42 | 438 | 978 | 52.709 | 297.291 | 0.46622 | NS |
| FC_Microglia | 9 | 160 | 15 | 375 | 698 | 25.451 | 143.549 | 0.0031005 | Outlier |
| TC_Fibroblasts | 54 | 138 | 17 | 173 | 505 | 28.915 | 163.085 | 0.0018636 | Uniform |
| OC_Endothelial | 11 | 60 | 2 | 114 | 236 | 10.692 | 60.308 | 0.94282 | NS |
| TC_CAMs | 3 | 17 | 3 | 107 | 203 | 3.012 | 16.988 | 0.99577 | NS |
| FC_Fibroblasts | 11 | 24 | 4 | 62 | 135 | 5.271 | 29.729 | 0.10499 | NS |
| OC_Astrocytes2 | 11 | 82 | 0 | 36 | 101 | 14.006 | 78.994 | 0.51824 | NS |
| OC_Oligodendrocytes | 11 | 57 | 2 | 29 | 92 | 10.241 | 57.759 | 0.85766 | NS |
| OC_Fibroblasts | 9 | 18 | 1 | 47 | 93 | 4.066 | 22.934 | 0.11695 | NS |
| TC_Endothelial | 4 | 36 | 2 | 27 | 67 | 6.024 | 33.976 | 0.49428 | NS |
| FC_Oligodendrocytes | 4 | 7 | 2 | 31 | 69 | 1.657 | 9.343 | 0.25297 | NS |
| TC_Oligodendrocytes | 3 | 16 | 3 | 20 | 39 | 2.861 | 16.139 | 0.95035 | NS |
| FC_CAMs | 2 | 7 | 1 | 25 | 38 | 1.355 | 7.645 | 0.69643 | NS |
| FC_Pericytes | 2 | 8 | 1 | 21 | 36 | 1.506 | 8.494 | 0.77141 | NS |

| | | | | | | | | | |
|---|---|---|---|---|---|---|---|---|---|
| OC_CAMs | 3 | 6 | 0 | 22 | 36 | 1.355 | 7.645 | 0.3654 | NS |
| TC_Lymphocytes | 0 | 2 | 2 | 18 | 40 | 0.301 | 1.699 | 0.56819 | NS |
| TC_SMCs | 0 | 9 | 4 | 10 | 37 | 1.355 | 7.645 | 0.22602 | NS |
| OC_Pericytes | 1 | 8 | 0 | 10 | 37 | 1.355 | 7.645 | 0.80384 | NS |
| TC_Neurons | 6 | 26 | 3 | 6 | 14 | 4.819 | 27.181 | 0.69371 | NS |
| TC_Mesenchymal | 4 | 6 | 3 | 6 | 24 | 1.506 | 8.494 | 0.21183 | NS |
| TC_Pericytes | 2 | 9 | 0 | 7 | 25 | 1.657 | 9.343 | 0.84408 | NS |
| FC_Endothelial | 1 | 2 | 1 | 11 | 27 | 0.452 | 2.548 | 0.60127 | NS |
| FC_SMCs | 1 | 3 | 0 | 10 | 16 | 0.602 | 3.398 | 0.72541 | NS |
| OC_Lymphocytes | 0 | 2 | 0 | 13 | 14 | 0.301 | 1.699 | 0.56819 | NS |
| FC_Neurons | 0 | 11 | 1 | 4 | 13 | 1.657 | 9.343 | 0.18075 | NS |
| FC_Lymphocytes | 1 | 1 | 0 | 4 | 23 | 0.301 | 1.699 | 0.45578 | NS |
| FC_Mesenchymal | 1 | 2 | 0 | 2 | 13 | 0.452 | 2.548 | 0.60127 | NS |
| OC_Neurons | 3 | 3 | 1 | 2 | 8 | 0.904 | 5.096 | 0.19643 | NS |
| OC_SMCs | 0 | 1 | 0 | 6 | 8 | 0.151 | 0.849 | 0.68654 | NS |
| FC_OPCs | 0 | 1 | 1 | 0 | 4 | 0.151 | 0.849 | 0.68654 | NS |
| TC_OPCs | 1 | 0 | 1 | 1 | 3 | 0.151 | 0.849 | 0.22433 | NS |
| OC_OPCs | 0 | 3 | 0 | 0 | 1 | 0.452 | 2.548 | 0.48456 | NS |
| OC_Mesenchymal | 0 | 0 | 0 | 0 | 4 | 0 | 0 | 1 | NS |

FTD083, haplotyped: 15942, With Celltype: 15942

| Region_Celltype | X1 | X2 | Both | Unknown | LowCoverage | Expected1 | Expected2 | P_value | Direction |
|---|---|---|---|---|---|---|---|---|---|
| OC_Unassigned | 100 | 159 | 35 | 736 | 1105 | 96.615 | 162.385 | 0.75921 | NS |
| TC_Unassigned | 34 | 50 | 6 | 624 | 954 | 31.334 | 52.666 | 0.67314 | NS |
| TC_Astrocytes1 | 214 | 457 | 44 | 287 | 641 | 250.303 | 420.697 | 0.037228 | NS |
| FC_Unassigned | 71 | 107 | 27 | 445 | 738 | 66.399 | 111.601 | 0.61645 | NS |
| TC_Microglia | 57 | 91 | 15 | 539 | 542 | 55.208 | 92.792 | 0.83004 | NS |
| FC_Astrocytes1 | 172 | 274 | 38 | 209 | 484 | 166.371 | 279.629 | 0.6977 | NS |
| TC_Neurons | 134 | 304 | 38 | 191 | 427 | 163.387 | 274.613 | 0.036014 | NS |
| OC_Microglia | 55 | 31 | 6 | 453 | 459 | 32.081 | 53.919 | 0.00047324 | Uniform |
| OC_Neurons | 162 | 230 | 28 | 170 | 387 | 146.227 | 245.773 | 0.24881 | NS |
| OC_Astrocytes1 | 54 | 83 | 6 | 100 | 214 | 51.105 | 85.895 | 0.7191 | NS |
| FC_Astrocytes2 | 42 | 95 | 7 | 94 | 189 | 51.105 | 85.895 | 0.24551 | NS |
| FC_Microglia | 24 | 14 | 2 | 186 | 176 | 14.175 | 23.825 | 0.024196 | NS |
| FC_Oligodendrocytes | 32 | 32 | 6 | 51 | 153 | 23.874 | 40.126 | 0.14755 | NS |
| OC_Astrocytes2 | 36 | 64 | 5 | 45 | 95 | 37.303 | 62.697 | 0.84837 | NS |
| OC_Fibroblasts | 17 | 14 | 5 | 89 | 117 | 11.564 | 19.436 | 0.16604 | NS |
| OC_Endothelial | 5 | 23 | 1 | 58 | 79 | 10.445 | 17.555 | 0.10352 | NS |
| FC_Neurons | 22 | 40 | 3 | 24 | 55 | 23.128 | 38.872 | 0.83327 | NS |
| OC_Oligodendrocytes | 10 | 32 | 4 | 19 | 64 | 15.667 | 26.333 | 0.17948 | NS |
| TC_Oligodendrocytes | 12 | 23 | 4 | 29 | 54 | 13.056 | 21.944 | 0.79233 | NS |
| TC_Astrocytes2 | 16 | 34 | 1 | 16 | 53 | 18.651 | 31.349 | 0.57739 | NS |
| TC_Endothelial | 4 | 4 | 2 | 43 | 41 | 2.984 | 5.016 | 0.60863 | NS |

| | | | | | | | | | |
|---|---|---|---|---|---|---|---|---|---|
| OC_CAMs | 11 | 3 | 0 | 39 | 41 | 5.222 | 8.778 | 0.026983 | NS |
| FC_Fibroblasts | 6 | 11 | 1 | 32 | 34 | 6.341 | 10.659 | 0.90306 | NS |
| TC_CAMs | 1 | 4 | 1 | 41 | 29 | 1.865 | 3.135 | 0.54512 | NS |
| OC_Pericytes | 6 | 3 | 1 | 28 | 31 | 3.357 | 5.643 | 0.21248 | NS |
| TC_Fibroblasts | 6 | 4 | 2 | 21 | 33 | 3.73 | 6.27 | 0.30991 | NS |
| FC_CAMs | 3 | 4 | 1 | 23 | 30 | 2.611 | 4.389 | 0.83208 | NS |
| TC_Lymphocytes | 2 | 4 | 1 | 27 | 25 | 2.238 | 3.762 | 0.88562 | NS |
| OC_Lymphocytes | 0 | 2 | 0 | 17 | 22 | 0.746 | 1.254 | 0.33823 | NS |
| OC_Mesenchymal | 2 | 4 | 1 | 16 | 17 | 2.238 | 3.762 | 0.88562 | NS |
| FC_Lymphocytes | 0 | 5 | 0 | 17 | 11 | 1.865 | 3.135 | 0.12998 | NS |
| TC_OPCs | 3 | 7 | 3 | 2 | 15 | 3.73 | 6.27 | 0.72965 | NS |
| OC_OPCs | 6 | 3 | 1 | 13 | 7 | 3.357 | 5.643 | 0.21248 | NS |
| FC_Endothelial | 0 | 5 | 0 | 8 | 15 | 1.865 | 3.135 | 0.12998 | NS |
| FC_Pericytes | 3 | 1 | 0 | 5 | 12 | 1.492 | 2.508 | 0.28265 | NS |
| TC_Pericytes | 1 | 2 | 0 | 3 | 12 | 1.119 | 1.881 | 0.91898 | NS |
| OC_SMCs | 2 | 0 | 0 | 3 | 8 | 0.746 | 1.254 | 0.17654 | NS |
| TC_SMCs | 0 | 2 | 0 | 3 | 8 | 0.746 | 1.254 | 0.33823 | NS |
| FC_SMCs | 0 | 0 | 0 | 3 | 4 | 0 | 0 | 1 | NS |
| TC_Mesenchymal | 0 | 1 | 0 | 2 | 3 | 0.373 | 0.627 | 0.4983 | NS |
| FC_OPCs | 0 | 1 | 0 | 0 | 0 | 0.373 | 0.627 | 0.4983 | NS |

FTD6243, haplotyped: 4680, With Celltype: 4680

| Region_Celltype | X1 | X2 | Both | Unknown | LowCoverage | Expected1 | Expected2 | P_value | Direction |
|---|---|---|---|---|---|---|---|---|---|
| FC_Unassigned | 553 | 613 | 110 | 1213 | 1938 | 549.63 | 616.37 | 0.88882 | NS |
| FC_Microglia | 3 | 7 | 1 | 91 | 71 | 4.714 | 5.286 | 0.43111 | NS |
| FC_Oligodendrocytes | 1 | 4 | 0 | 11 | 16 | 2.357 | 2.643 | 0.36354 | NS |
| FC_Fibroblasts | 1 | 4 | 0 | 11 | 12 | 2.357 | 2.643 | 0.36354 | NS |
| FC_CAMs | 0 | 0 | 0 | 3 | 5 | 0 | 0 | 1 | NS |
| FC_Astrocytes2 | 1 | 0 | 0 | 1 | 5 | 0.471 | 0.529 | 0.39662 | NS |
| FC_Neurons | 0 | 0 | 0 | 0 | 3 | 0 | 0 | 1 | NS |
| FC_Mesenchymal | 1 | 0 | 0 | 0 | 0 | 0.471 | 0.529 | 0.39662 | NS |
| FC_Pericytes | 0 | 0 | 0 | 0 | 1 | 0 | 0 | 1 | NS |
