## Supplemental Table 5 for "Single-cell X-chromosome inactivation analysis links biased chimerism to differential gene expression and epigenetic erosion"

--- gene_id: ENSG00000161055 gene_name: SCGB3A1 gene_location: 5:180590105-180591499 ---

| Set | LogFC | LogCPM | P-val |
|---|---|---|---|
| 133C_ATII | -0.83272699574301 | 10.3401621525001 | 0.00472257970106345 |
| 1372C_Multiplet | 5.63972127063087 | 9.45685320866899 | 0.000920821016017894 |
| 157I_Ciliated | -1.82880579923839 | 10.5176113948531 | 1.0535534770344e-10 |
| 174I_Ciliated | -3.02312674989198 | 10.9951452589783 | 3.21250985177547e-08 |
| 184CO_ATII | 3.29345149331353 | 12.4705694856288 | 7.86602731975662e-05 |
| 184CO_Ciliated | -3.60303402379491 | 13.8875024416301 | 4.08018293049752e-15 |
| 209I_Ciliated | 1.73988304849014 | 10.070139950904 | 0.00991298245575162 |
| 209I_Goblet | -2.44346454986086 | 16.3966705640076 | 0.00164731183803709 |
| 217CO_Macrophage_Alveolar | -2.58908162341196 | 9.52369160569821 | 7.28162154531729e-08 |
| 225I_Ciliated | -1.45620218699937 | 10.8431128138871 | 3.06363492773248e-05 |
| 235CO_Ciliated | -3.61209336961982 | 12.0435785388004 | 2.83349904864869e-08 |
| 235CO_Multiplet | 5.65999242552416 | 15.3399877777939 | 1.40004210422575e-08 |

--- gene_id: ENSG00000164265 gene_name: SCGB3A2 gene_location: 5:147870682-147882191 ---

| Set | LogFC | LogCPM | P-val |
|---|---|---|---|
| 157I_Multiplet | -4.91661063348983 | 10.9654433692835 | 1.05734977126917e-14 |
| 184CO_ATII | 3.69093702196779 | 12.5147071098607 | 1.09020618141065e-05 |
| 184CO_Multiplet | -5.05543263189613 | 10.924408628852 | 2.31083592162409e-07 |
| 209I_Ciliated | 6.11106388087748 | 10.6669814890312 | 2.07725333422133e-24 |
| 209I_Goblet | 3.8150316931267 | 10.8268172982013 | 1.30594875167357e-05 |
| 209I_Multiplet | 4.29529580075911 | 9.45474451260695 | 0.00909991125066625 |
| 235CO_Ciliated | -4.63316473473335 | 9.17611202121073 | 8.78345585977805e-06 |
| 235CO_Multiplet | -5.22773227454284 | 11.5552418074616 | 1.2273893356545e-08 |

--- gene_id: ENSG00000275385 gene_name: CCL18 gene_location: 17:36064272-36072032 ---

| Set | LogFC | LogCPM | P-val |
|---|---|---|---|
| 052CO_Macrophage | 3.11839379549067 | 11.2177626962196 | 3.30601175820177e-06 |
| 133C_cDC2 | -2.92940736741037 | 9.85368959757177 | 1.13910807495421e-06 |
| 157I_Club | -2.93505794796778 | 8.62843881275706 | 5.42449809082384e-06 |
| 157I_cDC2 | -2.83350517500068 | 8.93117626791208 | 2.18969113555264e-06 |
| 207CO_Macrophage | -3.38652482637476 | 10.37394517992 | 5.49439575065809e-05 |
| 209I_Goblet | -2.62705376310179 | 8.56018166062272 | 0.00873875979729114 |
| 235CO_Macrophage | 3.6691111949865 | 10.4258687263044 | 6.39409689071296e-08 |

--- gene_id: ENSG00000233913 gene_name: RPL10P9 gene_location: 5:168616352-168616996 ---

| Set | LogFC | LogCPM | P-val |
|---|---|---|---|
| 235CO_Ciliated | 4.94052028715848 | 9.67945115995587 | 1.76643894104006e-06 |
| 235CO_Macrophage | 4.49422837074085 | 10.9907578817884 | 1.72782399451818e-30 |
| 235CO_Macrophage_Alveolar | -3.50455738959724 | 10.1099465249766 | 3.8142650719698e-52 |
| 235CO_Multiplet | -3.41209034551441 | 10.5990325865489 | 0.000509588543628087 |
| 235CO_cDC2 | -5.15177985982394 | 11.8222656005791 | 3.30449117410524e-21 |
| 235CO_cMonocyte | -4.28563924727613 | 11.3165178554323 | 1.88154717317586e-25 |
| 253C_Macrophage_Alveolar | 5.06863216164149 | 10.7760232085633 | 2.1988687456798e-26 |

--- gene_id: ENSG00000170323 gene_name: FABP4 gene_location: 8:81478419-81483236 ---

| Set | LogFC | LogCPM | P-val |
|---|---|---|---|
| 052CO_Macrophage | 3.33617044068723 | 9.48949563786928 | 0.00365264703955052 |
| 133C_cDC2 | -3.41909993397344 | 8.74750104152503 | 1.32326320209964e-06 |
| 209I_Multiplet | -2.54196771144989 | 10.1511106310496 | 0.004744562718552 |
| 217CO_Macrophage_Alveolar | -1.42080854397404 | 11.8151240903485 | 0.00174330716782805 |
| 221I_Macrophage | 3.52650162566722 | 8.88743594313994 | 2.41642718689321e-05 |
| 454C_Macrophage | 4.51116201065003 | 9.94697479350305 | 0.00242297915613859 |

--- gene_id: ENSG00000149021 gene_name: SCGB1A1 gene_location: 11:62405103-62423195 ---

| Set | LogFC | LogCPM | P-val |
|---|---|---|---|
| 133C_Multiplet | 4.52410633558452 | 9.47079773356648 | 0.000522500063477503 |
| 1372C_Multiplet | 8.18055375750624 | 12.4601526597791 | 8.11126472770336e-13 |
| 157I_Ciliated | -3.1049386425624 | 10.7282197155457 | 7.97906414616147e-31 |
| 184CO_ATII | 6.70042775623947 | 12.0780352716314 | 9.96818108557245e-11 |
| 184CO_Ciliated | -3.3284725316168 | 12.7443931650917 | 2.05662582393838e-13 |
| 235CO_Multiplet | 8.09463488493452 | 15.5914004788426 | 4.11297343763741e-13 |

--- gene_id: ENSG00000187193 gene_name: MT1X gene_location: 16:56682470-56684196 ---

| Set | LogFC | LogCPM | P-val |
|---|---|---|---|
| 056CO_Macrophage_Alveolar | 2.70598127820839 | 9.64023838651794 | 0.000183889262722767 |
| 178CO_cDC2 | -2.89525527811256 | 11.1776994631334 | 0.000119553472864631 |
| 217CO_Macrophage | 3.18465510427808 | 10.6079409155934 | 0.0037149231908925 |
| 217CO_Macrophage_Alveolar | -2.07857310298725 | 10.0263727293432 | 3.38622725874198e-06 |
| 253C_Macrophage_Alveolar | 3.73121441074635 | 7.58501572879332 | 0.0028939534529852 |

--- gene_id: ENSG00000125144 gene_name: MT1G gene_location: 16:56666730-56668065 ---

| Set | LogFC | LogCPM | P-val |
|---|---|---|---|
| 056CO_Macrophage_Alveolar | 2.69405606884449 | 9.38512950486033 | 0.00100324534054038 |
| 157I_Macrophage | -1.99531333331223 | 8.94269286202094 | 0.00308751298868151 |
| 209I_cDC2 | -4.3508347300711 | 8.710335721617 | 4.09010525807208e-07 |
| 217CO_Macrophage | 4.76389670970278 | 9.8797521312892 | 0.000850431780535936 |
| 217CO_Macrophage_Alveolar | -2.65431711615432 | 9.28970961873501 | 7.28162154531729e-08 |

--- gene_id: ENSG00000168878 gene_name: SFTPB gene_location: 2:85657314-85668741 ---

| Set | LogFC | LogCPM | P-val |
|---|---|---|---|
| 209I_Aberrant_Basaloid | -1.82419132134652 | 9.46101460981985 | 7.55539497311412e-06 |
| 209I_Ciliated | 4.44569220595912 | 9.05333776314033 | 8.13430986388814e-07 |
| 209I_Multiplet | 4.29600198913161 | 10.2872563695447 | 0.000279187694302017 |
| 235CO_Ciliated | -5.10182392204283 | 8.84576361906071 | 3.26042876164647e-05 |
| 235CO_Multiplet | -5.25489112022185 | 10.9489628071366 | 2.26910986347752e-09 |

--- gene_id: ENSG00000102970 gene_name: CCL17 gene_location: 16:57404767-57416063 ---

| Set | LogFC | LogCPM | P-val |
|---|---|---|---|
| 098C_cDC2 | 6.33505611492764 | 11.6948751538907 | 4.32280975497239e-19 |
| 133C_Multiplet | 4.3994466000366 | 8.49941020644921 | 0.00230688186386959 |
| 1372C_cDC2 | 3.39540801544107 | 11.3021552933055 | 3.39898642140363e-09 |
| 221I_cDC2 | 2.66485553568726 | 12.1596664336109 | 0.00341280164484718 |
| 235CO_Multiplet | -4.62767358231415 | 8.96955930493474 | 0.000494318099420154 |

--- gene_id: ENSG00000168484 gene_name: SFTPC gene_location: 8:22156913-22164479 ---

| Set | LogFC | LogCPM | P-val |
|---|---|---|---|
| 1372C_Multiplet | -7.54569230956838 | 12.7585866970377 | 1.30507842772322e-18 |
| 157I_Club | 2.67034862826178 | 8.55686156440657 | 0.000365777834490236 |
| 157I_Multiplet | 8.30125163754418 | 11.9839090596047 | 7.73224697910216e-29 |
| 184CO_Multiplet | 5.97627644199167 | 12.4211090386367 | 3.13991704716562e-07 |
| 235CO_Multiplet | -6.82239612404405 | 12.7106341143399 | 1.70527046994923e-16 |

--- gene_id: ENSG00000122852 gene_name: SFTPA1 gene_location: 10:79610939-79615455 ---

| Set | LogFC | LogCPM | P-val |
|---|---|---|---|
| 1372C_Multiplet | -7.19527646189248 | 10.1541934514738 | 2.40354433921501e-09 |
| 157I_Basal | -5.10927667512907 | 8.36666165441319 | 0.0079962372048717 |
| 157I_Multiplet | 5.13923242277202 | 9.42584121054854 | 1.03465374439501e-09 |
| 235CO_Multiplet | -6.16268252355631 | 10.2066070732629 | 1.39205273616004e-08 |

--- gene_id: ENSG00000115523 gene_name: GNLY gene_location: 2:85685175-85698852 ---

| Set | LogFC | LogCPM | P-val |
|---|---|---|---|
| 052CO_Multiplet | -3.50081559887437 | 10.3445574701195 | 0.000660448672987058 |
| 178CO_cDC2 | 5.90217131770144 | 10.5659577886378 | 8.97252131151386e-05 |
| 209I_Multiplet | -4.76918393785817 | 8.38141983302534 | 3.54270446423666e-06 |
| 225I_Multiplet | -4.79349299597472 | 8.39011202486788 | 1.87971676319412e-05 |

--- gene_id: ENSG00000130203 gene_name: APOE gene_location: 19:44905791-44909393 ---

| Set | LogFC | LogCPM | P-val |
|---|---|---|---|
| 133C_cDC2 | -1.90599409739671 | 10.0880327006861 | 0.00881593175172022 |
| 157I_Fibroblast | -3.69052604603726 | 10.1048618166854 | 0.00673976434196385 |
| 207CO_Macrophage | -3.18989357749351 | 11.4283645722952 | 3.23019238727816e-06 |
| 235CO_cMonocyte | 3.32284887199699 | 10.4839621568762 | 1.42046397563271e-06 |

--- gene_id: ENSG00000137077 gene_name: CCL21 gene_location: 9:34709005-34710136 ---

| Set | LogFC | LogCPM | P-val |
|---|---|---|---|
| 098C_Multiplet | 3.92668738655431 | 11.3346709595739 | 9.85985123738137e-11 |
| 133C_Multiplet | -3.92626347202662 | 7.92706842991192 | 9.62823200654892e-06 |
| 1372C_Multiplet | 7.30424039531406 | 12.5555915958687 | 1.67294179324559e-11 |
| 157I_Multiplet | 5.06676868149927 | 8.72329735843483 | 3.89908328099993e-07 |

--- gene_id: ENSG00000169715 gene_name: MT1E gene_location: 16:56625475-56627112 ---

| Set | LogFC | LogCPM | P-val |
|---|---|---|---|
| 052CO_Macrophage_Alveolar | -1.25504396606015 | 9.18946804616481 | 0.00325510562006196 |
| 178CO_Macrophage | -4.50754138142765 | 10.5097293063017 | 8.39553741808559e-06 |
| 184CO_ATII | 2.86703446234464 | 11.3583944674305 | 0.0037047581156464 |
| 217CO_Macrophage | 3.84694102795917 | 9.90542175659708 | 0.00518509102523681 |

--- gene_id: ENSG00000133063 gene_name: CHIT1 gene_location: 1:203212827-203273641 ---

| Set | LogFC | LogCPM | P-val |
|---|---|---|---|
| 056CO_Macrophage | 2.9822113673037 | 10.0893659135769 | 3.40835576303106e-05 |
| 178CO_Macrophage | 4.54924742142297 | 9.95633548965437 | 0.00100736887936437 |
| 235CO_Macrophage | 3.83545857221444 | 10.0389447345436 | 7.69756367063637e-07 |

--- gene_id: ENSG00000185303 gene_name: SFTPA2 gene_location: 10:79555852-79560407 ---

| Set | LogFC | LogCPM | P-val |
|---|---|---|---|
| 1372C_Multiplet | -5.32765882690004 | 9.69008765652356 | 1.5353411392877e-05 |
| 157I_Multiplet | 3.8623739987468 | 9.33386338481106 | 2.55026454004283e-06 |
| 235CO_Multiplet | -6.34363789425946 | 10.1786507218424 | 5.38368433936238e-09 |

--- gene_id: ENSG00000090382 gene_name: LYZ gene_location: 12:69348381-69354234 ---

| Set | LogFC | LogCPM | P-val |
|---|---|---|---|
| 052CO_T_Cytotoxic | -2.68541573274654 | 10.1342660125541 | 2.99833564016839e-05 |
| 157I_ncMonocyte | -1.0815857454593 | 11.4746909808411 | 0.0075616275599786 |
| 209I_cDC2 | 1.87680140489216 | 12.0368058356811 | 0.000105400831527448 |

--- gene_id: ENSG00000165949 gene_name: IFI27 gene_location: 14:94104836-94116695 ---

| Set | LogFC | LogCPM | P-val |
|---|---|---|---|
| 098C_Multiplet | -2.36488162482828 | 10.423262934322 | 0.000194309756395702 |
| 184CO_cDC1 | -3.65830622619862 | 9.2393045288924 | 5.6955441498546e-06 |
| 221I_cDC2 | 5.23158476226488 | 9.75659858158892 | 0.00611370235887504 |

--- gene_id: ENSG00000115009 gene_name: CCL20 gene_location: 2:227805739-227817564 ---

| Set | LogFC | LogCPM | P-val |
|---|---|---|---|
| 098C_cDC2 | 5.05594694520151 | 10.0369113506387 | 6.02862053159615e-06 |
| 157I_Club | -3.44664467041971 | 8.69275333036078 | 7.10487810418281e-08 |
| 253C_Macrophage_Alveolar | -1.60554363386516 | 7.73321278139332 | 0.00633634218204304 |

--- gene_id: ENSG00000102265 gene_name: TIMP1 gene_location: X:47582408-47586789 ---

| Set | LogFC | LogCPM | P-val |
|---|---|---|---|
| 1372C_Lymphatic | 1.46660346377552 | 12.000297753201 | 0.00243348245745791 |
| 1372C_VE_Venous | -1.67190679014478 | 12.2511856287367 | 0.00278168245669716 |
| 209I_cDC2 | 2.48583737941641 | 10.7345681238269 | 7.59588353912376e-06 |

--- gene_id: ENSG00000131400 gene_name: NAPSA gene_location: 19:50358472-50365830 ---

| Set | LogFC | LogCPM | P-val |
|---|---|---|---|
| 1372C_Multiplet | -4.16546219364405 | 9.0512171315906 | 0.00589467774897835 |
| 157I_Multiplet | 4.00018983095931 | 8.95620009892101 | 7.7397805554154e-06 |
| 235CO_Multiplet | -4.65333722875525 | 9.07912567990254 | 0.000354132074410219 |

--- gene_id: ENSG00000108691 gene_name: CCL2 gene_location: 17:34255274-34257208 ---

| Set | LogFC | LogCPM | P-val |
|---|---|---|---|
| 157I_Club | -3.65329227741824 | 8.25489835503994 | 4.0813217715611e-06 |
| 157I_cMonocyte | -5.75920909715986 | 10.2998228058247 | 8.98048834752025e-11 |
| 221I_Macrophage | -3.72405698752385 | 9.227762759793 | 5.37091932479867e-14 |

--- gene_id: ENSG00000152583 gene_name: SPARCL1 gene_location: 4:87473335-87531061 ---

| Set | LogFC | LogCPM | P-val |
|---|---|---|---|
| 1372C_ATII | -3.7629886883274 | 7.48702337788925 | 0.000128351892514856 |
| 157I_Multiplet | 3.71575577896562 | 8.40256738117344 | 0.00352628712715084 |
| 184CO_Multiplet | -3.49972554398695 | 9.16224248378473 | 0.000710789359292581 |

--- gene_id: ENSG00000100234 gene_name: TIMP3 gene_location: 22:32801705-32863041 ---

| Set | LogFC | LogCPM | P-val |
|---|---|---|---|
| 052CO_Macrophage | 4.21319229411336 | 9.09069687693313 | 0.000238719297323189 |
| 157I_Multiplet | 3.7531657423653 | 8.35111003003344 | 0.0035650506077915 |

```
225I_cDC2                3.91090491464741          8.73832124275654               5.89007923484359e-05
--- gene_id: ENSG00000173432 gene_name: SAA1 gene_location: 11:18266260-18269977 ---
Set                      LogFC                     LogCPM                         P-val
157I_Ciliated            -1.69605388627823         8.71294501794338               0.00831248395325181
209I_Ciliated            -1.96739123396679         10.3213294229234               4.59899226966778e-07
235CO_Multiplet          5.89962548575956          9.83238661262514               7.53848178729479e-05
--- gene_id: ENSG00000125148 gene_name: MT2A gene_location: 16:56608584-56609497 ---
Set                      LogFC                     LogCPM                         P-val
184CO_ATII               3.53606545314932          12.0604880396629               5.34126081246805e-05
184CO_DC_Mature          -1.40902239720875         13.0102349913577               0.000746478824683256
439C_cMonocyte           -3.87161898739974         12.6423670508962               1.42360026197777e-07
--- gene_id: ENSG00000164266 gene_name: SPINK1 gene_location: 5:147824572-147831671 ---
Set                      LogFC                     LogCPM                         P-val
052CO_Multiplet          -3.90970423527493         9.42100588013103               0.00104858204152736
209I_Aberrant_Basaloid   -2.82970374986926         8.92450566912713               1.57845115682969e-12
--- gene_id: ENSG00000227507 gene_name: LTB gene_location: 6:31580525-31582522 ---
Set                      LogFC                     LogCPM                         P-val
157I_Multiplet           -3.31698190666529         8.34820852911586               0.00672568056984869
209I_Aberrant_Basaloid   1.92001450659668          8.73807282356313               0.00123786388366105
--- gene_id: ENSG00000173110 gene_name: HSPA6 gene_location: 1:161524540-161526894 ---
Set                      LogFC                     LogCPM                         P-val
209I_Aberrant_Basaloid   -2.14154889249679         8.54856482071373               4.87341369657539e-06
225I_cDC2                -3.06535485712317         8.48521812980279               0.00267080398780062
--- gene_id: ENSG00000277632 gene_name: CCL3 gene_location: 17:36088256-36090169 ---
Set                      LogFC                     LogCPM                         P-val
157I_cMonocyte           -3.55238194966383         9.66322141097485               0.00691424629057188
225I_Multiplet           -3.29936670107183         9.35587491654456               0.000708246037625539
--- gene_id: ENSG00000276070 gene_name: CCL4L2 gene_location: 17:36210924-36212878 ---
Set                      LogFC                     LogCPM                         P-val
209I_cDC2                -4.08698113957136         8.81370516665572               4.09010525807208e-07
225I_Multiplet           -3.76325169987112         9.38054549694921               3.74310742022393e-05
--- gene_id: ENSG00000164687 gene_name: FABP5 gene_location: 8:81280536-81284777 ---
Set                      LogFC                     LogCPM                         P-val
157I_Ciliated            -2.28552887983256         8.32683958988967               0.00325410597646085
184CO_cDC1               -3.33165082851126         9.71319258244248               1.86674571567038e-08
--- gene_id: ENSG00000215182 gene_name: MUC5AC gene_location: 11:1157953-1201138 ---
Set                      LogFC                     LogCPM                         P-val
157I_Multiplet           -3.65820408017362         8.25044422463194               0.0035650506077915
209I_Goblet              -3.35155168720622         8.38290484010406               0.000274242401338889
--- gene_id: ENSG00000117984 gene_name: CTSD gene_location: 11:1752752-1764573 ---
Set                      LogFC                     LogCPM                         P-val
052CO_T_Cytotoxic        -2.45929783433988         10.0090772760775               7.58488004663767e-05
235CO_cMonocyte          2.32367737957703          10.6552798579493               0.000270781526765713
```

--- gene_id: ENSG00000138755 gene_name: CXCL9 gene_location: 4:76001275-76007509 ---

| Set | LogFC | LogCPM | P-val |
|---|---|---|---|
| 052CO_Macrophage | 3.58505736492189 | 9.14306913617095 | 0.00282407428903863 |
| 052CO_Macrophage_Alveolar | 2.10798893511213 | 8.19527707065335 | 0.00325510562006196 |

--- gene_id: ENSG00000118785 gene_name: SPP1 gene_location: 4:87975667-87983532 ---

| Set | LogFC | LogCPM | P-val |
|---|---|---|---|
| 098C_Multiplet | -5.65459523304397 | 9.16830277765296 | 9.38292496829022e-11 |
| 157I_Basal | -4.50032352813705 | 10.4325497302512 | 0.00119202143903428 |

--- gene_id: ENSG00000163735 gene_name: CXCL5 gene_location: 4:73995642-73998677 ---

| Set | LogFC | LogCPM | P-val |
|---|---|---|---|
| 217CO_Macrophage | 4.250300451279 | 9.65824786011365 | 0.00518509102523681 |
| 235CO_cMonocyte | -2.85240793572309 | 9.84238285357989 | 0.00462025932436101 |

--- gene_id: ENSG00000133661 gene_name: SFTPD gene_location: 10:79937467-79982614 ---

| Set | LogFC | LogCPM | P-val |
|---|---|---|---|
| 1372C_Multiplet | -3.80751403786718 | 9.20219330352714 | 0.00809289713795883 |
| 235CO_Multiplet | -4.55234493982481 | 9.09487169320613 | 0.00044284511145804 |

--- gene_id: ENSG00000100453 gene_name: GZMB gene_location: 14:24630954-24634267 ---

| Set | LogFC | LogCPM | P-val |
|---|---|---|---|
| 209I_Multiplet | -3.52187341487346 | 8.04664265478437 | 0.00909991125066625 |
| 225I_Multiplet | -3.87214741560063 | 7.93373048500202 | 0.00267106199140038 |

--- gene_id: ENSG00000164972 gene_name: C9orf24 gene_location: 9:34379019-34397828 ---

| Set | LogFC | LogCPM | P-val |
|---|---|---|---|
| 209I_Aberrant_Basaloid | -3.75025328790802 | 7.42460487730683 | 1.36755660004719e-06 |
| 235CO_Multiplet | 5.01629688057177 | 9.51193752027888 | 0.00195729216641528 |

--- gene_id: ENSG00000163453 gene_name: IGFBP7 gene_location: 4:57030773-57110385 ---

| Set | LogFC | LogCPM | P-val |
|---|---|---|---|
| 157I_Club | -3.17861659226046 | 7.80965488104786 | 0.00308836457940088 |
| 157I_Multiplet | 2.87475921388853 | 9.61774179782084 | 0.000902623049522971 |

--- gene_id: ENSG00000166681 gene_name: BEX3 gene_location: X:103376395-103378164 ---

| Set | LogFC | LogCPM | P-val |
|---|---|---|---|
| 098C_Macrophage_Alveolar | 0.629439635840284 | 7.86149002798803 | 0.00211768526567966 |
| 133C_Macrophage_Alveolar | 1.39797724699611 | 8.00536251289645 | 3.93349592028239e-06 |

--- gene_id: ENSG00000275302 gene_name: CCL4 gene_location: 17:36103827-36105621 ---

| Set | LogFC | LogCPM | P-val |
|---|---|---|---|
| 184CO_Ciliated | -3.7864482445597 | 9.34649559047581 | 1.63937094046031e-05 |
| 225I_Multiplet | -5.22956464904619 | 9.73847395367271 | 3.82880358648697e-10 |

--- gene_id: ENSG00000069482 gene_name: GAL gene_location: 11:68683779-68691175 ---

| Set | LogFC | LogCPM | P-val |
|---|---|---|---|
| 174I_Macrophage | -2.37892798297277 | 7.93101493487779 | 0.00116787154805111 |
| 225I_Macrophage | -1.72217977459764 | 8.44889472467049 | 0.00326912439541166 |

--- gene_id: ENSG00000105374 gene_name: NKG7 gene_location: 19:51371606-51372701 ---

| Set | LogFC | LogCPM | P-val |
|---|---|---|---|
| 209I_Multiplet | -4.32347252998671 | 8.28492329479981 | 8.94197679367338e-05 |

| | | | |
|---|---|---|---|
| 225I_Multiplet | -5.10997721555364 | 8.23137144083545 | 1.25630212173563e-06 |

--- gene_id: ENSG00000142748 gene_name: FCN3 gene_location: 1:27369110-27374824 ---

| Set | LogFC | LogCPM | P-val |
|---|---|---|---|
| 052CO_Multiplet | -3.73554437071576 | 10.1555483076486 | 0.000532710166526524 |
| 1372C_Multiplet | 5.29116227151274 | 9.25800180890659 | 0.00301506040948703 |

--- gene_id: ENSG00000125999 gene_name: BPIFB1 gene_location: 20:33273480-33309871 ---

| Set | LogFC | LogCPM | P-val |
|---|---|---|---|
| 157I_Ciliated | 1.70679390527875 | 8.87532880636263 | 0.00564776505249782 |
| 235CO_Multiplet | 5.61639157686687 | 9.81625495549186 | 0.000257305220137281 |

--- gene_id: ENSG00000263639 gene_name: MSMB gene_location: 10:46033307-46048180 ---

| Set | LogFC | LogCPM | P-val |
|---|---|---|---|
| 209I_Goblet | -4.39610662084469 | 13.2559673758211 | 3.23799461739329e-14 |
| 235CO_Multiplet | 5.63665612866886 | 10.6083854469702 | 1.1862769072502e-05 |

--- gene_id: ENSG00000011465 gene_name: DCN gene_location: 12:91140484-91183217 ---

| Set | LogFC | LogCPM | P-val |
|---|---|---|---|
| 184CO_Multiplet | -5.11812894635497 | 9.79519944838052 | 2.31083592162409e-07 |
| 235CO_Multiplet | 4.74598966333717 | 10.1689329166973 | 0.00744244676207334 |

--- gene_id: ENSG00000160180 gene_name: TFF3 gene_location: 21:42311667-42315409 ---

| Set | LogFC | LogCPM | P-val |
|---|---|---|---|
| 098C_Multiplet | 3.85568436173771 | 9.12025049429577 | 0.000121976604033442 |
| 209I_Goblet | -5.04759154362271 | 10.9144844330105 | 5.81038970780734e-17 |

--- gene_id: ENSG00000107317 gene_name: PTGDS gene_location: 9:136975092-136981742 ---

| Set | LogFC | LogCPM | P-val |
|---|---|---|---|
| 052CO_Fibroblast | 2.71931541551487 | 11.9553549077022 | 0.000337315414071524 |
| 157I_Fibroblast | -3.81937309996729 | 12.2200533883345 | 3.49480176243205e-08 |

--- gene_id: ENSG00000149591 gene_name: TAGLN gene_location: 11:117199370-117207464 ---

| Set | LogFC | LogCPM | P-val |
|---|---|---|---|
| 1372C_ATII | -2.67360428675241 | 7.61718072481987 | 0.00827838261252021 |
| 1372C_VE_Venous | -3.21288840264873 | 10.5011153581597 | 4.26260414609543e-05 |

--- gene_id: ENSG00000189058 gene_name: APOD gene_location: 3:195568705-195584033 ---

| Set | LogFC | LogCPM | P-val |
|---|---|---|---|
| 157I_Fibroblast | -3.26774483606691 | 10.6160826643542 | 0.00363560488186653 |
| 225I_VE_Peribronchial | -3.22533451625951 | 8.18914417879239 | 5.37474142894096e-06 |

--- gene_id: ENSG00000163739 gene_name: CXCL1 gene_location: 4:73869393-73871308 ---

| Set | LogFC | LogCPM | P-val |
|---|---|---|---|
| 133C_cDC2 | -2.86987735982957 | 8.18674807027331 | 0.00321060902004344 |
| 235CO_Ciliated | -4.3954915415059 | 8.56611916671505 | 0.00435238555074835 |

--- gene_id: ENSG00000145824 gene_name: CXCL14 gene_location: 5:135570679-135579279 ---

| Set | LogFC | LogCPM | P-val |
|---|---|---|---|
| 157I_Fibroblast | -2.52906843419757 | 11.2750928916566 | 0.00673976434196385 |

--- gene_id: ENSG00000124731 gene_name: TREM1 gene_location: 6:41267926-41286682 ---

| Set | LogFC | LogCPM | P-val |
|---|---|---|---|
| 209I_cDC2 | 3.25005899745458 | 9.12047655803191 | 0.000448426397263157 |

```
--- gene_id: ENSG00000124237 gene_name: C20orf85 gene_location: 20:58150902-58161150 ---
Set                         LogFC                     LogCPM                         P-val
209I_Aberrant_Basaloid      -3.08760352437869         7.42151477405081               0.000117170142457616
--- gene_id: ENSG00000163734 gene_name: CXCL3 gene_location: 4:74036589-74038807 ---
Set                         LogFC                     LogCPM                         P-val
235CO_cDC2                  4.34804306296902          9.83922049538204               0.00021878835418558
--- gene_id: ENSG00000173369 gene_name: C1QB gene_location: 1:22652762-22661637 ---
Set                         LogFC                     LogCPM                         P-val
184CO_cDC1                  -3.10335249658321         9.36641523678747               5.11840556895574e-05
--- gene_id: ENSG00000205420 gene_name: KRT6A gene_location: 12:52487176-52493257 ---
Set                         LogFC                     LogCPM                         P-val
1372C_ATII                  -4.36665959345561         7.50293447521018               9.06370950995382e-07
--- gene_id: ENSG00000131724 gene_name: IL13RA1 gene_location: X:118727133-118794535 ---
Set                         LogFC                     LogCPM                         P-val
098C_Macrophage_Alveolar    -0.68400043722238         7.98762039628653               1.45722640586237e-15
--- gene_id: ENSG00000159167 gene_name: STC1 gene_location: 8:23841929-23854806 ---
Set                         LogFC                     LogCPM                         P-val
1372C_Lymphatic             -3.28746033463015         9.51308745337894               0.000186356231588742
--- gene_id: ENSG00000157601 gene_name: MX1 gene_location: 21:41420020-41470071 ---
Set                         LogFC                     LogCPM                         P-val
253C_Macrophage_Alveolar    -1.66382257738932         7.73814027696952               0.00239055158455849
--- gene_id: ENSG00000234906 gene_name: APOC2 gene_location: 19:44946035-44949565 ---
Set                         LogFC                     LogCPM                         P-val
052CO_Macrophage            -3.34479318802663         9.07747447581849               0.0076437822970523
--- gene_id: ENSG00000132507 gene_name: EIF5A gene_location: 17:7306999-7312463 ---
Set                         LogFC                     LogCPM                         P-val
1372C_ATII                  -1.90652753853053         9.17771488536395               0.00286167091480583
--- gene_id: ENSG00000160307 gene_name: S100B gene_location: 21:46598604-46605208 ---
Set                         LogFC                     LogCPM                         P-val
209I_cDC2                   -1.78693021582056         10.7610249212222               0.000806778751032012
--- gene_id: ENSG00000123610 gene_name: TNFAIP6 gene_location: 2:151357592-151380046 ---
Set                         LogFC                     LogCPM                         P-val
052CO_Macrophage_Alveolar   2.21428494750228          8.10165900270591               0.00433471305676568
--- gene_id: ENSG00000175899 gene_name: A2M gene_location: 12:9067664-9116229 ---
Set                         LogFC                     LogCPM                         P-val
052CO_Fibroblast            4.46892359510037          9.9767673354823                0.00499461079065134
--- gene_id: ENSG00000182853 gene_name: VMO1 gene_location: 17:4785285-4786433 ---
Set                         LogFC                     LogCPM                         P-val
098C_Macrophage_Alveolar    -0.628311051677203        8.03036761661385               1.6172745651372e-09
--- gene_id: ENSG00000126264 gene_name: HCST gene_location: 19:35902529-35904377 ---
Set                         LogFC                     LogCPM                         P-val
209I_cDC2                   2.33134543026617          9.46343520407008               0.00813021969049639
--- gene_id: ENSG00000277734 gene_name: TRAC gene_location: 14:22547506-22552156 ---
```

| Set | LogFC | LogCPM | P-val |
|---|---|---|---|
| 174I_cDC2 | -3.05488715121218 | 8.31337987302657 | 0.000155515162899085 |

--- gene_id: ENSG00000170458 gene_name: CD14 gene_location: 5:140631728-140633700 ---

| Set | LogFC | LogCPM | P-val |
|---|---|---|---|
| 209I_cDC2 | 3.60832580077838 | 9.51149749390338 | 5.85643344829337e-06 |

--- gene_id: ENSG00000148346 gene_name: LCN2 gene_location: 9:128149071-128153453 ---

| Set | LogFC | LogCPM | P-val |
|---|---|---|---|
| 209I_Goblet | -2.43525967704949 | 12.8623073702804 | 0.000530562692513621 |

--- gene_id: ENSG00000197696 gene_name: NMB gene_location: 15:84655129-84658563 ---

| Set | LogFC | LogCPM | P-val |
|---|---|---|---|
| 217CO_Macrophage_Alveolar | -2.52251777226138 | 8.40331664282008 | 0.000888642877089693 |

--- gene_id: ENSG00000145287 gene_name: PLAC8 gene_location: 4:83090048-83137075 ---

| Set | LogFC | LogCPM | P-val |
|---|---|---|---|
| 209I_cDC2 | -3.07541453419815 | 9.38379801222586 | 2.94514631963117e-06 |

--- gene_id: ENSG00000163736 gene_name: PPBP gene_location: 4:73986439-73988190 ---

| Set | LogFC | LogCPM | P-val |
|---|---|---|---|
| 221I_Macrophage_Alveolar | 6.53940232770878 | 9.65726310514013 | 1.95184238995641e-12 |

--- gene_id: ENSG00000186352 gene_name: ANKRD37 gene_location: 4:185396021-185400628 ---

| Set | LogFC | LogCPM | P-val |
|---|---|---|---|
| 209I_cDC2 | 3.72468621863021 | 8.47274204561191 | 0.00528013999016113 |

--- gene_id: ENSG00000115461 gene_name: IGFBP5 gene_location: 2:216672105-216695549 ---

| Set | LogFC | LogCPM | P-val |
|---|---|---|---|
| 1372C_VE_Venous | 4.29371185765526 | 9.91897296253848 | 3.0607833990728e-05 |

--- gene_id: ENSG00000163106 gene_name: HPGDS gene_location: 4:94298535-94342876 ---

| Set | LogFC | LogCPM | P-val |
|---|---|---|---|
| 209I_cDC2 | -3.07683280382531 | 8.32748212884245 | 0.00975808284486498 |

--- gene_id: ENSG00000269927 gene_name: ENSG00000269927 gene_location: 14:71141125-71143253 ---

| Set | LogFC | LogCPM | P-val |
|---|---|---|---|
| 209I_Multiplet | -3.79054279141543 | 8.05119890800385 | 0.00600033954252768 |

--- gene_id: ENSG00000257017 gene_name: HP gene_location: 16:72054505-72061055 ---

| Set | LogFC | LogCPM | P-val |
|---|---|---|---|
| 1372C_Lymphatic | 4.64897533371766 | 10.018140398945 | 1.89575747226825e-08 |

--- gene_id: ENSG00000166920 gene_name: C15orf48 gene_location: 15:45430579-45448761 ---

| Set | LogFC | LogCPM | P-val |
|---|---|---|---|
| 052CO_Macrophage_Alveolar | 2.003708433522 | 8.22853297435913 | 0.00433471305676568 |

--- gene_id: ENSG00000123095 gene_name: BHLHE41 gene_location: 12:26120030-26125037 ---

| Set | LogFC | LogCPM | P-val |
|---|---|---|---|
| 235CO_Macrophage | 2.29974884846921 | 9.39993416591362 | 0.00323877707603465 |

--- gene_id: ENSG00000166825 gene_name: ANPEP gene_location: 15:89784895-89815401 ---

| Set | LogFC | LogCPM | P-val |
|---|---|---|---|
| 209I_cDC2 | 3.72423997531082 | 8.5017694648623 | 0.00528013999016113 |

--- gene_id: ENSG00000120708 gene_name: TGFBI gene_location: 5:136028988-136063818 ---

| Set | LogFC | LogCPM | P-val |
|---|---|---|---|

| | | | |
|---|---|---|---|
| 157I_Basal | 5.71856129453116 | 8.83879827013386 | 0.0051220953004933 |

--- gene_id: ENSG00000176435 gene_name: CLEC14A gene_location: 14:38254000-38256093 ---

| Set | LogFC | LogCPM | P-val |
|---|---|---|---|
| 225I_Multiplet | -3.47544856743646 | 8.1748352824174 | 0.00144761847766162 |

--- gene_id: ENSG00000204388 gene_name: HSPA1B gene_location: 6:31827738-31830254 ---

| Set | LogFC | LogCPM | P-val |
|---|---|---|---|
| 1372C_VE_Venous | -3.70151014155589 | 10.218400720069 | 3.86510379532803e-05 |

--- gene_id: ENSG00000103811 gene_name: CTSH gene_location: 15:78921058-78949574 ---

| Set | LogFC | LogCPM | P-val |
|---|---|---|---|
| 1372C_Multiplet | -4.27257121061732 | 9.22662255263257 | 0.00404708986927753 |

--- gene_id: ENSG00000196154 gene_name: S100A4 gene_location: 1:153543613-153550136 ---

| Set | LogFC | LogCPM | P-val |
|---|---|---|---|
| 1372C_Lymphatic | -2.48674545397547 | 9.58300765666674 | 0.0090779744490068 |

--- gene_id: ENSG00000139329 gene_name: LUM gene_location: 12:91102629-91111494 ---

| Set | LogFC | LogCPM | P-val |
|---|---|---|---|
| 052CO_Multiplet | 3.50711375674292 | 10.4908564636997 | 0.000660448672987058 |

--- gene_id: ENSG00000109321 gene_name: AREG gene_location: 4:74445136-74455005 ---

| Set | LogFC | LogCPM | P-val |
|---|---|---|---|
| 209I_Goblet | -3.47884544545339 | 8.05384519800245 | 0.000849697267122368 |

--- gene_id: ENSG00000104918 gene_name: RETN gene_location: 19:7669049-7670455 ---

| Set | LogFC | LogCPM | P-val |
|---|---|---|---|
| 052CO_Macrophage | -3.45158287221687 | 9.1357998905505 | 0.0052850189917585 |

--- gene_id: ENSG00000106819 gene_name: ASPN gene_location: 9:92456205-92482506 ---

| Set | LogFC | LogCPM | P-val |
|---|---|---|---|
| 225I_Myofibroblast | 1.76296937178574 | 9.41727196643716 | 0.00238136965355101 |

--- gene_id: ENSG00000136244 gene_name: IL6 gene_location: 7:22725884-22732002 ---

| Set | LogFC | LogCPM | P-val |
|---|---|---|---|
| 157I_Fibroblast | 6.3411835431491 | 10.8249631453567 | 3.61051678200589e-08 |

--- gene_id: ENSG00000131981 gene_name: LGALS3 gene_location: 14:55124110-55145423 ---

| Set | LogFC | LogCPM | P-val |
|---|---|---|---|
| 052CO_Macrophage | 2.2709332520366 | 10.3681952403976 | 0.00737840391433035 |

--- gene_id: ENSG00000197746 gene_name: PSAP gene_location: 10:71816298-71851251 ---

| Set | LogFC | LogCPM | P-val |
|---|---|---|---|
| 052CO_T_Cytotoxic | -2.27384783336098 | 10.0461065508533 | 0.00187374714213927 |

--- gene_id: ENSG00000135094 gene_name: SDS gene_location: 12:113392445-113426301 ---

| Set | LogFC | LogCPM | P-val |
|---|---|---|---|
| 052CO_cDC2 | 3.89049391453801 | 8.80927737255076 | 0.00135456770374844 |

--- gene_id: ENSG00000126353 gene_name: CCR7 gene_location: 17:40553769-40565472 ---

| Set | LogFC | LogCPM | P-val |
|---|---|---|---|
| 098C_cDC2 | -3.47571217838808 | 9.35543430939957 | 0.000268575303788251 |

--- gene_id: ENSG00000086062 gene_name: B4GALT1 gene_location: 9:33104082-33167356 ---

| Set | LogFC | LogCPM | P-val |
|---|---|---|---|
| 1372C_ATII | -2.30915507520234 | 8.23827239676574 | 0.00164570180803638 |

--- gene_id: ENSG00000151632 gene_name: AKR1C2 gene_location: 10:4987775-5018031 ---

| Set | LogFC | LogCPM | P-val |
|---|---|---|---|
| 157I_Club | -3.28718663198874 | 8.03063923853567 | 0.000409907458454968 |

--- gene_id: ENSG00000133422 gene_name: MORC2 gene_location: 22:30925130-30968774 ---

| Set | LogFC | LogCPM | P-val |
|---|---|---|---|
| 237CO_B | -2.66778490501996 | 10.4150415283749 | 0.00394845368640304 |

--- gene_id: ENSG00000167779 gene_name: IGFBP6 gene_location: 12:53097436-53102345 ---

| Set | LogFC | LogCPM | P-val |
|---|---|---|---|
| 184CO_Multiplet | -3.90608393092785 | 9.3570865111475 | 0.000553647322573952 |

--- gene_id: ENSG00000115380 gene_name: EFEMP1 gene_location: 2:55865967-55924139 ---

| Set | LogFC | LogCPM | P-val |
|---|---|---|---|
| 1372C_VE_Venous | -2.91373025306952 | 10.1744543685663 | 0.00228738226585994 |

--- gene_id: ENSG00000122133 gene_name: PAEP gene_location: 9:135561756-135566955 ---

| Set | LogFC | LogCPM | P-val |
|---|---|---|---|
| 209I_Aberrant_Basaloid | -3.29268144301597 | 7.18226682584894 | 0.000638814725227347 |

--- gene_id: ENSG00000164692 gene_name: COL1A2 gene_location: 7:94394895-94431227 ---

| Set | LogFC | LogCPM | P-val |
|---|---|---|---|
| 052CO_Multiplet | 4.13474613842649 | 9.76333517074891 | 0.000660448672987058 |

--- gene_id: ENSG00000102575 gene_name: ACP5 gene_location: 19:11574653-11579993 ---

| Set | LogFC | LogCPM | P-val |
|---|---|---|---|
| 052CO_T_Cytotoxic | -2.48115525288141 | 9.71665236514538 | 0.00935664099426964 |

--- gene_id: ENSG00000124766 gene_name: SOX4 gene_location: 6:21593751-21598619 ---

| Set | LogFC | LogCPM | P-val |
|---|---|---|---|
| 098C_cDC2 | 4.16877232550897 | 9.58879903446586 | 0.00230429820721498 |

--- gene_id: ENSG00000106483 gene_name: SFRP4 gene_location: 7:37905932-38025695 ---

| Set | LogFC | LogCPM | P-val |
|---|---|---|---|
| 1372C_Multiplet | -4.40357859628813 | 9.21984609335318 | 0.00312839978281918 |

--- gene_id: ENSG00000204389 gene_name: HSPA1A gene_location: 6:31815543-31817946 ---

| Set | LogFC | LogCPM | P-val |
|---|---|---|---|
| 235CO_cMonocyte | 2.86866226915225 | 9.90229916462628 | 0.00964828931102752 |

--- gene_id: ENSG00000163395 gene_name: IGFN1 gene_location: 1:201190824-201228952 ---

| Set | LogFC | LogCPM | P-val |
|---|---|---|---|
| 225I_Myofibroblast | -2.64961405045052 | 8.28635030698595 | 0.00186363286234534 |

--- gene_id: ENSG00000255112 gene_name: CHMP1B gene_location: 18:11851413-11854444 ---

| Set | LogFC | LogCPM | P-val |
|---|---|---|---|
| 209I_cDC2 | 2.13135654343555 | 9.97934774666827 | 0.00423413246917306 |

--- gene_id: ENSG00000081041 gene_name: CXCL2 gene_location: 4:74097040-74099196 ---

| Set | LogFC | LogCPM | P-val |
|---|---|---|---|
| 235CO_Ciliated | -4.0351492477869 | 9.22359442040835 | 4.79526359104692e-05 |

--- gene_id: ENSG00000118271 gene_name: TTR gene_location: 18:31557009-31598833 ---

| Set | LogFC | LogCPM | P-val |
|---|---|---|---|
| 157I_Club | -3.23436795831652 | 7.8252844417411 | 0.0024744418591124 |

--- gene_id: ENSG00000123689 gene_name: G0S2 gene_location: 1:209675412-209676390 ---

```
Set                           LogFC                     LogCPM                          P-val
439C_Macrophage               2.69830983607959          9.18906717562249                0.000198731497620243
--- gene_id: ENSG00000227591 gene_name: HSD11B1-AS1 gene_location: 1:209661356-209724125 ---
Set                           LogFC                     LogCPM                          P-val
184CO_DC_Mature               -3.00183638304378         9.44642196794688                0.00881728489370582
--- gene_id: ENSG00000087086 gene_name: FTL gene_location: 19:48965309-48966879 ---
Set                           LogFC                     LogCPM                          P-val
157I_Club                     -1.76207505819695         12.6611345025426                0.000127922285435058
--- gene_id: ENSG00000124882 gene_name: EREG gene_location: 4:74365145-74388749 ---
Set                           LogFC                     LogCPM                          P-val
209I_cDC2                     3.13097047094464          8.84106071207175                0.00379092615697161
--- gene_id: ENSG00000102760 gene_name: RGCC gene_location: 13:41457550-41470871 ---
Set                           LogFC                     LogCPM                          P-val
209I_cDC2                     2.03461378789956          10.1512813147333                0.00423413246917306
--- gene_id: ENSG00000012223 gene_name: LTF gene_location: 3:46435645-46485234 ---
Set                           LogFC                     LogCPM                          P-val
209I_Multiplet                5.98884064954033          9.5671045567765                 8.94197679367338e-05
--- gene_id: ENSG00000175701 gene_name: MTLN gene_location: 2:110211529-110245420 ---
Set                           LogFC                     LogCPM                          P-val
098C_Macrophage_Alveolar      -0.52813936459333         8.24164390060098                3.88827241946349e-12
--- gene_id: ENSG00000240583 gene_name: AQP1 gene_location: 7:30911853-30925517 ---
Set                           LogFC                     LogCPM                          P-val
157I_Multiplet                3.49803797101889          8.21389875300854                0.0093065208619798
--- gene_id: ENSG00000164047 gene_name: CAMP gene_location: 3:48223347-48225491 ---
Set                           LogFC                     LogCPM                          P-val
098C_Multiplet                -3.40695168271414         8.50118006203448                0.00871753089056714
--- gene_id: ENSG00000185745 gene_name: IFIT1 gene_location: 10:89392546-89406487 ---
Set                           LogFC                     LogCPM                          P-val
253C_Macrophage_Alveolar      -2.02702902749263         7.30851583189488                0.00239055158455849
--- gene_id: ENSG00000164287 gene_name: CDC20B gene_location: 5:55112971-55173177 ---
Set                           LogFC                     LogCPM                          P-val
157I_Ciliated                 2.87830787461837          8.20310221205965                0.00157990523669042
--- gene_id: ENSG00000100600 gene_name: LGMN gene_location: 14:92703807-92748679 ---
Set                           LogFC                     LogCPM                          P-val
052CO_Macrophage              3.77917585633033          9.48697440689669                0.000159667153742855
--- gene_id: ENSG00000108821 gene_name: COL1A1 gene_location: 17:50184101-50201632 ---
Set                           LogFC                     LogCPM                          P-val
157I_Basal                    6.32107233108523          9.24537481743012                0.00119202143903428
--- gene_id: ENSG00000197353 gene_name: LYPD2 gene_location: 8:142750150-142752532 ---
Set                           LogFC                     LogCPM                          P-val
157I_ncMonocyte               3.18154079238957          9.43901123661212                0.00634362129652463
--- gene_id: ENSG00000169245 gene_name: CXCL10 gene_location: 4:76021118-76023497 ---
Set                           LogFC                     LogCPM                          P-val
```

| | | | |
|---|---|---|---|
| 221I_cDC2 | 5.78736254726823 | 10.1036075100091 | 0.00105592585365135 |

--- gene_id: ENSG00000120885 gene_name: CLU gene_location: 8:27596917-27614700 ---

| Set | LogFC | LogCPM | P-val |
|---|---|---|---|
| 157I_Multiplet | 2.86392879488089 | 8.93906605530596 | 0.00613148051884766 |

--- gene_id: ENSG00000137801 gene_name: THBS1 gene_location: 15:39581079-39599466 ---

| Set | LogFC | LogCPM | P-val |
|---|---|---|---|
| 209I_cDC2 | 2.75014103682816 | 8.87543810763968 | 0.00975808284486498 |

--- gene_id: ENSG00000136689 gene_name: IL1RN gene_location: 2:113099315-113134016 ---

| Set | LogFC | LogCPM | P-val |
|---|---|---|---|
| 209I_cMonocyte | -4.71891280689621 | 9.63161315692499 | 0.0020162945668261 |

--- gene_id: ENSG00000008517 gene_name: IL32 gene_location: 16:3065297-3082192 ---

| Set | LogFC | LogCPM | P-val |
|---|---|---|---|
| 207CO_Macrophage | -3.47239292347243 | 9.57249272159828 | 0.00426845506349222 |

--- gene_id: ENSG00000129538 gene_name: RNASE1 gene_location: 14:20801228-20802855 ---

| Set | LogFC | LogCPM | P-val |
|---|---|---|---|
| 235CO_Multiplet | -4.18925498539278 | 9.35832186428895 | 0.000266678117654338 |

--- gene_id: ENSG00000130208 gene_name: APOC1 gene_location: 19:44914247-44919349 ---

| Set | LogFC | LogCPM | P-val |
|---|---|---|---|
| 207CO_Macrophage | -3.11272198658354 | 11.1924878218076 | 9.61110520206033e-06 |

--- gene_id: ENSG00000204866 gene_name: IGFL2 gene_location: 19:46143106-46161298 ---

| Set | LogFC | LogCPM | P-val |
|---|---|---|---|
| 209I_Aberrant_Basaloid | 2.85740906980821 | 7.68721046232135 | 0.00540044624320199 |

--- gene_id: ENSG00000106341 gene_name: PPP1R17 gene_location: 7:31687215-31708455 ---

| Set | LogFC | LogCPM | P-val |
|---|---|---|---|
| 184CO_ncMonocyte | 0.860080850459067 | 9.64692211901496 | 0.00967414798536446 |

--- gene_id: ENSG00000205364 gene_name: MT1M gene_location: 16:56632659-56633981 ---

| Set | LogFC | LogCPM | P-val |
|---|---|---|---|
| 184CO_Multiplet | -3.67028201558575 | 9.2916123614272 | 0.000553647322573952 |

--- gene_id: ENSG00000085265 gene_name: FCN1 gene_location: 9:134903232-134917912 ---

| Set | LogFC | LogCPM | P-val |
|---|---|---|---|
| 209I_cDC2 | 5.05043978229317 | 9.61885874289857 | 6.20822867332865e-09 |

--- gene_id: ENSG00000164104 gene_name: HMGB2 gene_location: 4:173331376-173334432 ---

| Set | LogFC | LogCPM | P-val |
|---|---|---|---|
| 157I_Club | 2.71885046174433 | 8.13730710016817 | 0.00548368568720831 |

--- gene_id: ENSG00000118523 gene_name: CCN2 gene_location: 6:131948176-131951372 ---

| Set | LogFC | LogCPM | P-val |
|---|---|---|---|
| 209I_Aberrant_Basaloid | 1.81292850267849 | 8.76501716497352 | 0.00535121382278162 |

--- gene_id: ENSG00000138433 gene_name: CIR1 gene_location: 2:174348022-174395712 ---

| Set | LogFC | LogCPM | P-val |
|---|---|---|---|
| 178CO_Macrophage | 4.03938767824445 | 10.4933619778961 | 0.000559688178497077 |

--- gene_id: ENSG00000250722 gene_name: SELENOP gene_location: 5:42799880-42887392 ---

| Set | LogFC | LogCPM | P-val |
|---|---|---|---|
| 217CO_Macrophage | -3.39826213558709 | 9.7401391036619 | 0.00659588631340416 |

--- gene_id: ENSG00000169429 gene_name: CXCL8 gene_location: 4:73740519-73743716 ---

| Set | LogFC | LogCPM | P-val |
|---|---|---|---|
| 1372C_cDC2 | -1.95240319628896 | 10.8541222343315 | 0.00198940554915985 |

--- gene_id: ENSG00000198417 gene_name: MT1F gene_location: 16:56657731-56660698 ---

| Set | LogFC | LogCPM | P-val |
|---|---|---|---|
| 217CO_Macrophage_Alveolar | -2.43182209265084 | 8.43314986179474 | 0.00149943607627406 |

--- gene_id: ENSG00000163220 gene_name: S100A9 gene_location: 1:153357854-153361023 ---

| Set | LogFC | LogCPM | P-val |
|---|---|---|---|
| 209I_cDC2 | 5.4822914341429 | 9.95428030236718 | 4.89330677257677e-11 |

--- gene_id: ENSG00000134539 gene_name: KLRD1 gene_location: 12:10226058-10329608 ---

| Set | LogFC | LogCPM | P-val |
|---|---|---|---|
| 225I_Multiplet | -3.74313099845997 | 7.85292116559184 | 0.00231823254745899 |

--- gene_id: ENSG00000106178 gene_name: CCL24 gene_location: 7:75810825-75823356 ---

| Set | LogFC | LogCPM | P-val |
|---|---|---|---|
| 221I_Macrophage | -2.63182073135721 | 8.41924780356643 | 0.00010259013643767 |

--- gene_id: ENSG00000187608 gene_name: ISG15 gene_location: 1:1001138-1014540 ---

| Set | LogFC | LogCPM | P-val |
|---|---|---|---|
| 209I_Goblet | -3.15710722411417 | 8.23445374957615 | 0.00302849109399017 |

--- gene_id: ENSG00000128383 gene_name: APOBEC3A gene_location: 22:38952741-38992778 ---

| Set | LogFC | LogCPM | P-val |
|---|---|---|---|
| 209I_Multiplet | -3.69143175583808 | 8.03647991981848 | 0.00909991125066625 |

--- gene_id: ENSG00000181195 gene_name: PENK gene_location: 8:56436674-56446671 ---

| Set | LogFC | LogCPM | P-val |
|---|---|---|---|
| 1372C_Lymphatic | -3.34071181288766 | 9.40356554849647 | 0.000382751393511191 |

--- gene_id: ENSG00000213088 gene_name: ACKR1 gene_location: 1:159203307-159206500 ---

| Set | LogFC | LogCPM | P-val |
|---|---|---|---|
| 225I_Multiplet | -4.43146163153008 | 8.35588102399344 | 1.87971676319412e-05 |

--- gene_id: ENSG00000262406 gene_name: MMP12 gene_location: 11:102862736-102874982 ---

| Set | LogFC | LogCPM | P-val |
|---|---|---|---|
| 174I_Macrophage | -4.03183629039286 | 8.2574403250938 | 1.26341454590244e-12 |

--- gene_id: ENSG00000184292 gene_name: TACSTD2 gene_location: 1:58575433-58577252 ---

| Set | LogFC | LogCPM | P-val |
|---|---|---|---|
| 225I_cDC2 | -3.08545472614066 | 8.47413226430913 | 0.00126988713131812 |

--- gene_id: ENSG00000198804 gene_name: MT-CO1 gene_location: MT:5904-7445 ---

| Set | LogFC | LogCPM | P-val |
|---|---|---|---|
| 052CO_T_Cytotoxic | -0.860557706359398 | 13.4873755462633 | 0.00935664099426964 |

--- gene_id: ENSG00000133048 gene_name: CHI3L1 gene_location: 1:203178931-203186704 ---

| Set | LogFC | LogCPM | P-val |
|---|---|---|---|
| 052CO_Macrophage | 4.83933030414875 | 9.35881372303382 | 3.30601175820177e-06 |

--- gene_id: ENSG00000143546 gene_name: S100A8 gene_location: 1:153390032-153391073 ---

| Set | LogFC | LogCPM | P-val |
|---|---|---|---|
| 209I_cDC2 | 4.65245922034763 | 9.72794621486532 | 7.97467348242523e-09 |

--- gene_id: ENSG00000124102 gene_name: PI3 gene_location: 20:45174902-45176544 ---

```
Set                         LogFC                     LogCPM                        P-val
1372C_ATII                  -3.59698889083684         8.00714835482817              6.0875087921971e-08
--- gene_id: ENSG00000117318 gene_name: ID3 gene_location: 1:23557926-23559501 ---
Set                         LogFC                     LogCPM                        P-val
209I_cDC2                   3.34446008731685          8.96851365741477              0.000806778751032012
--- gene_id: ENSG00000257764 gene_name: ENSG00000257764 gene_location: 12:69353493-69354225 ---
Set                         LogFC                     LogCPM                        P-val
209I_cDC2                   2.39099872926963          9.51365313025691              0.00406840539670693
--- gene_id: ENSG00000261040 gene_name: WFDC21P gene_location: 17:60083562-60091885 ---
Set                         LogFC                     LogCPM                        P-val
253C_Macrophage_Alveolar    -1.84348633600149         7.39151597054049              0.00633634218204304
--- gene_id: ENSG00000163993 gene_name: S100P gene_location: 4:6693878-6697170 ---
Set                         LogFC                     LogCPM                        P-val
157I_Club                   3.09414023448772          7.88202524608283              0.00601948915458148
--- gene_id: ENSG00000143387 gene_name: CTSK gene_location: 1:150794880-150809577 ---
Set                         LogFC                     LogCPM                        P-val
157I_Basal                  -6.57236034039487         9.30313733044806              0.000151443475793639
--- gene_id: ENSG00000160789 gene_name: LMNA gene_location: 1:156082573-156140081 ---
Set                         LogFC                     LogCPM                        P-val
209I_cDC2                   2.56324000762549          9.71250487669527              0.000999139152028508
--- gene_id: ENSG00000169554 gene_name: ZEB2 gene_location: 2:144364364-144521057 ---
Set                         LogFC                     LogCPM                        P-val
209I_cDC2                   3.93588597278262          8.55946330006788              0.00193249401925017
--- gene_id: ENSG00000276085 gene_name: CCL3L1 gene_location: 17:36194869-36196758 ---
Set                         LogFC                     LogCPM                        P-val
225I_Multiplet              -3.87547770126287         8.76000117856909              1.64325840008385e-05
--- gene_id: ENSG00000245532 gene_name: NEAT1 gene_location: 11:65422774-65445540 ---
Set                         LogFC                     LogCPM                        P-val
056CO_T_Cytotoxic           -3.03653832204289         12.7195536441098              0.000252621302966368
--- gene_id: ENSG00000232629 gene_name: HLA-DQB2 gene_location: 6:32756098-32763532 ---
Set                         LogFC                     LogCPM                        P-val
225I_cDC2                   -2.22379967935647         8.83834209760049              0.00267080398780062
--- gene_id: ENSG00000138207 gene_name: RBP4 gene_location: 10:93591687-93601744 ---
Set                         LogFC                     LogCPM                        P-val
056CO_Macrophage_Alveolar   1.8843635198444           9.58357502168413              0.00869124583600962
--- gene_id: ENSG00000102393 gene_name: GLA gene_location: X:101393273-101408012 ---
Set                         LogFC                     LogCPM                        P-val
098C_Macrophage_Alveolar    -0.42315443469197         7.96722391129104              0.00101265309166126
--- gene_id: ENSG00000189223 gene_name: PAX8-AS1 gene_location: 2:113211421-113276581 ---
Set                         LogFC                     LogCPM                        P-val
209I_Macrophage             1.28743495683592          7.77743639031302              9.92186336417071e-09
```
