## Supplemental Table 6 for "Single-cell X-chromosome inactivation analysis links biased chimerism to differential gene expression and epigenetic erosion"

--- gene_id: ENSG00000130988 gene_name: RGN gene_location: X:47078355-47093314 ---

| Set | LogFC | LogCPM | P-val |
|---|---|---|---|
| FTD073_FC_Astrocytes2 | -0.864731748921502 | 10.6570631725459 | 3.19295909393543e-05 |
| CTR081_TC_Astrocytes1 | -0.612424402477503 | 11.1462582337334 | 1.03918496136109e-07 |
| CTR081_FC_Astrocytes1 | -0.503739692178465 | 11.4089481375491 | 1.21492418487747e-10 |
| IB5431_OTC_excitatory | -0.665134179177575 | 9.74089909729238 | 2.48255337707705e-08 |
| IB6001_OC_astrocytes | -0.931332387503176 | 10.9723512908229 | 1.17694122130102e-13 |
| FTD014_FC_Astrocytes1 | -1.26352443082106 | 10.6193994200998 | 3.55156843648279e-17 |
| FTD073_FC_Astrocytes1 | -1.02023461061573 | 10.6845528537087 | 3.28961634915076e-18 |
| FTD014_TC_Astrocytes2 | -1.52866099971427 | 9.96863769180006 | 2.4322160575256e-21 |
| IB5431_OTC_astrocytes | -1.36338510374179 | 10.5231753231797 | 2.10447424787078e-35 |
| IB5431_CB_Bergmann1 | -1.0627007540495 | 11.3488033371719 | 1.15730605711562e-11 |
| IB5431_OC_astrocytes | -0.973752098759503 | 10.5438226905954 | 6.20459916773126e-20 |
| FTD014_TC_Astrocytes1 | -1.43720939327605 | 10.0923716314253 | 4.82432193572997e-35 |
| FTD083_TC_Astrocytes1 | 1.00351299280087 | 10.5461888781004 | 2.01296333363816e-05 |
| IB5431_CB_Fibroblasts | -3.31658897390921 | 13.8440398085369 | 0.00884184393501855 |
| FTD073_TC_Astrocytes2 | -0.844538144300179 | 10.4711964036176 | 6.92336270400841e-05 |
| IB5691_OC_astrocytes | -0.793388555350467 | 11.2291889287215 | 6.15079432630017e-05 |
| IB6001_OTC_astrocytes | -1.38551556242126 | 10.5643559430654 | 6.35797127136192e-17 |
| IB5431_CB_Astrocytes | -1.61290090372187 | 11.5085940992608 | 0.00124480507433553 |
| IB5691_OTC_astrocytes | -0.94596520500889 | 10.748161522025 | 5.43105395471611e-08 |
| FTD073_TC_Astrocytes1 | -1.53794901501185 | 10.108668708617 | 9.83041320606011e-24 |

--- gene_id: ENSG00000102362 gene_name: SYTL4 gene_location: X:100671783-100732123 ---

| Set | LogFC | LogCPM | P-val |
|---|---|---|---|
| IB5431_OC_astrocytes | -0.663191925325933 | 10.6644491134876 | 5.43710347344549e-08 |
| IB5869_OC_astrocytes | -0.805318703292856 | 10.8364151192295 | 2.91059209887768e-09 |
| FTD083_TC_Astrocytes1 | -1.0374075458266 | 10.5123759979871 | 7.14483802566274e-06 |
| FTD014_TC_Astrocytes1 | 1.14838705325396 | 10.2210345496862 | 1.16673110026086e-22 |
| IB5869_OTC_astrocytes | -0.74994679065327 | 10.5363138935816 | 3.00377623329979e-06 |
| IB5508_OTC_astrocytes | 0.894095333382043 | 10.8474649429652 | 1.64643782883834e-14 |
| FTD014_FC_Astrocytes1 | 0.895558173838351 | 10.6914986517687 | 8.66308600607023e-08 |
| CTR018_FC_Astrocytes1 | -0.513534741637104 | 10.3547403137324 | 1.55595118718993e-12 |
| FTD014_TC_Astrocytes2 | 0.772885363418748 | 10.119105055962 | 2.6224645701183e-05 |
| IB5431_OTC_astrocytes | -0.690954537435951 | 10.5963544846177 | 6.31017270894574e-07 |
| CTR018_TC_Astrocytes1 | -0.75746305720691 | 10.4603777147691 | 6.09826730196859e-08 |
| IB5508_OC_astrocytes | 0.728757510234143 | 10.9595137832112 | 0.00192445779120371 |
| FTD014_FC_Astrocytes2 | 0.917376886627348 | 10.7425871258172 | 0.00157312006506741 |

--- gene_id: ENSG00000101883 gene_name: RHOXF1 gene_location: X:120109051-120245267 ---

| Set | LogFC | LogCPM | P-val |
|---|---|---|---|
| IB5431_OTC_astrocytes | -0.373984305541689 | 10.4368857335272 | 6.62635881242164e-07 |
| CTR018_FC_Astrocytes1 | -0.317651398095973 | 10.1859556215008 | 1.67381085112585e-16 |
| IB5431_OC_astrocytes | -0.281571097237608 | 10.4870837032901 | 0.00265593261548799 |
| FTD073_FC_Astrocytes1 | -0.24955851341774 | 10.6237877051824 | 3.84668129922642e-05 |

```
FTD073_TC_Astrocytes1          -0.475477733593493       10.0113454357688        2.04598170246383e-05
CTR018_TC_Astrocytes1          -0.342413272861048       10.29669763803          5.74953851185627e-05
IB5431_CB_Neurins_In_Purkinje  -0.47934076392445        10.4944555010425        1.00556710551651e-09
--- gene_id: ENSG00000235244 gene_name: DANT2 gene_location: X:115694880-115969148 ---
Set                            LogFC                    LogCPM                  P-val
IB5508_CB_Neurons_Ex           -1.08609113327093        11.6681150370766        0.00636333411660823
IB5431_CB_Bergmann1            0.774650413696785        11.5189683061962        2.46657699654102e-05
IB5508_CB_Bergmann1            -0.747125072990789       11.164670024673         7.20189732460755e-10
IB5869_CB_Neurins_In_Purkinje  0.844525614497645        10.9025789662931        4.22044993749502e-11
--- gene_id: ENSG00000245532 gene_name: NEAT1 gene_location: 11:65422774-65445540 ---
Set                            LogFC                    LogCPM                  P-val
FTD073_TC_Neurons              -2.56936964915941        10.9707677726242        0.0025255014521107
FTD083_FC_Neurons              -2.15069175061           11.4010276530637        7.51802470441908e-05
--- gene_id: ENSG00000110436 gene_name: SLC1A2 gene_location: 11:35251205-35420063 ---
Set                            LogFC                    LogCPM                  P-val
IB5508_OC_pericytes            -2.35837022490123        12.5621276378857        0.00503803543329482
CTR018_TC_Oligodendrocytes     -3.75901123779153        12.6893462265695        0.000223567287055581
--- gene_id: ENSG00000148513 gene_name: ANKRD30A gene_location: 10:37125598-37232567 ---
Set                            LogFC                    LogCPM                  P-val
IB5431_OC_neuron               2.07090514479481         9.97686105222279        0.000986353774208035
--- gene_id: ENSG00000188536 gene_name: HBA2 gene_location: 16:172876-173710 ---
Set                            LogFC                    LogCPM                  P-val
CTR148_FC_Neurons              -3.32939138352444        9.2573724677293         2.98498695727589e-07
--- gene_id: ENSG00000131018 gene_name: SYNE1 gene_location: 6:152121687-152637801 ---
Set                            LogFC                    LogCPM                  P-val
IB5508_OC_pericytes            -2.03184102965733        12.6064877200671        0.00859384122119039
--- gene_id: ENSG00000250305 gene_name: TRMT9B gene_location: 8:12945642-13031503 ---
Set                            LogFC                    LogCPM                  P-val
IB5431_CB_Fibroblasts          -3.30604255457195        13.841668958432         0.00884184393501855
--- gene_id: ENSG00000237298 gene_name: TTN-AS1 gene_location: 2:178521183-178779963 ---
Set                            LogFC                    LogCPM                  P-val
CTR018_FC_Astrocytes1          -0.316615996533461       10.2879181257779        0.00954541444361825
--- gene_id: ENSG00000198804 gene_name: MT-CO1 gene_location: MT:5904-7445 ---
Set                            LogFC                    LogCPM                  P-val
IB6125_OTC_pericytes           3.26610602792966         13.2944328655107        0.00816665751367035
--- gene_id: ENSG00000144040 gene_name: SFXN5 gene_location: 2:72942036-73075619 ---
Set                            LogFC                    LogCPM                  P-val
IB6001_OTC_pericytes           2.90195736218817         11.7471472497845        0.000767368076280855
--- gene_id: ENSG00000000003 gene_name: TSPAN6 gene_location: X:100627108-100639991 ---
Set                            LogFC                    LogCPM                  P-val
IB5869_OTC_astrocytes          -0.660450631645688       10.4728881049839        1.78383112098662e-05
--- gene_id: ENSG00000124253 gene_name: PCK1 gene_location: 20:57561080-57568121 ---
Set                            LogFC                    LogCPM                  P-val
```

CTR018_FC_Endothelial -3.26512193064519 11.9159251646594 8.18652086329288e-05
--- gene_id: ENSG00000123560 gene_name: PLP1 gene_location: X:103773718-103792619 ---

| Set | LogFC | LogCPM | P-val |
|---|---|---|---|
| FTD014_TC_OPCs | -6.07101084686221 | 11.5729598699431 | 8.87385703135769e-07 |

--- gene_id: ENSG00000198712 gene_name: MT-CO2 gene_location: MT:7586-8269 ---

| Set | LogFC | LogCPM | P-val |
|---|---|---|---|
| IB6125_OTC_pericytes | 3.40497835670714 | 13.2081631026086 | 0.00816665751367035 |

--- gene_id: ENSG00000118271 gene_name: TTR gene_location: 18:31557009-31598833 ---

| Set | LogFC | LogCPM | P-val |
|---|---|---|---|
| IB5431_CB_Fibroblasts | -7.91886003709671 | 16.0060116595117 | 7.96956405328649e-10 |

--- gene_id: ENSG00000145390 gene_name: USP53 gene_location: 4:119212587-119295518 ---

| Set | LogFC | LogCPM | P-val |
|---|---|---|---|
| FTD083_OC_Fibroblasts | 2.9698009397088 | 13.3155718540069 | 0.00506213944820065 |

--- gene_id: ENSG00000118785 gene_name: SPP1 gene_location: 4:87975667-87983532 ---

| Set | LogFC | LogCPM | P-val |
|---|---|---|---|
| FTD038_OC_CAMs | -2.65058119015326 | 14.4451955106568 | 0.00272942399449416 |

--- gene_id: ENSG00000124766 gene_name: SOX4 gene_location: 6:21593751-21598619 ---

| Set | LogFC | LogCPM | P-val |
|---|---|---|---|
| FTD014_TC_OPCs | -3.95885804228277 | 11.6615178638588 | 0.000386619839959514 |

--- gene_id: ENSG00000244734 gene_name: HBB gene_location: 11:5225464-5229395 ---

| Set | LogFC | LogCPM | P-val |
|---|---|---|---|
| CTR148_FC_Neurons | -4.84021152935336 | 9.87047130306675 | 4.33504921764284e-27 |

--- gene_id: ENSG00000233382 gene_name: NKAPP1 gene_location: X:120178488-120245284 ---

| Set | LogFC | LogCPM | P-val |
|---|---|---|---|
| IB5431_CB_Neurins_In_Purkinje | -0.394552720147285 | 10.4983444005701 | 0.00157487897278553 |

--- gene_id: ENSG00000189223 gene_name: PAX8-AS1 gene_location: 2:113211421-113276581 ---

| Set | LogFC | LogCPM | P-val |
|---|---|---|---|
| IB5431_OC_astrocytes | -0.344287795896894 | 10.4873379032139 | 5.12346602936548e-07 |

--- gene_id: ENSG00000198727 gene_name: MT-CYB gene_location: MT:14747-15887 ---

| Set | LogFC | LogCPM | P-val |
|---|---|---|---|
| IB6125_OTC_pericytes | 3.50432654142455 | 13.22442956605 | 0.00816665751367035 |

--- gene_id: ENSG00000198938 gene_name: MT-CO3 gene_location: MT:9207-9990 ---

| Set | LogFC | LogCPM | P-val |
|---|---|---|---|
| IB6125_OTC_pericytes | 3.29663742603915 | 13.143277562577 | 0.00988850909133735 |

--- gene_id: ENSG00000009694 gene_name: TENM1 gene_location: X:124375903-125204312 ---

| Set | LogFC | LogCPM | P-val |
|---|---|---|---|
| IB5508_CB_Bergmann1 | -0.849000309866203 | 11.1419261698664 | 3.80346581044074e-12 |

--- gene_id: ENSG00000258385 gene_name: ENSG00000258385 gene_location: 14:41296286-41298734 ---

| Set | LogFC | LogCPM | P-val |
|---|---|---|---|
| CTR018_FC_Astrocytes1 | -0.113616876402902 | 10.1802761591301 | 0.000181958062992378 |

--- gene_id: ENSG00000229117 gene_name: RPL41 gene_location: 12:56116590-56117967 ---

| Set | LogFC | LogCPM | P-val |
|---|---|---|---|
| IB5431_CB_Fibroblasts | -3.33384730357473 | 13.8480643789778 | 0.00884184393501855 |

--- gene_id: ENSG00000206172 gene_name: HBA1 gene_location: 16:176680-177522 ---

| Set | LogFC | LogCPM | P-val |
|---|---|---|---|
| CTR148_FC_Neurons | -3.17908324725557 | 9.24024425966097 | 2.21352939356429e-06 |

--- gene_id: ENSG00000198947 gene_name: DMD gene_location: X:31097677-33339609 ---

| Set | LogFC | LogCPM | P-val |
|---|---|---|---|
| CTR148_FC_Astrocytes1 | -0.490838880229006 | 10.2943142649803 | 1.12155760920221e-05 |

--- gene_id: ENSG00000079215 gene_name: SLC1A3 gene_location: 5:36596588-36688334 ---

| Set | LogFC | LogCPM | P-val |
|---|---|---|---|
| CTR018_TC_Oligodendrocytes | -3.2557423620682 | 13.04657071131 | 0.000223567287055581 |

--- gene_id: ENSG00000102287 gene_name: GABRE gene_location: X:151953124-151974680 ---

| Set | LogFC | LogCPM | P-val |
|---|---|---|---|
| CTR018_FC_Astrocytes1 | 0.492480812843149 | 10.2103238095465 | 1.06819758284182e-11 |
